# Comparative evaluation of genomic footprinting algorithms for predicting transcription factor binding sites in single-cell data

**DOI:** 10.1101/2025.08.07.669008

**Authors:** Amanda Everitt, Sean Whalen, Katherine S. Pollard

## Abstract

Transcription factors (TFs) have millions of potential binding sites across the human genome, but only a fraction are bound in a given context. Genomic footprinting aims to identify context-specific binding sites by detecting patterns in open chromatin data. While powerful, these approaches face technical challenges, especially in single-cell applications. We developed a benchmarking framework for cell-type specific footprinting and used it to evaluate the consistency, reproducibility, and equivalency of three leading methods across data quality scenarios and as a function of cell-type similarity. Peak-level read coverage emerged as the strongest predictor of stable footprints. Motivated by limited reproducibility across tools, we built an ensemble model that improved concordance with ChIP-seq. To encourage broader adoption and continued tool development, we provide practical guidelines for robust genomic footprinting in single-cell datasets and a roadmap for extracting deeper insights about how gene regulatory networks vary across cell types in complex tissues.

## INTRODUCTION

Single-cell experiments highlight the complexities and subtleties of gene regulation, especially the crucial role transcription factors (TFs) play in shaping cell fate trajectories and disease progression^1–4^. TFs are regulatory proteins that bind DNA to impact gene expression. However, determining their active binding sites is not straightforward. TFs operate within intricate context-dependent networks^5,6^, behave differently in combination^7^, and exhibit varying degrees of sequence specificity^8–10^ and biophysical interactions^11,12^. These factors complicate efforts to measure TF functionality and underscore the value of single-cell datasets, where context-dependent behaviors can be more effectively disentangled.

Genomic footprinting is an approach for genome-wide prediction of active TF binding sites (TFBS) for multiple TFs simultaneously. This method capitalizes on the observation that bound TFs leave a distinctive pattern (footprint) in experiments which measure open chromatin regions like Assays for Transposase-Accessible Chromatin (ATAC-seq) and DNase-I hypersensitive sites (DNase-seq). This pattern resembles a ‘peak-dip-peak’ shape, where the dip corresponds to a ∼6-20bp region where the DNA-binding protein outcompetes the cleavage enzyme (i.e. DNase or Tn5) and shields the DNA being cut. Given the reference genome sequence at the footprint region, sites can be mapped to probable TFs based on position weight matrices (PWM) representing their sequence preferences.

Genomic footprinting has evolved over decades^13–21^, but the wide range of methodological choices and underlying assumptions continue to complicate direct comparisons. Most tools take aligned reads, peak regions, and PWMs as input and output footprint genomic coordinates, scores, and PWM matches. However, the path from input to output varies significantly along four key dimensions: *de novo* versus motif-centric algorithms, supervised versus unsupervised training, bias correction method, and inferential versus descriptive statistics. *De novo* algorithms detect footprints before PWM matching, while motif-centric algorithms identify motif positions first and then assess footprint presence. Many tools use chromatin immunoprecipitation (ChIP-seq) data as ground truth for supervised training, whereas others independently learn patterns. Like many sequencing-based measurements^22^, ATAC-seq and DNase-seq are influenced by non-random cut preferences of cleavage enzymes, and tools adopt varying strategies to model this bias. Finally, some methods derive scores from null distributions while others directly summarize read density.

While genomic footprinting has provided valuable and novel insights, its application remains challenging and has faced considerable criticism^23^. Notably, uncertainty about the required read depth^24,25^ and limited bioinformatic guidelines hinder widespread application. These challenges are further amplified in single-cell data, where read coverage at individual TFBSs may be too sparse to infer occupancy, even after aggregating reads across similar cells (pseudobulking), and many analytical decisions are required. Pseudobulking based on cell type or condition can make single-cell ATAC-seq footprinting sufficiently accurate to examine genome-wide trends^26^, but unique challenges arise from reduced data quality and distributional properties unlike those from bulk-sequencing. For instance, a homogenous cell cluster may reach a high signal-to-noise ratio with fewer reads, whereas a heterogeneous cluster would require more reads to capture accurate footprint patterns, even if individual sites are averaged genome-wide per TF.

Existing footprinting tools have not yet been rigorously and independently benchmarked across diverse single-cell conditions. This is partly due to the computational and conceptual challenge of generating large numbers of realistic benchmarking datasets comprising read alignments, not simply count matrices.

To address this gap, we developed a rigorous read-level downsampling pipeline, scBAMpler, and used it to benchmark three leading footprinting methods – PRINT^26^, TOBIAS^15^, and HINT^14^– on their ability to measure cell type-specific TF binding in single-cell open chromatin data. Rather than evaluate design choices, we aimed to provide a neutral, practical framework to guide users and developers in improving the consistency and reproducibility of single-cell footprinting. Specifically, we evaluated footprint consistency under varying data quality, reproducibility across tools, equivalency to ChIP-seq, and TF-specific biases and trends. Our results will help users design single-cell experiments, optimize pseudobulking strategies, and perform meaningful cross-sample comparisons of TFBSs.

## RESULTS

Our benchmark centered on four computational components: scBAMpler, tool selection, dataset generation, and performance evaluations. We first describe each component, followed by in-depth analyses of key findings, which we integrate into a final recommendation section (Figure 1A).

**Figure 1.**
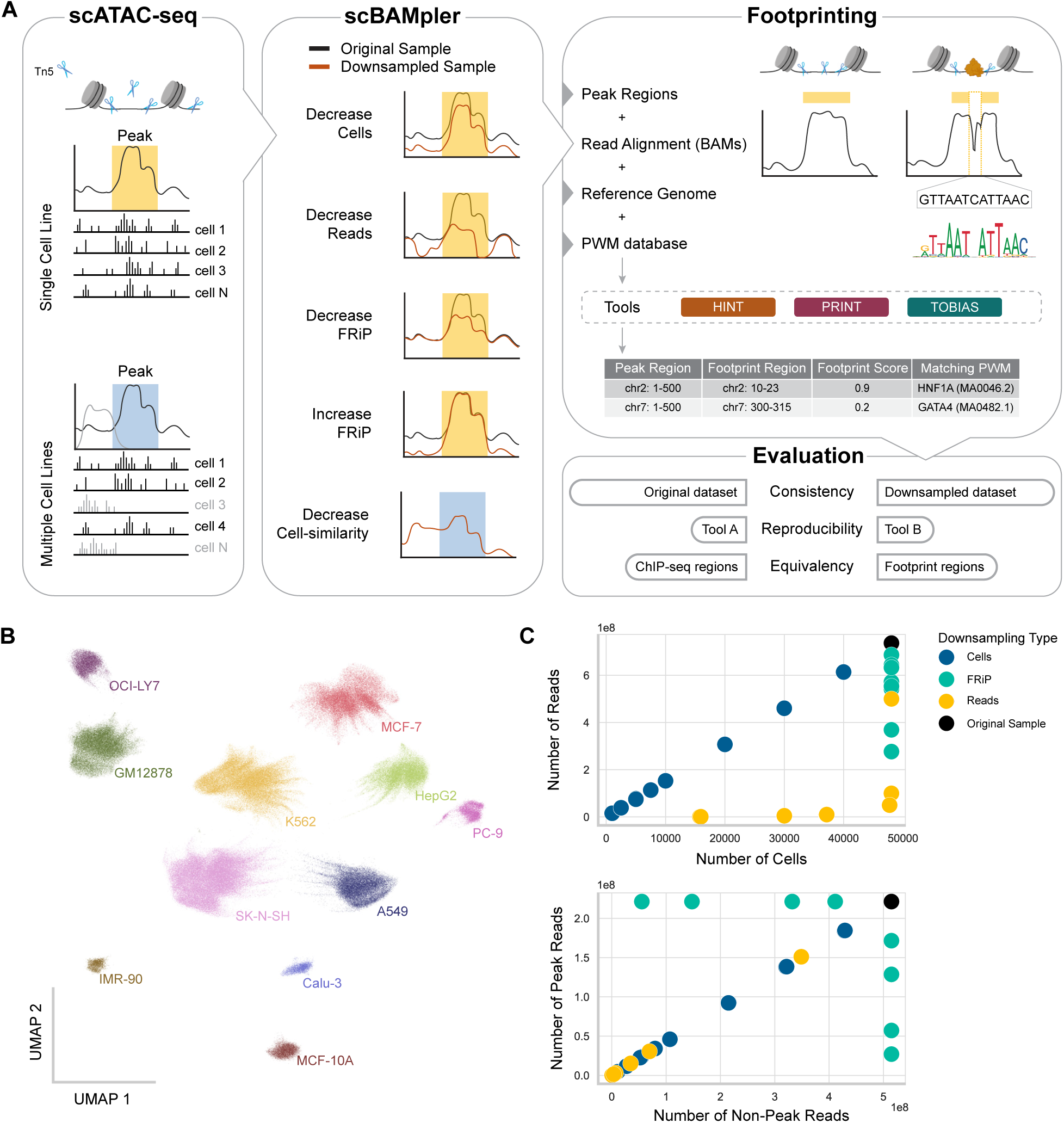
Overview of Benchmarking Study Design. **(A)** Single-cell ATAC-seq captures Tn5 cut sites in open chromatin, generating read pileups indicating accessible regions (peaks). In homogeneous populations (e.g. single cell lines), cells produce a shared signal, while heterogeneous populations (e.g. mixed cell lines) show distinct profiles. Our tool, scBAMpler, strategically downsamples alignments—by reducing cells, reads, or FRiP—to simulate varying signal quality. Multiple cell-types can be used to create samples with varying cell-cell similarity as well. The resulting read alignment files (BAMs) can be input to any footprinting tool to detect local depletions in Tn5 cuts, indicating TF binding. The sequence underlying the depleted region is mapped to known motifs via PWMs, ultimately returning footprint coordinates, scores, and TF matches. We evaluate these outputs for consistency, reproducibility, and equivalency across sampling strategies. **(B)** UMAP of the ENCODE snATAC-dataset, colored and labeled by cell line (n=256,150). **(C)** Scatterplot showing downsampling effects in K562. Points represent downsampled datasets, colored by the variable being evaluated. Downsampling reads (yellow) maintained cell counts until extreme lows, with constant FRiP. Downsampling cells (blue) showed a linear reads-to-cell relationship with constant FRiP. Altering FRiP (emerald) kept cell counts constant with minor read count variations. Although convenient, referencing datasets by cell count alone can be misleading; for instance, 50,000-cell samples can range from 600M to 100M reads depending on how other parameters are tuned.

### Establishing a read-level downsampling framework: scBAMpler

To systematically assess how experimental data quality affects footprinting, we sought to generate samples that isolate individual factors. Because footprinting relies on read-level data (e.g. BAM files), existing platforms that generate count matrices were not applicable^27,28^. We considered two platforms which produce read-level outputs^29,30^, but they either modeled read distributions from bulk ATAC data or introduced synthetic reads that could bias results. Therefore, we developed a new read-level downsampling framework, scBAMpler, that can alter a single-cell (sc) or single-nucleus (sn) chromatin accessibility dataset’s read count, cell count, fraction of reads in peaks (FRiP), and cell-to-cell homogeneity while preserving the original cell attributes. While these metrics are interrelated, each captures a distinct aspect of data quality and provides a different entry point for users. Because scBAMpler is flexible and general, we expect it will be useful for applications beyond this study.

### Tool selection and key methodological considerations

Our objective was to delineate general limitations of footprinting from shortcomings specific to individual algorithms. We selected three widely used tools that include Tn5 bias correction and have maintained code bases: HINT^14^, PRINT^26^, and TOBIAS^15^. Our benchmark is designed to allow future methods meeting this criteria to be incorporated and evaluated as released.

Key methodological distinctions impact interpretations both within- and across-tools (Table S1). First, HINT identifies footprint regions with or without a PWM match, while TOBIAS is strictly motif-centric. PRINT is motif-centric when PWMs are provided, but de novo when they are not. Since most users seek TF-level resolution, we evaluate PRINT in its motif-centric mode unless otherwise noted. Second, HINT uses nucleosome-free reads (fragment length (0,145] bp), whereas TOBIAS and PRINT use insertion sites (i.e. the location where Tn5 integrates sequencing adapters). Third, the meaning of the footprint score varies. In HINT, it reflects read depth; in PRINT, it indicates local depletions of insertion sites; and in TOBIAS, it combines depletion with general accessibility. Finally, some tools require a threshold to classify sites as bound or unbound. For TOBIAS, we used the internally calculated ‘bound threshold’; for PRINT, we used 0.3 (approximately the 65th percentile) and a scale of 30, which broadly captures footprints while excluding most nucleosomes. Although numerous tool-, TF-, and context-specific optimizations are possible, fine-tuning is beyond the scope of this benchmark, which aims to evaluate footprinting performance using typical settings.

### Comprehensive benchmark dataset captures pseudobulk characteristics

Since footprinting inherently lacks a ground truth, we constructed a large, well-characterized snATAC-seq dataset by jointly processing 11 ENCODE^31^ cell lines (Figure 1B). We selected five lines (HepG2, K562, MCF-7, GM12878, SK-N-SH) to define distinct cell type clusters that are stable, clearly understood, and can be evaluated by external evidence. For each line, we then used scBAMpler to generate 22 conditions varying in cell counts, read counts, and FRiP – all in triplicate to control for random sampling effects. Finally, we used scBAMpler to downsample the full dataset of 11 cell type clusters, generating samples with varying levels of intra-cluster similarity. Our method reliably creates samples with the specified properties (Figure 1C), and the resulting peak calls align with expected signal-to-noise shifts (Figure S1A). Through this framework we constructed the largest footprinting benchmark dataset to date, comprising nearly 400 BAM files representing diverse conditions, and the first to emphasize cell-cell homogeneity and signal-to-noise ratios rather than random read downsampling.

### Standardized evaluation of tools using two ground truths

Following the creation of read-level datasets, we implemented a secondary pipeline to run each footprinting tool and standardize its output. Tools were given identical inputs to isolate BAM file reads as the only source of variation. Footprint regions were called for every downsampled BAM file and for the full-depth pseudobulk sample prior to downsampling. In our primary performance evaluations, this full-depth “original” condition served as a best-case reference for computing precision, recall and their harmonic mean (F1 score) for each downsampled condition. This comprehensive, controlled dataset provides a robust foundation for systematically evaluating both within-tool consistency in response to data quality variations and across-tool differences in detection.

As a second strategy, we evaluated each footprinting algorithm by how well it replicated ChIP-seq, which maps protein-bound DNA using TF-specific antibodies. While widely used and informative, ChIP-seq is an imperfect gold standard^23,32,33^ due to experimental factors like indirect binding, antibody specificity, and crosslinking efficiency. Even replicated experiments for the same TF, cell line, and antibody under basal conditions can vary substantially^34^. Beyond experimental confounders, footprinting accuracy is inherently constrained by how well ATAC signal and PWMs approximate ChIP signals — assumptions often built into prior tool comparisons. To decouple these factors, we first evaluated ChIP-seq concordance with motif matches and ATAC-seq signal. We then used downsampled datasets to evaluate how agreement between ChIP and footprints decline with read quality, exploring differences across tools, TFs, and score thresholds. By using ENCODE cell lines, ChIP-seq data was available for 1,064 TF-cell line pairs.

### Dependence of footprint calls on peak read coverage

To understand fluctuations of individual footprint regions, we calculated how often regions were retained, lost, or incorrectly gained in downsampled conditions versus the original, non-downsampled, dataset. We report performance as F1 scores calculated per cell line across all peaks and JASPAR^35^ PWMs. Metrics were similar across condition triplicates (mean Pearson r=0.97 ± 0.08, Figure S2A), so we report average performance throughout.

As expected, all tools struggled to recover original footprints as data quality decreased (Figures 2A and S2B). HINT and PRINT showed gradual and consistent decreases as a function of each downsampling parameter, with the most pronounced losses occurring below 5,000 cells and 1e7 read pairs. This trend was consistent across cell lines. TOBIAS maintained high F1 scores as data quality initially declined but dropped sharply at lower quality levels.

**Figure 2.**
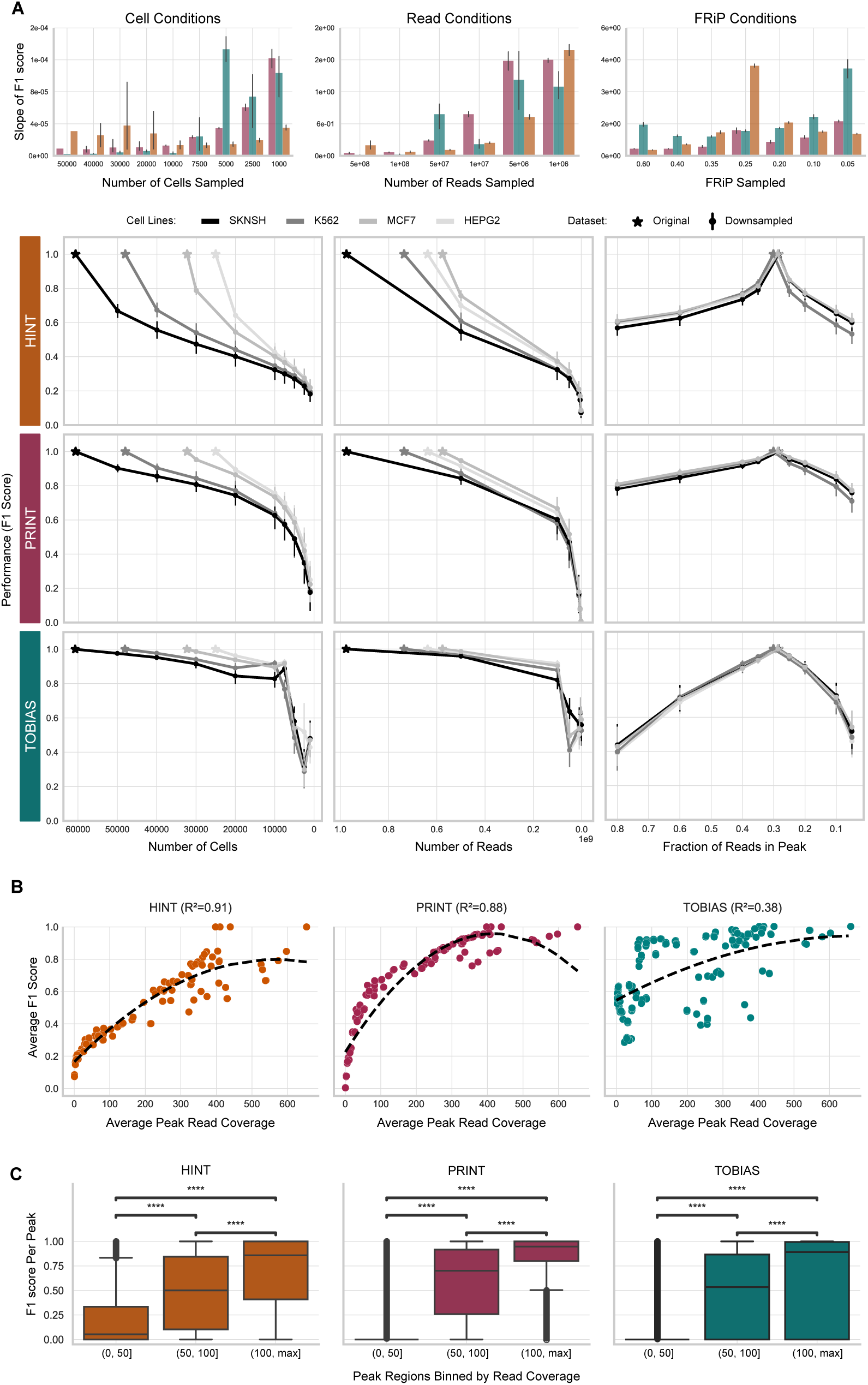
Downsampling Framework Identifies Peak Read Coverage as Key Predictor. **(A)** Line plots show performance changes (F1 score) across downsampling conditions. Stars indicate the starting values from original datasets. Each dot represents a downsampled datasets mean ± SD across PWMs. Above, a grouped bar chart shows the F1 score slopes at each discretized condition with error bars indicating the SD across cell lines. This helps highlight shared points of inflection. Because plots represent changes relative to original data (stars), deviations in either direction are expected. For example, increased FRiP may produce more conservative footprint sets. The star position is not biologically a ground truth, but rather a consistent point of reference. **(B)** Scatterplot illustrates the relationship between a sample’s average read depth in peak regions (xaxis) and F1 score (y-axis). Each point is a downsampled dataset. The dashed black line shows a polynomial regression; R² values are shown in parentheses. Among all metrics tested (FRiP, cell number, peak/non-peak read pairs, total read pairs, average peak coverage), average peak coverage had the strongest univariate correlation and most consistent performance decline (See also Figure S2C). **(C)** F1 scores were calculated per peak and grouped by read coverage. Regions with <50 and 50– 100 reads show significantly lower F1 scores across tools. Pairwise t-test significance indicated by asterisks (**** = p < 1e-4).

Intrigued by the varying dependence of the three methods on read support, we jointly modeled the F1 scores of all samples—across different downsampling conditions and cell lines—as a function of data quality metrics. We found average peak-read coverage was the strongest predictor and the only metric to exhibit a synchronized decline (Figures 2B and S2C). TOBIAS was least dependent on peak-read coverage alone, but its predictability improved substantially with the inclusion of sample FRiP (R²=0.38 to 0.74), reflecting TOBIAS’s use of relative signal in score calculations. Finally, we calculated the F1 scores per ATAC-peak and found those with fewer than 100 reads had significantly lower scores (Figure 2C). This suggests that applying a universal filtering threshold of 100 reads per peak can improve the consistency of footprint calls—which, when implemented, did improve performance (Figure S2D).

Together, these findings suggest that performance declines were driven by false negatives in regions with fewer than 100 reads, suggesting a potentially important filtering threshold—particularly when interpreting the absence of a footprint in one cell population. Additionally, although TOBIAS produces more stable results, it appears least affected by read support — a potentially undesirable trait when the goal is to detect context-specific TFBSs — whereas HINT and PRINT behave more as expected, showing improved performance with higher peak-read levels.

### Performance patterns highlight methodological consequences

Many of the observed performance trends stem from fundamental methodological differences between tools. A common question is whether de novo tools require higher read depth than motif-centric tools due to their broader search space and increased risk of false positives^25,36^. As motif-centric tools, TOBIAS and PRINT consider the same motif sites across all downsampling conditions, primarily gaining false negatives as read support declines (Figures S2E-G). In contrast, HINT, which lacks predefined sites, gains both false negatives and positives. This is reflected in its improved performance in a PWM-agnostic F1 score calculation, unlike the other tools (Figure S2H). Although de novo methods like HINT may appear less consistent, our analysis shows their footprint regions are largely stable, with small coordinate shifts affecting PWM match scores and creating a misleading impression of instability.

We also investigated the jagged performance drops observed in TOBIAS and found that large shifts in its internally estimated ‘bound threshold’ were responsible (Figure S2I). This threshold is designed to adapt footprinting to each sample’s properties but becomes less reliable at pseudobulk-level inputs (e.g. <10,000 cells). GM12878 was especially affected by this instability (Figure S2G). When reusing the original sample’s threshold across conditions, performance stabilized and resembled other tools (Figure S2J). This underscores how differences in sample quality can distort analyses when key parameters are estimated from low quality data.

Overall, footprinting consistency depends on both data-quality and tool-specific behaviors. For example, TOBIAS is particularly sensitive to FRiP due to its score calculation and its bound threshold can have significant implications. Achieving consistent footprinting performance (F1 > 0.6) required around 100 million read pairs per cell population. In our dataset (20000 read-pairs/cell, 0.29 pseudobulk FRiP), this was roughly 6,000 cells, though other datasets will return different cell-level estimates (Figure S2K). This recommendation reflects TF-specific results for PRINT and TOBIAS, and TF-agnostic results for HINT.

### Cell diversity impacts footprint consistency

Having established peak-read coverage as the key sequencing factor, we next examined how cell-cell homogeneity affects footprinting consistency. Homogeneity within each pseudobulk depends on both the epigenetic variability in a complex tissue and the clustering decisions made during analysis. Generally, clustering and pseudobulking resemble wave interference: merging similar read signals enhances pattern detection, while dissimilar signals obscure it. Although clustering promotes constructive interference, footprinting operates at much lower resolution than high-dimensional methods like UMAP, making optimal clustering resolutions less obvious. This tradeoff between signal clarity and read depth—sharper signals in small clusters versus deeper but noisier signals in large clusters—is especially relevant for continuous-state systems like development or disease.

We aimed to identify where pseudobulking shifts from enhancing to impairing performance, and how read depth influences this tradeoff—ultimately to inform optimal clustering strategies. As a reference, we defined a ground truth set of footprint regions from each cell line aggregated into a pseudobulk. To evaluate similarity between synthetic populations and this reference, we computed a distance metric based on the peak-by-cell count matrix that is comparable across experiments and avoids dependence on dimensionality reduction parameters.

To isolate the effect of this similarity metric on footprinting, we generated nearly 300,000 synthetic pseudobulk populations with varying cell-line mixtures and peak read depths that span real dataset properties while minimizing confounders (Figures S3A-C). From these, we selected 80 samples for footprinting—20 per read-depth condition—differing only in their distance to the reference. Footprinting results were then compared to the reference pseudobulk using a union peak set to capture footprints from any contributing cell line.

When visualizing the F1 score as a function of decreasing cell similarity, all tools show distinct negative slopes depending on the starting peak read depth (Figure 3A). This difference largely disappears when focusing only on peak regions with >100 reads, highlighting that peak filtering can largely mitigate false negatives from lost footprint regions. Although overall trajectories appear similar, PRINT consistently shows higher precision, while TOBIAS achieves greater recall—reflecting differing dependencies on read depth and methodologies (Figure S3D). As expected, the number of footprints for cell line–specific TFs^37^ varied with cell population diversity (Figure 3B). Together, these findings show that footprinting is sensitive to cell type heterogeneity and suggest that, because of its lower granularity compared to high-dimensional embeddings, pseudobulking cells based on peak-count similarity may be more effective.

**Figure 3.**
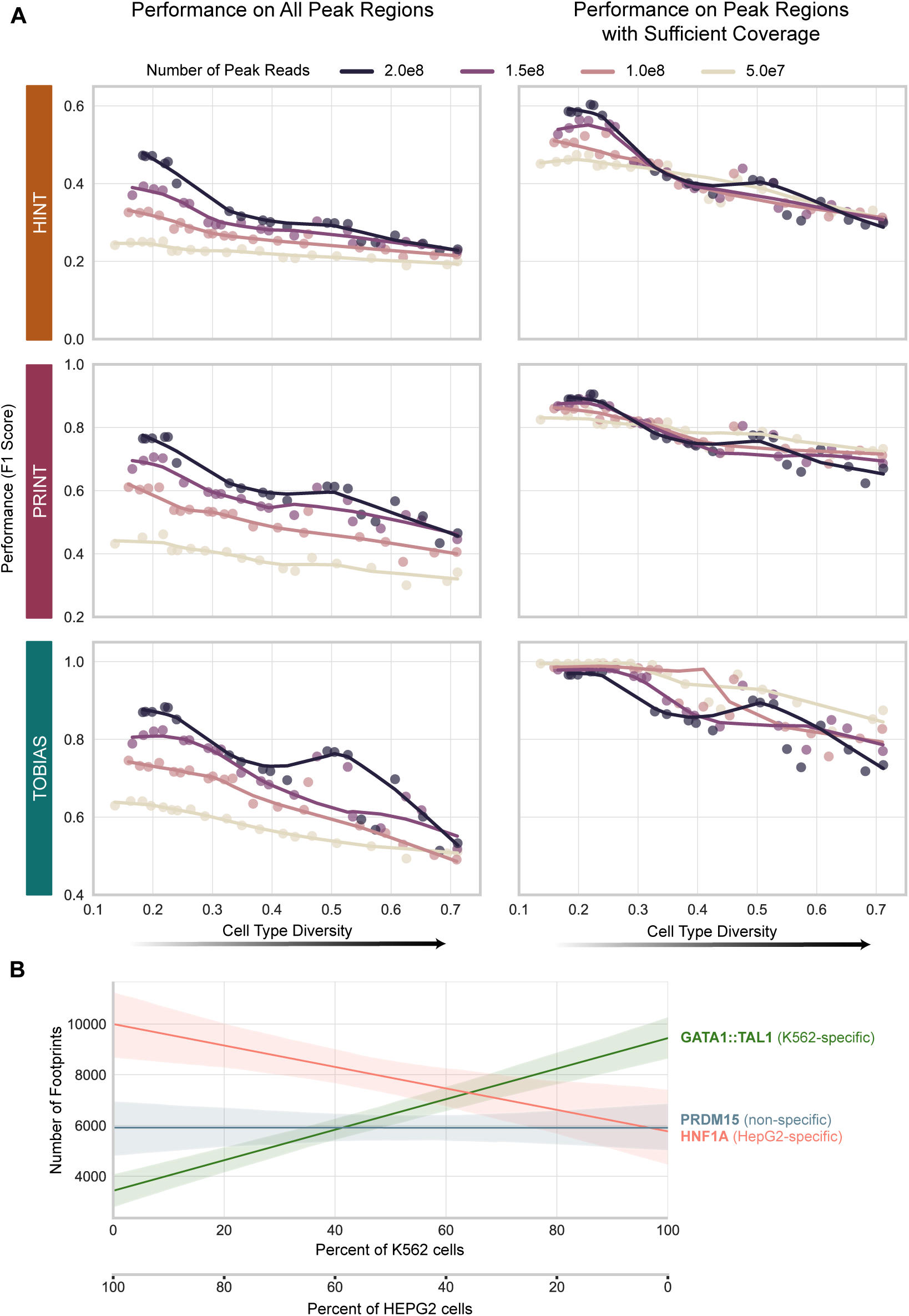
Pseudobulk Heterogeneity Obscures Footprinting Results. **A)** Each plot shows how performance (F1 score) changes with increasing cell type diversity across 80 hypothetical cell populations. Each point represents the average F1 score across 350 representative motifs for given dataset, colored by peak read-depth category. The x-axis is the average distance to the reference cell line centroid (see Methods). LOESS curves fit within each category help to visualize overall trends. To enable consistent comparisons, TOBIAS’s bound threshold was fixed at 0.025. Left column plots use all union peaks; right column plots include only peaks with >100 reads in the reference, which reduces false negatives and boosts performance. **B)** For 34 samples composed exclusively of K562 or HepG2 cells, we performed linear regression to model footprints counts as a function of pseudobulk composition. As K562 proportion increases, footprint counts for an associated TF GATA1:TAL1) rise, while those for a HepG2-associated TF HNF1A) decline. A TF not associated with either (PRDM15) shows no significant change. Shaded areas represent the 95% confidence intervals.

### Methods exhibit limited reproducibility

Since methodological choices affected consistency within each tool, we next assessed how much footprinting results agreed across tools, and whether this agreement changed with downsampling.

To establish a baseline, we first calculated the number of shared footprint regions across the original samples by counting footprints per open-chromatin region (OCR). Strikingly only 7% of footprint regions were shared across the three tools on average, dropping to 3% when requiring a matching PWM (Figure 4A). Optimizing score thresholds yielded minimal improvement (Figures S4A-C), and genome-wide metrics were consistent across tools (Figure S4D), suggesting that low concordance stems from more fundamental differences. In de novo tools, small coordinate shifts can lead to slightly different motif matches—often switching between closely related PWM family members. For motif-centric tools, the discrepancies trace back to the initial candidate TFBS identification, where 64% of sites were unique to TOBIAS (Figure 4B). Because PWM matching is handled internally, users typically cannot control the algorithm or input coordinates. While we were able to standardize the p-value threshold, this did not improve agreement, again pointing to deeper algorithmic differences. Without precise control over this step, it is difficult to separate the impact of TFBS selection from footprinting itself. This substantial difference between tools underscores how subtle methodological choices affect reproducibility in ways that are often invisible to users.

**Figure 4.**
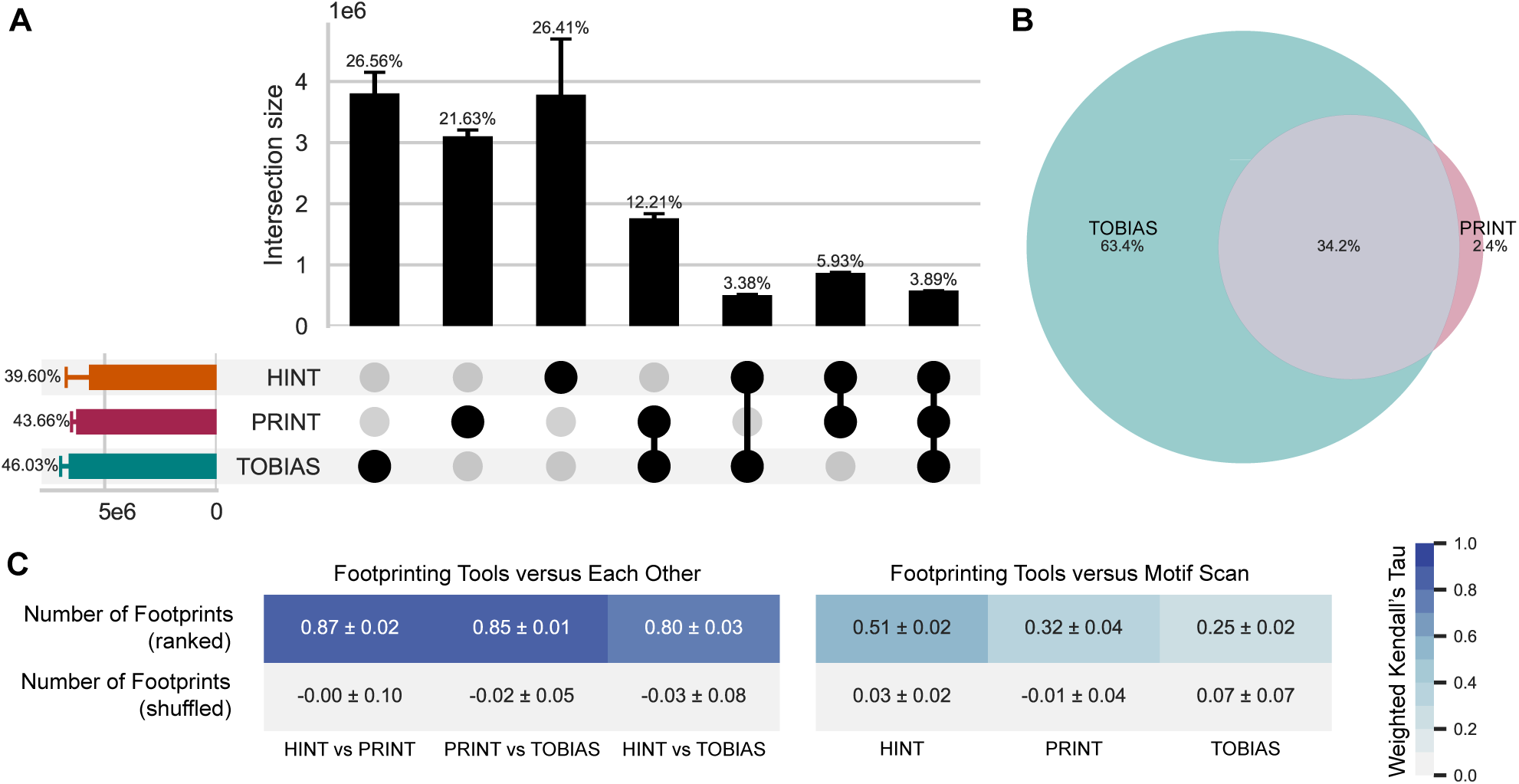
Local Footprint Calls Are Distinct Across Tools. **(A)** We counted how many footprints were called per OCR for each tool and compared how closely those values aligned across tools. When requiring the same PWM in the same OCR, only 3.89% of regions are shared by all three tools (column 7), while ∼20% are unique to one tool (columns 1–3). Vertical bars represent pairwise intersection percentages (mean ± SD across cell lines). Horizontal bars show each tool’s contribution to the union set, indicating roughly equal representation. By not requiring overlapping footprint coordinates within the OCR, this comparison offers a more generous view of tool agreement. **(B)** Motif-centric tools show limited overlap in candidate TFBSs. Prior to footprinting, tools define regions where footprints will be evaluated; TOBIAS (green) and PRINT (pink) shared 34.2% of calls on average across cell lines. **(C)** Despite low locus-level overlap, tools similarly rank PWMs by total footprints counts. Left, each pairwise tool comparison are shown, with the weighted Kendall’s tau (mean ± SD) displayed as text and tile color. For reference, metrics calculated on shuffled ranks are included. Right, tools do not rank PWMs similarly to a motif scan, which identifies all possible TFBSs in peaks.

We reasoned that even if tools identify largely unique footprint locations, they may still detect a similar set of PWMs and rank them comparably. To test this, we counted footprints per PWM, ranked them, and computed a weighted Kendall’s tau which gives more weight to frequent TFs. We found strong concordance across tools in the original samples and next analyzed downsampling conditions (cells, FRiP, reads), again observing high agreement across cell lines and tools (Figure 4C). PRINT and HINT were the most concordant. Reassuringly, PWM rankings were more consistent across footprinting tools than with rankings based on motif scans (i.e. the number of potential TFBS in peaks) or motif enrichment scores (Figure S4E), highlighting that footprinting tools capture a signal distinct from related approaches.

Our results indicate that footprinting tools capture largely unique signals, likely stemming from factors beyond user control, such as initial read processing, Tn5 bias correction, and PWM-match algorithm. Despite regional discrepancies, we observed global concordance in overall metrics and PWM rankings. Collectively, these findings suggest that the tools may be too methodologically distinct to align directly and that ensemble approaches leveraging information across multiple tools may offer a better path forward.

### Transcription factors with high GC content motifs result in robust footprinting

Since tools ranked PWMs similarly, we sought to understand which TF motifs were most frequently identified and what factors contributed. Previous studies showed PWM characteristics – GC content, information content, length, genome-wide prevalence – affect footprinting results^17^. However, we also expect TFs to produce stronger footprint signals in cell types where they are biologically active.

To obtain a robust measure of PWM consistency, we normalized and averaged PWM performance across all scBAMpler downsampling conditions and cell lines. We then correlated this averaged footprinting signal with PWM properties. GC-rich motifs were the most likely to exhibit—and retain—higher footprint counts (Figure 5A). Members of the SP/Kruppel-like and E2F family were among the most detected, while ARID and HD-CUT ranked among the least. No consistent patterns were observed for PWM length, information content, peak prevalence, or RNA expression levels (Figures S5A-B). This indicates that ranking PWMs by footprint count alone favors GC-rich motifs and broadly active TFs including universal stripe factors^11^ (USFs), while not necessarily enriching for expressed TFs.

**Figure 5.**
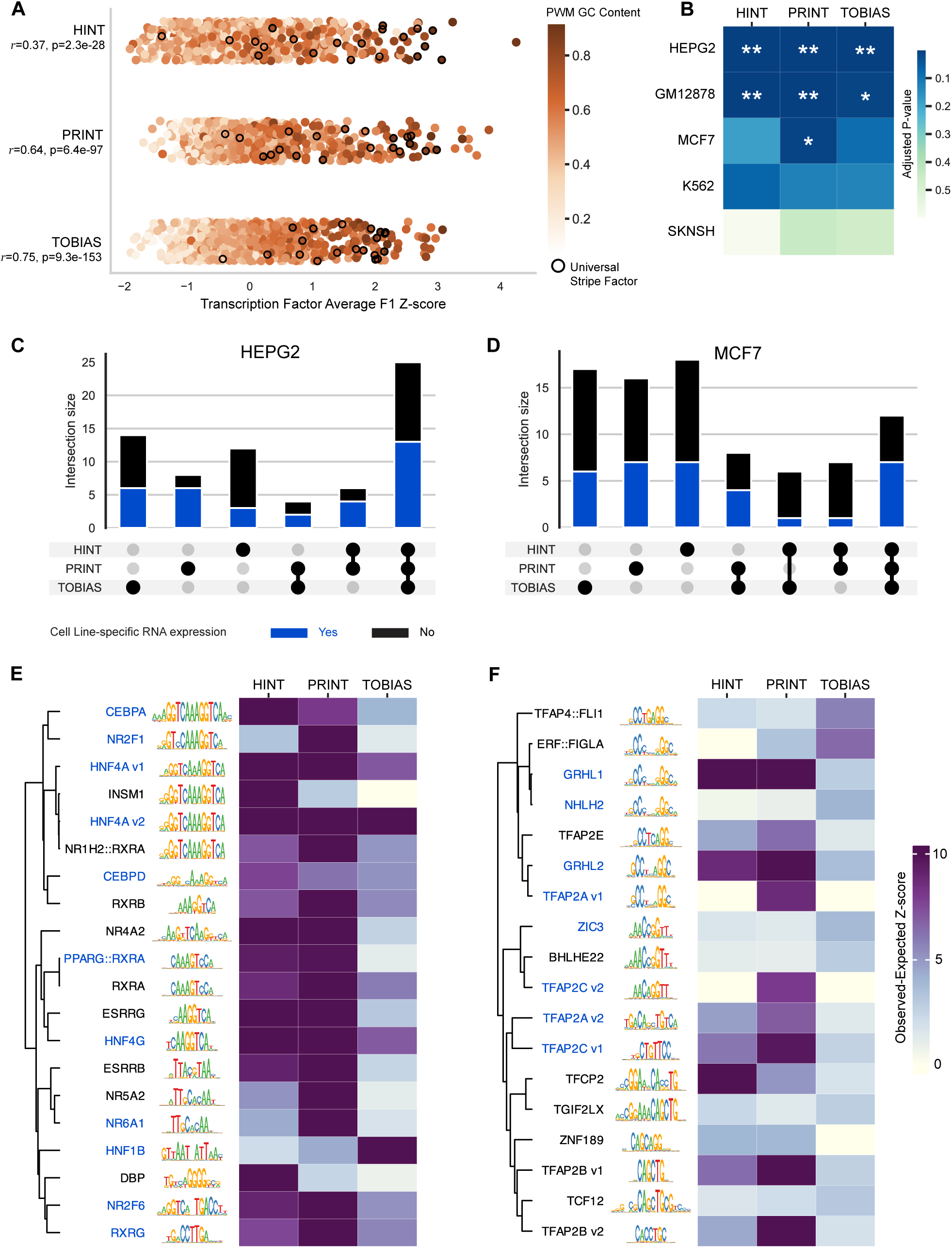
All Tools Highlight TFs with Cell Line–Specific Expression. **(A)** Strip plot reveals a positive relationship between PWM GC content and average footprint signal in downsampling scenarios. Color indicates GC content; each dot represents a JASPAR PWM. Pearson correlation and associated p-value shown. Universal stripe factors, which are commonly active in cells, are outlined in black. **(B)** Permutation tests evaluated whether TFs with cell line–specific RNA expression (CLE-TFs) had higher normalized footprint counts than expected by chance. Heatmaps display Benjamini-Hochberg adjusted p-values across cell lines (rows) and tools (columns). Statistical significance is indicated by stars (** = p < 0.001; * = p < 0.05). CLE-TFs are defined as the top 20% of genes with elevated RNA expression relative to the mean across other cell lines (See Methods). **(C** and **D)** All tools reliably identify CLE-TFs (blue), when comparing a TF’s normalized footprint count in one cell line to its average across others. The overlap of TFs ranked in the top 5% by normalized footprint counts in at least one tool (n = 43) is shown in the UpSet plot. Bar chart colors indicate CLE-TFs in each intersection, for both (C) HepG2, a liver-derived cell line, and (D) MCF-7, a breast cancer cell line. **(E** and **F)** Heatmaps show varied detection strength for CLE-TFs. The top 15 TFs by normalized footprint counts in any tool are shown. Rows represent TFs, labeled with their PWM sequence logos, and ordered by hierarchical clustering (complete linkage of Euclidean distances between PWM matrices). Tools perform similarly in (E) HepG2, whereas PRINT shows uniquely strong performance in (F) MCF-7. Multiple PWMs for the same TF are indicated by version numbers. Since closely related PWMs can produce nearly identical hits, treating them independently may underestimate detection performance.

While GC-rich motif enrichment may reflect footprinting bias, it could also result from the elevated GC content in OCRs^38^. To distinguish this, we generated synthetic PWMs that occur in the genome but do not correspond to known TFs and hence should not be associated with specific patterns in ATAC-seq data (Figures S5C-D). We repeated footprinting using these synthetic PWMs and tested whether their percent of occupied TFBSs could be predicted from PWM properties alone. Linear models trained on synthetic PWMs explained substantially more variance than those trained or applied to real JASPAR PWMs (Table 1). We estimated ∼20% of the variance in real PWM occupancy could be attributed to PWM properties. Results were consistent across tools and cell lines, although GLIS1 and BCL6B were notable outliers, with predictions consistently overestimating their footprint occupancy. We conclude that real PWMs encode information that footprinting algorithms are leveraging beyond what is captured by basic sequence properties.

**Table 1.**
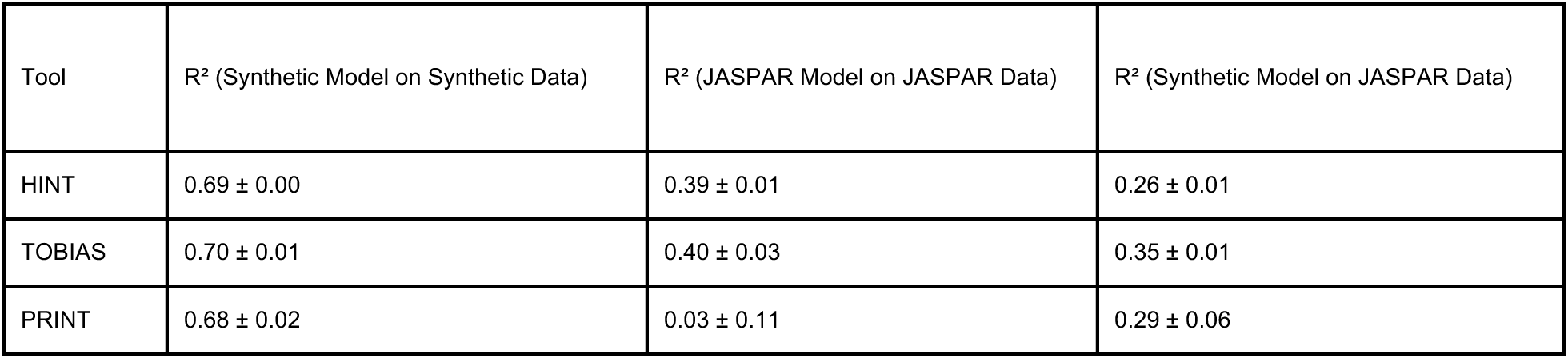
PWM Sequence Features Better Predict Synthetic Than Real Motif Occupancy.

In sum, TFs with GC-rich motifs are more reliably detected by footprinting methods, making them stronger candidates for analysis. This trend is consistent across tools and offers users a straightforward way to assess whether footprinting is appropriate for their TFs of interest. It also highlights that conclusions drawn from single-sample analyses offer limited biological insight, as much of the signal is likely shared across conditions—underscoring the importance of cross-sample comparisons.

### Differential binding modules confounded by sample variability

A primary goal of footprinting is to identify TFs with differential binding across cell types, based on the assumption that active TFs leave stronger footprint signals. However, not all tools support differential analysis, and those that do, implement it differently. To enable a fair comparison, we first applied a simplified observed-versus-expected strategy to identify TFs with cell line–specific RNA expression (CLE-TFs) and then compared the differential binding modules of HINT and TOBIAS.

By comparing each TF’s normalized footprint count in one cell line to its average across others, we reliably identified CLE-TFs (Figures S5E-G). Permutation tests confirmed that all tools significantly prioritized CLE-TFs in HepG2 and GM12878, though the specific TFs and their signal strengths varied substantially (Figures 5B-F and S5H). These results were highly consistent when data quality was matched across samples; repeating the analysis on cell-downsampled datasets recovered ∼80% of the top TFs from 20,000 to 1,000 cells (Figure S5I). In contrast, using the original samples which differed in cell number, produced widely varying TF sets—highlighting that matched data quality mattered more than total cell number.

To evaluate the differential binding modules of HINT and TOBIAS, we first compared replicates from the same cell line with matched data quality (e.g. two samples of 10,000 cells drawn from the same population). Since replicates are highly similar, few motifs should be flagged as differentially bound, providing a proxy for false positives. Both tools yielded low fold-changes but inflated unadjusted p-values, with TOBIAS displaying a noticeable breakpoint around p = 1e-50 (Figure 6A). To our knowledge, neither tool corrects for the number of motifs tested or the number of potential TFBSs per motif, limiting the interpretability of unadjusted p-values across datasets. Therefore, as a reference, we included a HepG2 vs K562 comparison. Selecting motifs within the top 95th percentile, while commonly recommended, only ranks motifs within a dataset and does not convey absolute significance or effect size.

**Figure 6.**
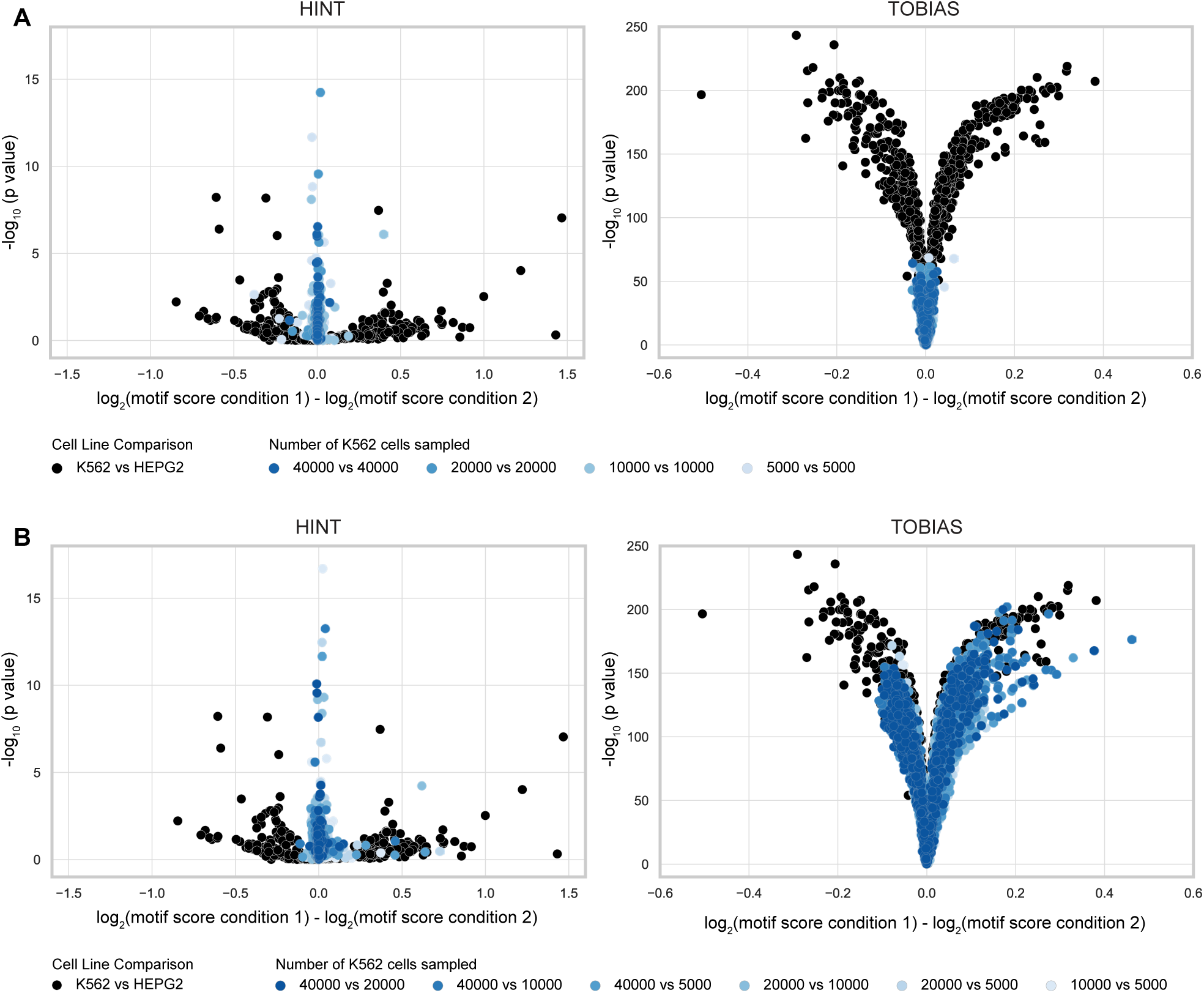
Interpreting Differential Binding Requires a Proper Reference. **(A)** Stacked volcano plots display –log₁₀(p-values) versus fold-changes for motifs. Comparisons between distinct cell lines are shown in black, while technical replicates from the same cell line with matched data quality are shown in blues. Replicates serve as a proxy for areas of false-positives. **(B)** As in Figure 6A, but comparing replicates from the same cell line with unmatched cell numbers, to assess how sample depth affects on fold-changes and p-value inflation.

We hypothesized that p-value inflation would worsen with mismatched data quality. Indeed, when comparing samples from the same cell line but with different cell counts, HINT appropriately minimized fold-changes through depth scaling, whereas TOBIAS produced values as large as those seen between distinct cell lines (Figure 6B). Similar trends emerged when comparing read depth and FRiP (Figures S6A-C). Together, these findings suggest that without carefully matched sample depths, false positives may be indistinguishable from true biological signals in differential footprinting analyses.

### Decoupling tool and method limitations in ChIP-seq baseline

Utilizing ChIP-seq as a gold standard for TF binding, we evaluated three indirect but related components of footprinting pipelines: PWM matches, ATAC-seq signal, and footprint score thresholds.

We intuitively expect the PWM of the profiled TF to be present in its ChIP-seq regions, yet this is often not the case. The best-matching PWM is frequently not the profiled TF but a family member — or even an unrelated TF^39^. In light of this, we first evaluated whether the PWM for each profiled TF could predict its ChIP-seq binding. For many TFs, the matched PWM provided only modest gains over randomized controls (Figures S7A-B), prompting us to restrict our analysis to TFs with reliable PWM recall.

The read depth required for accurate footprinting has been estimated from 50–60^17,21^ million to 100–200 million^13,18,40,41^. We hypothesized that this inflection point reflects the diminishing overlap between ATAC- or DNase-seq and annotated ChIP peaks. Because ATAC-seq signal is diffuse and spans broader regions than ChIP-seq’s targeted peaks, achieving footprint-level resolution at TF-specific ChIP sites would require substantially higher sequencing depth — a challenging task given the assays’ fundamentally different chromatin measurements. To test this, we performed peak calling across downsampled conditions and tracked the overlap with ChIP regions as depth declined. Across TFs we observed a plateau between 100–200 million reads (Figure S7C), supporting this depth cutoff.

Overlap between footprint- and ChIP-regions is commonly used to assess accuracy; however, it is closely tied to the footprint score threshold used to distinguish bound from unbound sites. To evaluate its impact, we used ChIP as ground truth and calculated F1 scores in the original, full-depth samples at each tool’s default threshold and across all score deciles. Optimal thresholds varied widely across tools and cell lines, often substantially outperforming default settings (Figures 7A and S7D). Comparing the performance gain to the required threshold shift revealed that per-TF tuning can significantly boost performance (Figure 7B). For example, TFs with common motifs like SP1 and KLF1 benefitted from stricter thresholds, whereas MAFK required a more lenient threshold in TOBIAS but a stricter one in PRINT. We further dissected tool behavior by deeply tuning performance for two TFs – CTCF and JUND – and found that filtering steps (e.g. requiring motif presence) disproportionately affected TFs (Figure S7E-G). Together, these results emphasize the importance of tuning score thresholds—not only to enable fair tool comparisons, but to account for TF-specific variability in footprinting behavior.

**Figure 7.**
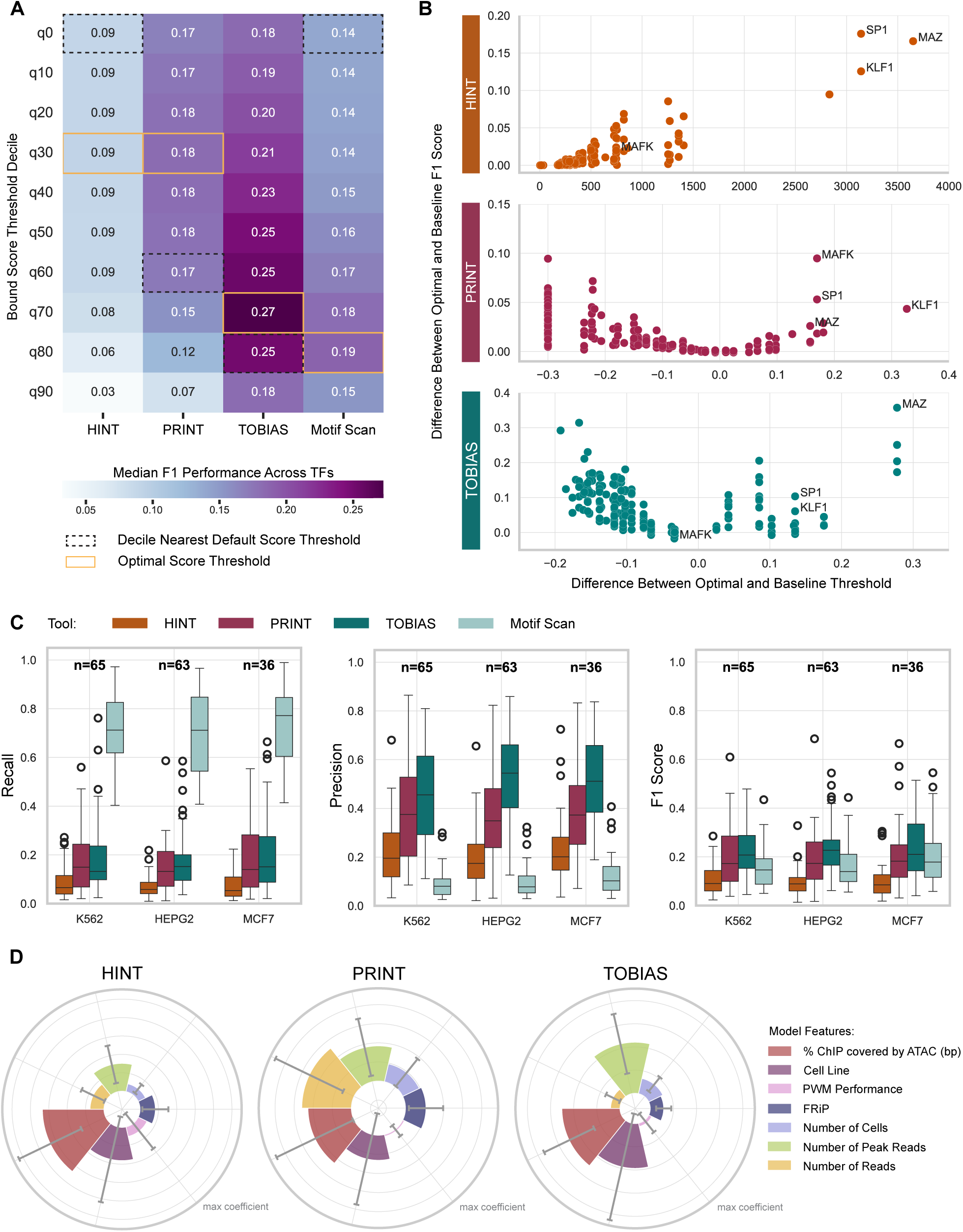
ChIP-seq Equivalency Varies Widely by TF, Tool, and Dataset. **(B)** Heatmap showing median TF performance (F1 score) across score deciles for each footprinting tool. Deciles were computed per tool, and footprints exceeding each threshold were compared to ChIP-seq peaks to assess performance. Rows represent deciles, columns represent tools, and cells are colored and labeled by median F1. The best-performing decile (orange box) often differs from the decile closest to the default threshold (black dashed box), indicating that default settings may be suboptimal. Results are shown for HepG2; similar trends are observed across cell lines (See Figure S7D). **(C)** Scatterplot illustrating the relationship between the improvement in F1 score possible and the change in threshold required. Each point represents a TF-cell line pair, with the x-axis showing the difference between optimal and default score thresholds, and the y-axis the resulting F1 gain. Because HINT lacks a defined default (effectively zero), all threshold adjustments are positive. In contrast, adjustments for TOBIAS and PRINT can be either higher or lower than defaults. **(D)** Box plots of recall, precision, and F1 scores by tool (color), across TFs and cell lines (columns). The number of retained TFs per cell line is indicated as text. Whiskers represent 1.5x the interquartile range. **(E)** Ridge regression identified features predictive of F1 score changes across downsampling conditions, using ChIP-seq as ground truth. A separate model was fit for each tool-TF pair with a fixed alpha. The radar bar plot shows the absolute value of each feature’s coefficient across TFs (mean ± SD). The most consistent predictor was the percentage of ChIP-peak base pairs overlapping ATAC-seq peaks

### Ensemble models improve footprint-ChIP signal equivalency

Using the insights above, we compared footprinting to ChIP-seq by (1) measuring the overlap between ChIP- and static footprint-regions, and (2) assessing how well continuous footprint scores predict ChIP-seq signal.

Using standard score thresholds that reflect typical workflows, all footprinting tools showed lower recall but higher precision than motif scanning, as expected (Figures 7C and S7H). TOBIAS had the highest precision, followed by PRINT—a pattern that persisted under downsampling with only modest performance declines (Figure S7I). TF performance varied by cell line—independent of read depth— explaining previously observed nonlinear declines^13^. To identify features uniquely driving performance changes under downsampling, we trained regression models using sample- and motif-level variables (e.g., cell count, standalone PWM F1 score). Across tools, ChIP-ATAC peak overlap was consistently the strongest predictor, while other features showed tool-specific effects (Figure 7D). Collectively, these findings suggest that ATAC-seq signal quality has a greater impact on footprinting performance than PWM quality or tool choice. Performance declines were mainly due to increased false negatives, reflecting general limitations of ATAC-seq rather than tool shortcomings.

Motif scanning performed reasonably well, consistent with prior reports^17^ (Figure S7J). Both TOBIAS and motif scanning cast wide nets for candidate TFBS: TOBIAS identifies ∼2.5x more sites than PRINT but ultimately filters to similarly sized final sets. While this is influenced by score thresholds, our decile-based analysis supports the broader interpretation. Counterintuitively, methods relying less on ATAC-seq signal performed best—implying that ignoring signal intensity can be advantageous in prediction tasks structured this way. That same trait was a disadvantage in our earlier consistency-based analyses. This has important implications for how footprinting tools are evaluated, particularly in a field where ground truth is limited, uncertain, and often biased.

To assess how well footprint scores predict ChIP-seq signals, we trained machine learning models for two TFs in HepG2: CTCF, a top-performer, and JUND, a moderate-performer. No single tool consistently outperformed the others. For CTCF, PRINT achieved the highest area under the precision-recall curve (auPR) at 0.64, followed by TOBIAS (0.59) and HINT (0.29). For JUND, TOBIAS led with an auPR of 0.28, with PRINT (0.23) and HINT (0.14) trailing closely. Prior analyses showed local differences between tools despite similar global patterns, suggesting that ensemble approaches could improve prediction. To test this, we implemented an ensemble model that integrates scores from all three tools as input features, learning to combine them into a single predictive output. This model captures patterns and relative importance across tools, effectively weighting each score based on its contribution to ChIP-seq signal prediction. All two- and three-tool ensembles outperformed individual models, with the three-tool ensemble reaching an auPR of 0.72 (CTCF) and 0.30 (JUND). These results suggest that integrating footprint scores—each capturing distinct signal aspects or assumptions—can modestly improve the accuracy of ChIP-seq signal prediction.

### Recommendations

We now offer practical recommendations based on our findings. For strong performance (F1 > 0.6), we recommend targeting ∼100 million read pairs per cell population – roughly 6,000 cells, though this varies by dataset. This depth marks an inflection point in both downsampling and ChIP-seq comparisons. With these guidelines, users have several options.

If the goal is to evaluate global differential TF binding across cell populations, closely matching sample quality is essential. While exact matches are not necessary, large discrepancies can cause false positives from detection variability rather than biology. We recommend identifying the cell population with the lowest quality. If it contains >10,000 cells with an average peak-read depth >100, it is well-suited for footprinting and robust detection of cell type-specific differences. Below 10,000 cells, footprint recovery drops sharply—by 20–30% at 5,000 and 5% at 1,000. Users should consider downsampling all populations to match the smallest group or splitting larger ones into pseudoreplicates (e.g. via constrained k-means clustering). We recommend downsampling by cell count (which reduces signal at true peaks) rather than read depth (which introduces artificial variability from new peaks). Because UMAP clustering captures finer resolution than footprinting, merging similar clusters can often boost depth while preserving biological meaning. For differential binding, we advise against using the top 95th percentile as hits. Since PRINT lacks a built-in differential binding module and HINT and TOBIAS underperform in this area, users could define cutoffs using one of three strategies: (1) apply a custom p-value correction; (2) create a background distribution via bootstrapping; or (3) use a reference scale as shown in Fig 6A.

If users are willing to omit cell-type specific peaks and restrict analysis to peaks with >100 reads in both cell populations, precise data-quality matching becomes less critical, since the dominant factor in our benchmarking was peak read depth. Although this filtering may reduce how representative differential-binding effect sizes are for the entire population, it substantially improves reliability and reproducibility. In our datasets, applying a 100 peak-read threshold retained roughly 100,000 peaks with 4,000 cells (Figure S8A). This is especially useful for low-cell-number datasets, where it is better to evaluate well-powered regions than risk high false negative rates.

We also recommend limiting all locus-specific analyses to regions where both populations have sufficient coverage (>100 reads), as this provides the most reliable way to identify population-specific footprints. Otherwise, high false negative rates can lead to many incorrectly reported differences. We found PRINT was the most straightforward tool for locus-specific analysis.

For users with <1,000 cells, FRiP < 0.1, or low peak-read counts, we do not recommend local or global footprinting. Instead, consider alternative analyses like motif enrichment (e.g. HOMER^42^, ArchR^43^), which can still be informative if peak calling remains reliable.

Regardless of the tool or workflow, we found:

1. Footprinting on a single sample highlights broadly active and GC-rich TFs; the strength of footprinting lies in comparison, so we always recommend introducing a baseline.
2. Motif-centric methods suit most needs, but de novo approaches are better for exploring non-canonical motifs or avoiding PWM-related issues.
3. When calling peak regions, more conservative filtering can reduce noise. At a minimum, we recommend a MACS3^44^ peak-summit filter of 20 (Figure S8B).
4. If TFs of interest are known, consider the GC content and the added predictive value of footprinting over motif scanning (e.g., Figure 7J; TRACE^17^ Figure 4B). CTCF is a great candidate, whereas NFATC3 may be difficult to interpret.
5. RNA-seq can support differential binding interpretation but was not effective as a filtering step.
6. Users integrating results across footprinting tools will likely find union-based methods retain more biologically relevant information than intersecting sets.
7. Continuous footprint scores are typically more informative than binary bound/unbound calls. When possible, reformulate analyses to leverage this – for example, by weighing differentially accessible regions by maximum footprinting scores to prioritize candidate regions of interest.

For future developers, we recommend directly assessing sample-to-sample variation and implementing user safeguards. Adopting a standardized file set would greatly improve comparisons and user experience as well. Specifically, we suggest releasing: (1) Tn5 bias-corrected reads (bigWig), (2) Combined peak- and footprint-coordinates (BED8), (3) support for predefined TFBSs sets, and (4) aggregate footprints as matrices, not images.

## DISCUSSION

Using a new downsampling framework, scBAMpler, and an extensive read-level benchmarking dataset we evaluated HINT, TOBIAS, and PRINT with diverse approaches, revealing the substantial impact of methodological choices. For instance, motif-centric tools largely consider distinct TFBSs, making their limited overlap unsurprising. Though we highlight many areas for improvement, our findings support a promising future for single-cell genomic footprinting, where consistent and reproducible analyses are obtainable with intentional bioinformatic strategies. Notably, our results confirm that large fluctuations in sample properties significantly impact footprint detection and comparing samples across this spectrum risks false positives from detection bias. This challenge is amplified in differential binding analyses when tools overlook distributional differences, leading to highly confounded outcomes. Consistent with prior work^13,18,40,41^, we observed performance dips around 100 million reads in both our consistency and equivalency analyses. However, our work clarifies the underlying causes. In consistency analyses (non-downsampled versus downsampled), performance loss stems from false negatives in regions with <100 reads. In equivalency analyses (ChIP-seq versus footprinting), true positive rates remain stable, but ATAC-seq’s ability to mirror ChIP-seq declines below 100 million reads. This clarifies that false positives are not linked to read depth, but false negatives are.

Because each tool prioritizes fundamentally different information, it is difficult to name a single best program. PRINT delivers conservative, consistent footprints while remaining sensitive to read support. Its scores effectively identified cell line–specific TFs and had a comparable ChIP-seq precision. However, its advanced features, like multi-scale footprinting and nucleosome detection, comes at a high computational cost and a workflow better suited to advanced users (Figure S8C). TOBIAS achieved the highest precision to ChIP-seq and features a user-friendly codebase, but appeared rather agnostic to read distributions within peaks. HINT performs similarly to other methods in consistency analyses but was limited by a footprint score that was the least effective at ranking strong footprints. Overall, tools identify TFs with varying strength, suggesting method choice may favor certain TFs and that ensemble scoring could offer the best results.

Overall, we were interested to find that many limitations of footprinting stem from constraints of motif matching. This step determines which sites motif-centric algorithms evaluate, affects detection power, and introduces high redundancy—making interpretation difficult without predefined TFs of interest. While PWMs are experimentally derived, this does not guarantee that they can fully or precisely capture TF behavior across diverse cellular contexts. Recent advances in motif modeling could substantially benefit the footprinting field^42,45–47^.

While we aimed to fairly evaluate each method, several caveats remain. First, future work is needed to clarify how Tn5 bias correction impacts across-tool reproducibility, particularly whether superimposition differentially affects TFs. Second, we used a full PWM database to reflect how users might generate hypotheses, though this adds noise compared to starting with candidate TFs. Third, we did not apply our guidelines in the context of a biological question-driven scATAC-seq analysis, choosing instead to focus on analyses with a ground truth to enable meaningful interpretation of performance differences.

As more tools emerge to distill and transform chromatin accessibility data into TF binding predictions^26,48–52^, it will become increasingly crucial to evaluate and apply them effectively. While powerful, deep-learning models often operate at such a high level of abstraction that effort will be needed to preserve interpretability and future benchmark utility. Our analyses highlight that despite a shared goal, footprinting tools differ substantially in their underlying assumptions and outputs–often more than casual users expect. This is exacerbated by the lack of a definitive ground truth for most TF binding contexts. Without greater transparency and modularity in tool design, these challenges will continue. We recommend broader adoption of practices like those in TOBIAS, which provides well-structured intermediate outputs that enable clearer comparison and integration across methods.

This reflects a broader debate—both practical and philosophical—about how much fields should be penalized for lacking clear validation benchmarks. If a user’s ultimate goal is to predict ChIP-seq signal, alternatives like simple motif scanning or histone marks (e.g., H3K27ac) likely outperform footprinting. Many artifacts in existing evaluation paradigms—including our own—limit the ability to highlight what footprinting uniquely offers. Single-cell footprinting can identify *in vivo* binding events in primary tissues, developmental stages, or intermediate cell types that are often unculturable or poorly represented in reference datasets. Moreover, de novo footprinting methods can detect entirely novel binding sites that do not match any known PWMs, capturing context-specific regulation other approaches miss. Future work could extend our benchmarking framework to test whether including de novo footprints (e.g. HINT’s derived binding sites without PWM annotations) improves ChIP-seq concordance. Finally, footprinting could inform and refine PWM development itself—ironically, one of the limitations we provide. This circularity highlights a central issue: footprinting’s greatest strengths may lie precisely in contexts where standard benchmarks, like ChIP-seq concordance, are least effective.

Genomic footprinting offers a broad, cost-effective approach to extract insights from the expanding set of single-cell ATAC-seq data. Given the context-dependent nature of TF interactions with DNA, identifying differential binding across cell types—especially in developmental or disease contexts—can be highly informative. Although performance varies by TF, strong results for even a few make footprinting valuable. Early limitations, like Tn5 bias, are now better understood, and current challenges—like PWM-match accuracy and pseudobulking—have promising solutions. Amid the growing momentum behind genomic footprinting, we present both practical guidelines for single-cell implementation and a strategic framework for revealing gene regulation diversity in complex tissues.

## RESOURCE AVAILABILITY

This paper analyzes existing, publicly available data, accessible at ENCODE^31^. The codebase for scBAMpler is available at https://github.com/aseveritt/scBAMpler. All accompanying data, code, and analyses are deposited at https://github.com/aseveritt/scFootprintBenchmark. Any additional information required to reanalyze the data reported in this paper is available from the lead contact upon request.

## ACKNOWLEDGMENTS

We thank Jenelle Wallace and David Quigley for reviewing the manuscript. We thank Alex Pollen and Ryan Corces for their helpful guidance on the project. We thank Cindy Pino, Caitlin Brown, and Hasan Alkario for their helpful feedback on figure creation. This work was supported by the National Institutes of Health (grant#MH123178, #AG087959, and #MH134981), the Biswas Transformative Computational Biology Program, and Gladstone Institutes. A.E. was additionally funded by the UCSF PhD program in Biological and Medical Informatics and ARCS Foundation.

## AUTHOR CONTRIBUTIONS

A.E. performed all analyses and prepared the manuscript. S.W. helped guide the project and performed ensemble modeling. K.S.P supervised all aspects of the work. All authors helped conceptualize and design the project and edit the manuscript.

## METHODS

### Generation of snATAC-seq dataset

Single nuclei ATAC-seq data for 11 cell lines constituting 23 total samples were obtained from the ENCODE portal as bam files aligned to the human reference genome (GRCh38)^31^. Fragment files were constructed with Sinto (v0.10.0)^55^ and used as input to ArchR (v1.0.2)^43^. The dataset was filtered to remove doublets (filterRatio = 1) and cells which did not cluster in UMAP space with their cell line of origin. The final dataset included 256,150 cells.

Because footprinting is performed at the read-level, we wanted to generate filtered bam files that exactly replicated the filtered dataset. The original bam files were filtered to remove reads that (1) originated from a removed cell barcode (2) aligned outside of the core genome (defined as chr1-22, X, Y), and (3) failed to pass Sinto’s fragment quality filters (e.g. MAPQ score, PCR duplicates, exceeding maximum fragment size). This required an in-house edit of Sinto such that reads-names are an optional output file.

Peaks were called for each cell line using MACS3 (v3.0.1)^44^ and parameters (-g 2.9e+09 --call-summits -- keep-dup all --nomodel --nolambda --shift -100 --extsize 200 -q 0.05). Similar to others’ findings^30^, we found MACS3 and HMMRATAC performed the best as peak callers. Some footprinting tools, like PRINT, require peaks to have fixed-widths. Therefore, peak summits were filtered and standardized to 500bp using an in-house implementation of ArchR’s iterative overlap procedure. Unreliable genomic regions which overlap more than 25% of a region from the cellranger GRCh38 exclusion list were also removed. A “union” peak file sourcing summits from all 11 cell lines was similarly constructed for the cell-similarity analyses.

ENCODE accessions: ENCFF542OFM, ENCFF282TVW, ENCFF283AEI, ENCFF505ZAE, ENCFF089OZP, ENCFF525ZMO, ENCFF851IFA, ENCFF473MZT, ENCFF834CUN, ENCFF866BBY, ENCFF180UYN, ENCFF767NVC, ENCFF948XGC, ENCFF213ZUI, ENCFF853WXT, ENCFF300ZOG, ENCFF880AYO, ENCFF703NGW, ENCFF032WCD, ENCFF463CKH, ENCFF443RDC, ENCFF012ZOS, ENCFF322BLG

### scBAMpler development

#### Downsampling procedure

To systematically assess the impact of data quality on footprinting, we developed scBAMpler, a tool for read-level downsampling that modifies one aspect of data quality at a time—while preserving the original distribution of reads across cell barcodes. For each cell type, scBAMpler constructs a dictionary where each key is a cell barcode and each value is a tuple containing (i) mapped read names, (ii) the total read count, and (iii) which reads map to peak regions (defined as having ≥75% overlap with a peak). To reduce storage overhead, all data is numerically encoded.

Using these dictionaries, scBAMpler generates lists of read names which are then used to subset the original BAM files to produce the final downsampled datasets. Subsampling is repeated with different random seeds to ensure reproducibility.

Three types of downsampling are currently supported: cells, reads, and FRiP. To downsample by cell, *N* cell barcodes are removed and all associated reads are discarded. To downsample by read depth, *N* read pairs are removed, proportionally distributed across cell barcodes based on each cells’ total read count. To decrease FRiP, *N* peak read pairs are removed (proportional to each cell’s peak read count); conversely to increase FRiP, *N* non-peak read pairs are removed. The tool takes as input a BAM file, a peak file, and downsampling parameters, and outputs a subset BAM and/or fragment file. scBAMpler relies on SAMtools^57^, bedtools^58^, pandas^59^.

In total, we generated 321 downsampled BAM files, which represents triplicates of 6 read-downsampled, 9 cell-downsampled, and 7 FRiP-downsampled conditions across the five cell lines (K562, HepG2, MCF-7, GM12878, SK-N-SH). For each dataset, footprinting was performed using the original cell line–specific peak file.

#### Cell-similarity procedure

To evaluate how cell-to-cell heterogeneity impacts footprinting, we extended scBAMpler to support multi– cell-type (or, in this manuscript, multi–cell line) inputs. Our goal was to ground the analysis in a distance metric based on the peak-by-cell count matrix so that it is more comparable across experiments and less reliant on users’ choices for dimensionality reduction. Conceptually, scBAMpler generates random combinations of cells using a top-down strategy that allows precise control over peak read depth, FRiP, cluster cohesion, and distance to a reference population. This approach is designed to be computationally tractable while still covering the parameter space observed in real datasets.

A naive strategy would be to shuffle cells randomly to simulate hypothetical populations; however, computing pairwise distances between all cells quickly becomes infeasible at large scale. Instead, scBAMpler uses constrained K-means clustering on a precomputed dimensionality-reduced embedding (e.g., UMAP, t-SNE, cisTopic space) to group cells into clusters of exactly 100 highly similar cells. This strategy effectively compresses the space of cells—for example, reducing 200,000 cells to 2,000 centroids—while preserving local similarity. The peak-by-cell matrix is normalized using term frequency– inverse document frequency (TF-IDF) prior to computing the average peak-count profile for each centroid. Pairwise Pearson correlations are then calculated across all centroid profiles to construct a Euclidean distance matrix. The input to this step includes the raw peak-by-cell count matrix and a corresponding low-dimensional embedding, stored in HDF5 format. The output consists of a dataframe describing the centroid populations, associated summary statistics, and the correlation matrix used for downstream population construction.

Using these inputs, scBAMpler randomly shuffles centroids to efficiently generate hypothetical cell populations that span a broad range of characteristics, including peak read depth, cluster cohesion (measured as the sum of squared Pearson correlations to the centroid), and distance to known cell-type profiles. Users can filter or select populations of interest based on these metrics. For each selected population, scBAMpler outputs a BAM and/or fragment file, along with a summary statistics file. These BAM files may include reads from multiple cell types, depending on the composition of the selected population. This expanded functionality relies on KMeansConstrained^56^, scipy^60^, Sinto (v0.10.0)^55^, SAMtools^57^, bedtools^58^, and pandas^59^.

In this study, we applied scBAMpler to the dataset comprising 256,150 cells from 11 cell lines, using a peak-count matrix constructed from the union peak file. These cells were grouped into 2,561 centroids of 100 cells each (See Figures S3A-B). By shuffling centroids many times, we generated ∼300,000 hypothetical cell populations, ranging in size from 500 to 20,000 cells (See Figure S3C). To focus on representative populations, we excluded outliers with cluster cohesion < 0.06 or a pseudobulk FRiP outside the range 0.18 to 0.28. Because we wanted to distinguish the influence of cell homogeneity from peak read-depth, we selected four peak read-depth categories: 5e7, 1e8, 1.5e8, and 2e8 which is approximately the 25th, 50th, 75th, and 95th percentile. Within each peak read-depth category, we selected 20 populations with varying distances to a chosen reference cell line profile, resulting in a total of 80 hypothetical cell-populations for footprinting (See Figure S3C).

### Footprinting parameters

The Regulatory Analysis Toolbox (RGT) v1.0.2, which includes HINT^14^, was installed via pip. For each dataset, HINT was applied with parameters: rgt-hint footprinting --organism hg38 --atac-seq --paired-end; rgt-motifanalysis matching --organism hg38 --motif-dbs JASPAR2022. Occasionally, HINT footprint regions fall outside, but near, the input peak regions. We required 50% of the footprint region to fall within a peak region to be retained.

TOBIAS^15^ v0.16.2-b was applied with default parameters for ATACorrect, ScoreBigwig, BINDetect. Analyses were performed on the BINDetect output *overviews.txt files where bound=1 unless otherwise stated.

The PRINT repository was cloned from github (https://github.com/HYsxe/PRINT) on 03/15/2023^26^. The pre-computed whole genome bias file for hg38 were used at the expected insertions (“loadTFBSModel”). For each condition, the groupInfo and barcode grouping were set to 1 for all cell barcodes (“barcodeGrouping”). The dispersion models between 2 and 199 were applied (“dispModel”). Count tensors were computed (“getCountTensor”) and footprinting scoring was performed at scale 30 (“getFootprints”). Finally, possible TFBS (“getTFBS”) and footprint calls were evaluated (“getTFBindingSE”). The final “TFBindingSE” object was output as a bed file; analyses were performed using 0.3 as the bound footprinting score threshold, approximately the 65th score quantile in original samples, unless otherwise stated.

Genome- and peak-wide PWM matches were generated with the R package monaLisa (v1.6.0)^54^ using parameters (findMotifHits(method=”matchPWM”, min.score=6).

For all tools, the JASPAR 2022^35^ core vertebrate non-redundant list was used as the PWM database (n=841). PWM motif-clusters, DNA-binding domain, and motif-family were accessed via the jaspar.genereg.net API.

### Evaluation methodology

#### Within-tool comparisons: downsampling

For each tool, local footprinting results (often BED files) were transformed into a matrix where rows represent cell-line peak regions (OCRs), columns correspond to TF motifs (PWMs; n = 841), and each cell contains the number of footprints detected for that PWM in that peak region. For HINT, this was generated from the *_mpbs.bed files; for TOBIAS the *_overviews.txt where bound=1; and for PRINT the *_granges.bed file where habituation score ≥ 0.3. The matrix generated from the original dataset for each tool was treated as the ground truth. We tested various matrix formations (e.g., OCR × TF, OCR × TF-family, footprint coordinates × TF) and found the final conclusions were unchanged.

To evaluate performance, we defined true positives (TP) as peak:TF pairs found in both the original and downsampled matrices. False negatives (FN) were pairs present only in the original matrix, and false positives (FP) were pairs found only in the downsampled matrix. True negatives were not used in our assessments. Precision, recall, and F1 scores were first calculated per PWM for each dataset. Because we observed high reproducibility across triplicates of the same conditions, we averaged the PWM scores across triplicates. In figures that show a single point per condition (e.g., Figures 2A, S2B, S2D-H, and S2J-K), we report the mean ± standard deviation of the averaged PWM F1 distribution (i.e., the macro-average). This provides information about global PWM performance rather than triplicate performance, which we found more informative. In selected analyses where PWM match was not considered part of the true-positive criteria (e.g., Figures S2H and S2K), the matrix was simplified to a single-column format where each entry represents the total number of footprints in a region, regardless of TF identity.

To calculate peak read coverage, we used bamCoverage (v3.5.5) ^61^ to generate BigWig files from each BAM file. The signal was summed over each peak region to calculate the read depth per peak. These values were used to calculate the average peak read coverage per sample (See Figures 2B and S2C), assign F1 scores to individual peak regions (See Figure 2C), and evaluate performance on coverage-filtered subsets of peaks (See Figure 2D).

#### Within-tool comparisons: cell homogeneity

For each of the 80 selected hypothetical cell populations, local footprinting results were transformed into a matrix of union peak regions by TF motifs, as described above. A corresponding matrix was also generated for each of the four reference cell lines (HepG2, K562, MCF-7, SK-N-SH), using all cells assigned to that line. Each hypothetical population was constructed relative to one of these cell lines— that is, the population was selected based on its average distance to a specific cell line centroid. The footprinting matrix from the corresponding reference cell line was used as the ground truth for evaluating performance metrics (precision, recall, and F1 score).

Unlike earlier analyses, there are no explicit triplicates for these synthetic populations. Therefore, we report the mean performance across all PWMs for each dataset (See Figures 3A and S3D). To help visualization, LOWESS curves (locally weighted regression, frac = 0.4) were fit for each peak read-depth category. As in previous sections, we observed that TOBIAS’s dynamic bound threshold resulted in less structured performance results. Therefore, to improve cross-comparisons, we fixed the bound threshold at 0.025 for this analysis—roughly equal to the average threshold used across the four reference cell lines.

#### Across-tool comparisons

Using the previously described OCR × PWM count matrices, we directly calculated how often cell values aligned across tools. We report both the absolute intersection size at each tool’s default bound-threshold (See Figure 4A) and the Jaccard coefficient across deciles of each tool’s score distribution (See Figures S4A–C). Notably, overlap was not substantially higher when using simplified, PWM-agnostic matrices in which cells represent the total number of footprints per region, regardless of TF identity.

For the motif-centric tools, we also generated peak × PWM matrices *prior* to applying any bound-threshold, representing the full set of candidate TFBSs considered by each method (See Figure 4B). Given the low overlap observed, we selected two cell lines and re-called footprint regions with TOBIAS, adjusting the BINDetect p-value cutoff from 1×10⁻⁴ to 1×10⁻⁵—matching the threshold but not algorithm used by PRINT—and found similarly low overlap.

We ranked PWMs by the total number of footprints detected per sample (i.e., column sums of the peak × PWM matrix), assigning rank 1 to the most frequently observed motif. These rankings showed strong agreement across tools, as reflected by high weighted Kendall’s tau values (computed via scipy.stats.weightedtau^60^). In contrast, rankings based on the total number of TFBSs in peak regions from the R package monaLisa^54^ showed lower concordance (See Figure 4C). We also performed motif enrichment analysis using HOMER^62^ (findMotifsGenome.pl -hg38r -size 200 -nomotif -genomeBg - mknown) on each original sample, ranking PWMs by 1 – log₁₀(*p*-value). These rankings also showed limited similarity to footprint-based ranks (See Figure S4E).

#### PWM property analyses

To compare PWM performance across downsampling conditions without bias from absolute performance values, we first Z-scored the PWM-level F1 scores within each dataset. When concatenated, this produces an 841 × 963 matrix, with rows representing motifs and columns representing 321 downsampling conditions applied across 3 tools. These Z-scores were highly consistent across triplicates, cell lines, tools, and downsampling types—indicating that although the number of footprints per TF may vary, their relative rankings remain stable for the vast majority of motifs. An exception to this pattern was cell-type-specific TFs, whose rankings deviated from their expected position—highlighting how using a full PWM database, even when including inactive TFs, can help identify these biologically meaningful outliers.

To explore what governed these stable rankings, we focused the analysis on read-downsampling conditions for clarity and simplicity. We averaged the F1 Z-scores across all read-downsampling conditions to produce a single robust value per PWM, resulting in an 841 × 15 matrix, with rows representing motifs and columns representing 3 tools applied to 5 cell lines. We then tested whether specific PWM features explained the observed rankings. Using custom scripts and the monaLisa^54^ R package, we extracted the following features from each JASPAR2022 motif (represented as a position probability matrix): GC content, repeat content, PWM length, information content (IC, using the Schneider correction), genome-wide TFBS prevalence in hg38, and Universal Stripe Factor^11^ (USF) membership.

Pearson correlation coefficients were used to evaluate associations between numeric PWM features and average F1 Z-scores; point-biserial correlation was used for the binary USF membership. We found consistent positive associations with both GC content and USF membership, but no consistent trends for repeat content, PWM length, or genome-wide TFBS prevalence (See Figures 5A and S5A–B).

To further evaluate the observed GC content trend, we generated synthetic PWMs by training a Dirichlet mixture model on the position count matrices (PCMs) from the JASPAR 2022 database. Synthetic PWMs were then sampled according to the criteria outlined in the TFBSTools^63^ R package (v1.38.0; dmmEM(K=6, alg=“C”)). To fully capture the diversity of real motifs, we trained separate models on (i) the entire JASPAR database (n=114), (ii) individual PWM clusters with more than 30 members (n = 210), and (iii) a subset of USF motifs (n=30). Each resulting PCM was converted to a position probability matrix (PPM), and motif properties were calculated using the same in-house code described above. We selected synthetic motifs to span realistic ranges of IC, length, and genome prevalence observed in real JASPAR PWMs (See Figures S5C–D). A small number of outliers were excluded based on the following criteria: IC < 5, IC > 25, repeat content > 70%, or chromosome 10 prevalence > 2.5 × 10⁶. In total, we generated 262 synthetic PWMs, which were then supplied to each footprinting tool and analyzed under the same read-downsampling conditions as the original motifs.

We reasoned that if PWM occupancy (#TFBS bound / total TFBS) depended mainly on sequence features, then a linear model trained on synthetic PWMs should generalize to JASPAR PWMs, and vice versa. For each cell line, we created 70/30 train-test splits separately for the synthetic and JASPAR PWMs. For each tool, we fit an ordinary least squares (OLS) regression model using the formula: Percent Occupied ∼ GC Content * IC * Repeat Content. We evaluated the model’s R² in four settings: training and testing on synthetic PWMs, training on synthetic and testing on JASPAR PWMs, training and testing on JASPAR PWMs, and training on JASPAR and testing on synthetic PWMs. The mean and standard deviation of R² values across all cell lines and tools are reported in Table 1.

#### Differential binding analyses

For each tool and cell line, we Z-scored the number of footprint regions per PWM to normalize across motifs within a dataset. To assess cell line specificity, we then computed the deviation of each PWM’s normalized score from its average across the other four cell lines: (observed – mean expected) / SD expected.

To identify TFs with cell line-enriched RNA expression (CLE-TFs), we obtained transcripts per million (TPM) values from ENCODE RNA Expression Reports (polyA+ assay). For each gene, we calculated the observed TPM versus the expected TPM across the other cell lines using the same formula. TFs in the top 20% of this distribution were designated as CLE-TFs.

To evaluate whether footprinting tools prioritize CLE-TFs, we compared the observed-expected scores for CLE-TFs to 10,000 random TF sets of the same size. Empirical p-values were Benjamini–Hochberg corrected (See Figure 5B). For each cell line, we show an UpSet plot of the top 5% ranked TFs and a heatmap of the top 15 TFs, highlighting CLE-TF members (See Figures 5C–E and S5E–H). We repeated this analysis on other matched cell line conditions (10K, 5K, 1K) as well as the unmatched original datasets (HepG2: 25043, K562: 48060, MCF-7: 32303, GM12878:30246, SK-N-SH:60714 cells) and evaluated how consistent the top 5% of TFs remained (See Figure S5I).

To evaluate the differential binding modules provided by HINT and TOBIAS, we tested three comparison settings: (i) K562 vs HepG2 using the original samples, (ii) matched K562 samples (within-condition), and (iii) unmatched K562 samples (across-condition). For TOBIAS, each sample pair was analyzed using the BINDetect module with default parameters. For HINT, differential footprinting was performed using the command ‘rgt-hint differential --organism hg38 --bc’. To facilitate direct comparison between tools, HINT fold changes were computed as: log₂(protection score_rep1_ + tag count_rep1_) – log₂(protection score_rep2_ + tag count_rep2_). Results for the cell-downsampling comparisons are presented in Figures 6A–B, with the full set shown in Figures S6A–C.

#### ChIP-seq Peak Processing and PWM Evaluation

ChIP-seq data for the five core cell lines were sourced from the ReMap2022 database (remap2022_*_nr_macs2_hg38_v1_0.bed)^53^. For each cell line, we identified the shared set of TFs present in both the JASPAR2022 database and the corresponding ReMap file (GM12878: 89/155, HepG2: 131/285, K562: 165/413, MCF-7: 78/167, SK-N-SH: 30/44). ChIP regions were intersected with cell line-specific snATAC-seq peaks, requiring at least 50% overlap of the ChIP region. The results were converted into matrices where rows represent shared peak regions, columns correspond to TFs, and cell values indicate the number of ChIP peaks for each TF within each ATAC peak.

To assess which ChIP:TF signals could be reasonably predicted from their PWM, we measured how often ATAC peaks contained both a ChIP peak and the corresponding PWM sequence.Because results were similar whether using binarized or integer matrices, we used binarized matrices for all performance calculations, setting any value greater than 1 to 1. For each ChIP’d TF, we tested three matching scenarios: (i) the matched PWM, (ii) the matched PWM with randomized locations (i.e. permuted row values), and (iii) unmatched PWMs (See Figures S7A–B). Using these results, we refined our set of high-confidence ChIP TFs to those with had (i) recall > 0.4, (ii) F1 score > 0.05, and (iii) their matched PWM produced the highest F1 score. If multiple PWMs were available for a single TF, the PWM with the highest F1 score was retained. The resulting number of TFs passing these filters per cell line were: GM12878: 30/89, HepG2: 63/131, K562: 65/165, MCF-7: 36/78, and SK-N-SH: 14/30.

To evaluate how downsampling affects the overlap between ChIP-seq regions and ATAC-seq peaks, we used bedtools to compute two metrics for each condition: (1) the number of ChIP regions with at least 50% overlap with an ATAC peak, and (2) the total number of base pairs from ChIP regions overlapping ATAC peaks. Both metrics plateaued quickly across downsampling conditions, indicating that beyond a certain threshold, further reductions in data quality have minimal impact on recovery. However, the absolute percentage of overlap was highly variable across TFs (See Figure S7C).

#### ChIP-seq comparisons

For each tool and cell line, footprinting results were converted into a binary matrix: rows represent scATAC-seq peak regions overlapping ChIP-seq regions, columns correspond to retained PWMs, and cell values indicate whether a footprint was detected in that region. The previously described binary matrix generated from ChIP-seq data was used as ground truth.

We began by analyzing the original (non-downsampled) datasets to test how different footprint score thresholds affected performance metrics. For each tool–cell line pair, we calculated every 10th quantile of the footprint score distribution. At each decile, only footprints exceeding the corresponding threshold were included in the performance calculations. The decile yielding the highest F1 score was defined as “optimal.” We then compared this to the decile closest to each tool’s default threshold to evaluate whether tuning—per TF, across TFs, or across cell lines—could improve performance (see Figure 7B and S7G; 7A; and S7D, respectively).

Additionally, we assessed how alternative tuning strategies influenced performance, all of which involved subsetting either the ChIP or footprint matrix (see Figures S7E–F). The comparisons included:

- Standard comparison: all common peak regions and default thresholds (See Figures 7C and S7H)
- Shuffled footprint locations: row-wise permutation of the footprint matrix
- ChIP regions with PWM match: restrict rows to peaks that also contain the PWM sequence
- Optimal score threshold: apply the optimal threshold which maximized F1 scores
- ATAC peaks with >100 reads: restrict rows to peaks with sufficient ATAC read coverage
- Top 5000 footprinted regions: restrict to the highest scoring footprint regions for each TF
- All TFBSs: include all candidate motif matches, with no score threshold (motif-centric tools only)

Because performance trends varied widely across TFs, we used the standard comparison for evaluating changes across downsampling conditions (e.g., Figures S7I–J).

To investigate which features explain the change in F1 scores across downsampling conditions, we fit ridge regression models (cross-validation = 10, alpha = 0.5) for each TF-tool pair. Predictor variables included cell line, FRiP, number of cells, number of peak read pairs, total number of read pairs, percent overlap between ChIP regions and ATAC peaks (in basepairs), and the F1 score of the PWM alone (as calculated in the “ChIP-seq Peak Processing and PWM Evaluation” section). To avoid poorly performing fits, only models with R² above the median were included. For the remaining models, the mean and standard deviation of feature coefficients were summarized as radar plots (See Figure 7D).

#### Ensemble models

The original HepG2 scATAC-seq peak regions were annotated with the maximum footprint score from HINT, PRINT, and TOBIAS for two TFs–CTCF and JUND–using their best performing motif as described above (CTCF_MA0139.1 and FOS::JUND_MA1141.1). For all combinations of footprinting tools, a gradient boosting classifier (scikit-learn^64^ 1.6.1, HistGradientBoostingClassifier, max_depth = 3) was trained using footprint scores as features and binarized ChIP-seq peak overlaps as labels. auPRC (average precision) was calculated on held-out chromosomes using stratified group k-fold evaluation (n_splits = 5, shuffle = True), and the average auPRC across folds was reported. If a tool calls no footprint within a peak, the score is represented as a missing value (NaN); gradient boosting natively handles missing values without needing to replace them with an arbitrary numeric value like 0.

