## Supplemental Figures for "Comparative evaluation of genomic footprinting algorithms for predicting transcription factor binding sites in single-cell data"

Everitt et al. 2024

1A

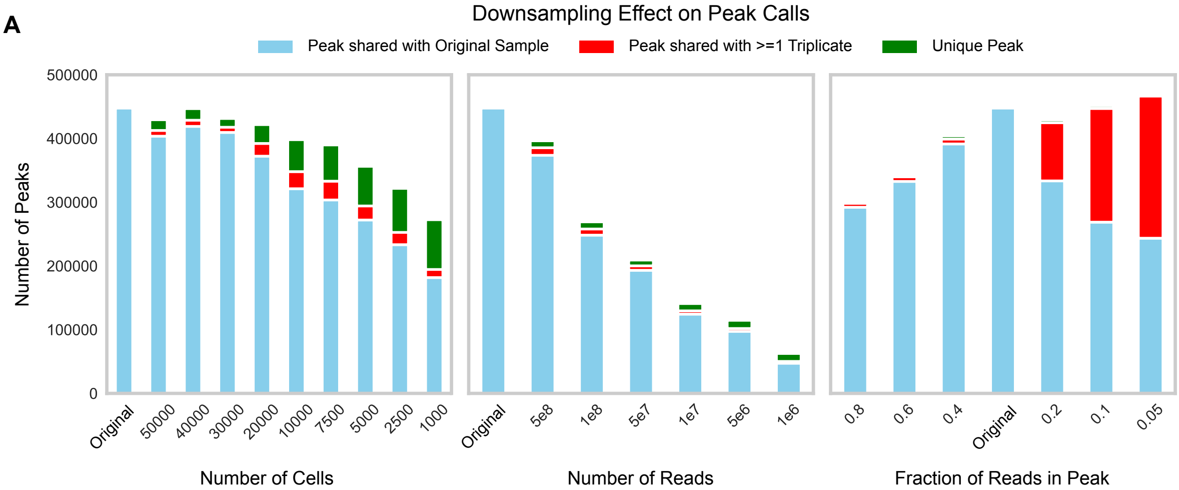

**Figure S1A) Stacked bar chart showing how downsampling affects peak calling.** Using identical peak-calling commands, we evaluated the similarity between peak regions identified in the original samples versus the downsampled BAM files. Peaks were considered overlapping if they shared at least 30%, or 150bp, with an original peak. Peaks retained from the original samples are shown in blue. As cell counts are reduced, original peaks are largely preserved until extreme downsampling, where peaks unique to a single triplicate (red) or shared between triplicates but not with the original set (green) begin to emerge. Downsampling reads led to a linear loss of shared peaks. Similarly, increasing FRiP (via removal of non-peak reads) leads to a sharp reduction in shared peaks. Conversely, decreasing FRiP (via removal of peak reads) introduces more unique peaks, due to changes in signal strength. Bar height reflects the mean values across the five cell lines (HepG2, K562, MCF-7, GM12878, SK-N-SH).

2A

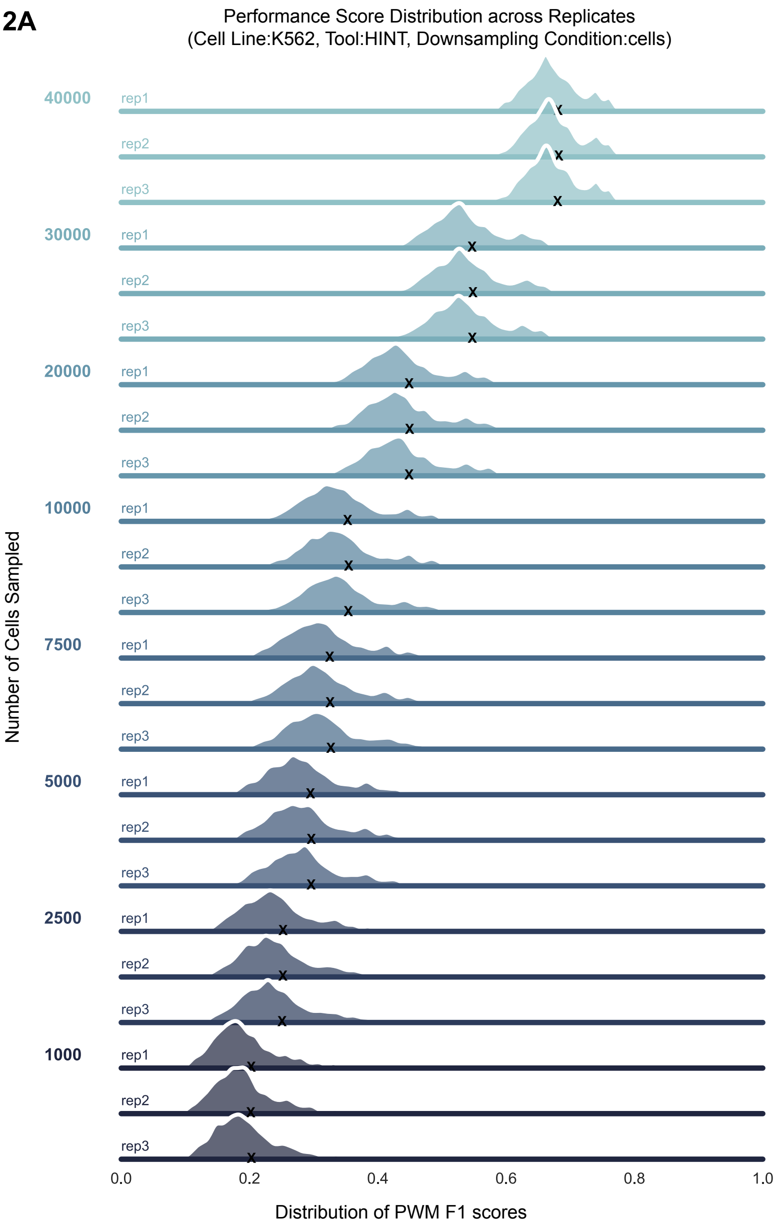

**Figure S2A) Stacked ridge plot depicting similar PWM F1 score distributions across triplicates.** Each triplicate produced highly consistent F1 score distributions, motivating the use of the mean F1 score across triplicates for subsequent figures. Downsampling conditions are labeled on the left, each line represents a replicate, with black Xs indicating distribution means. For illustration, HINT scores from downsampled K562 cells are shown, though the trend was consistent across all cell lines and tools.

## 2B

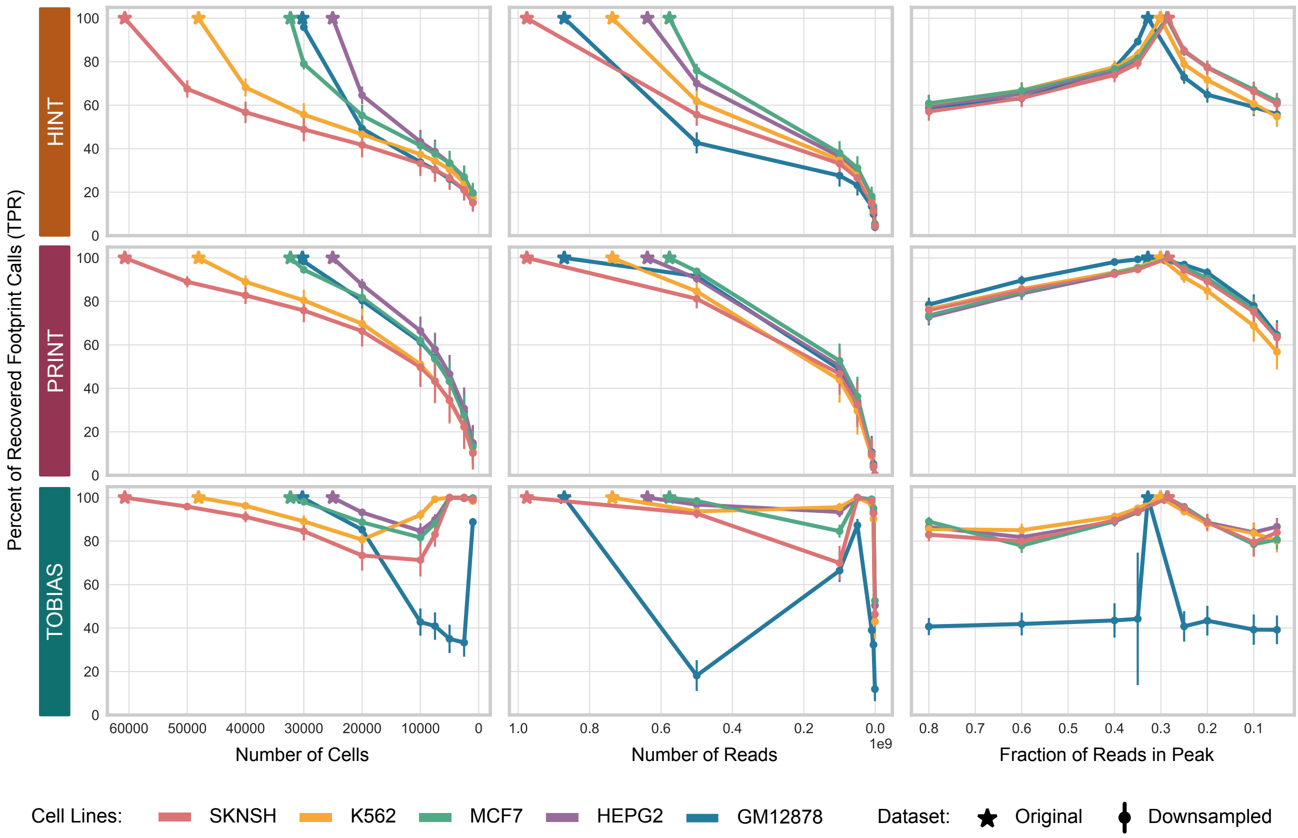

**Figure S2B) Line plots illustrating the percent of footprint regions in the original sample (star) recovered in downsampling conditions (dots).** Points show mean ± SD across triplicates. Each line represents a cell line; rows are tools, columns are downsampling conditions.

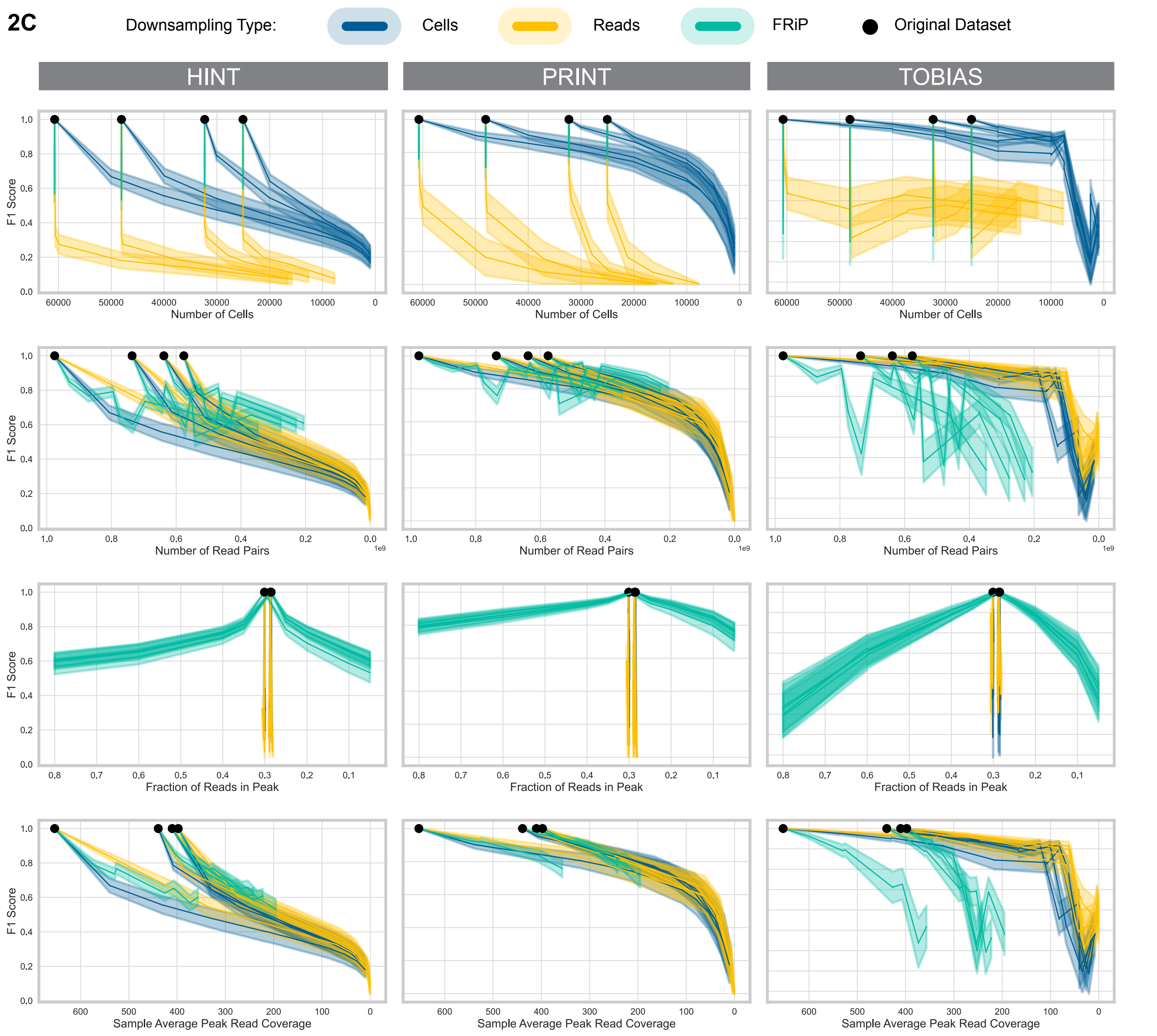

**Figure S2C) Peak read coverage is the only consistent predictor of F1 scores across all downsampling conditions.** Unlike Figure 2A, which shows one downsampling type per plot, this figure presents all three sampling types together. If a single property drove F1 score performance, we would expect a synchronized decline across all cell lines and sampling types. Each plot displays results for four cell lines (HepG2, K562, MCF-7, SK-N-SH), with the initial (baseline) value marked with a black dot. For each cell line, three downsampling strategies are represented by color with the mean F1 score ± SD across triplicates.

- Row 1: F1 scores decline more rapidly in the reads (yellow) and FRiP (emerald) conditions, despite similar cell counts to the cell condition (blue).
- Row 2: Equal read counts result in similar F1 scores across conditions, though FRiP shows greater, unstructured variability.
- Row 3: F1 scores shift in the cells and reads conditions – producing a practically vertical line – despite similar FRiP values.
- Row 4: When average peak read coverage is matched, F1 scores remain consistent across all sampling conditions and cell lines. This pattern holds for both HINT and PRINT, while FRiP continues to influence TOBIAS estimates more than expected.

**2D**

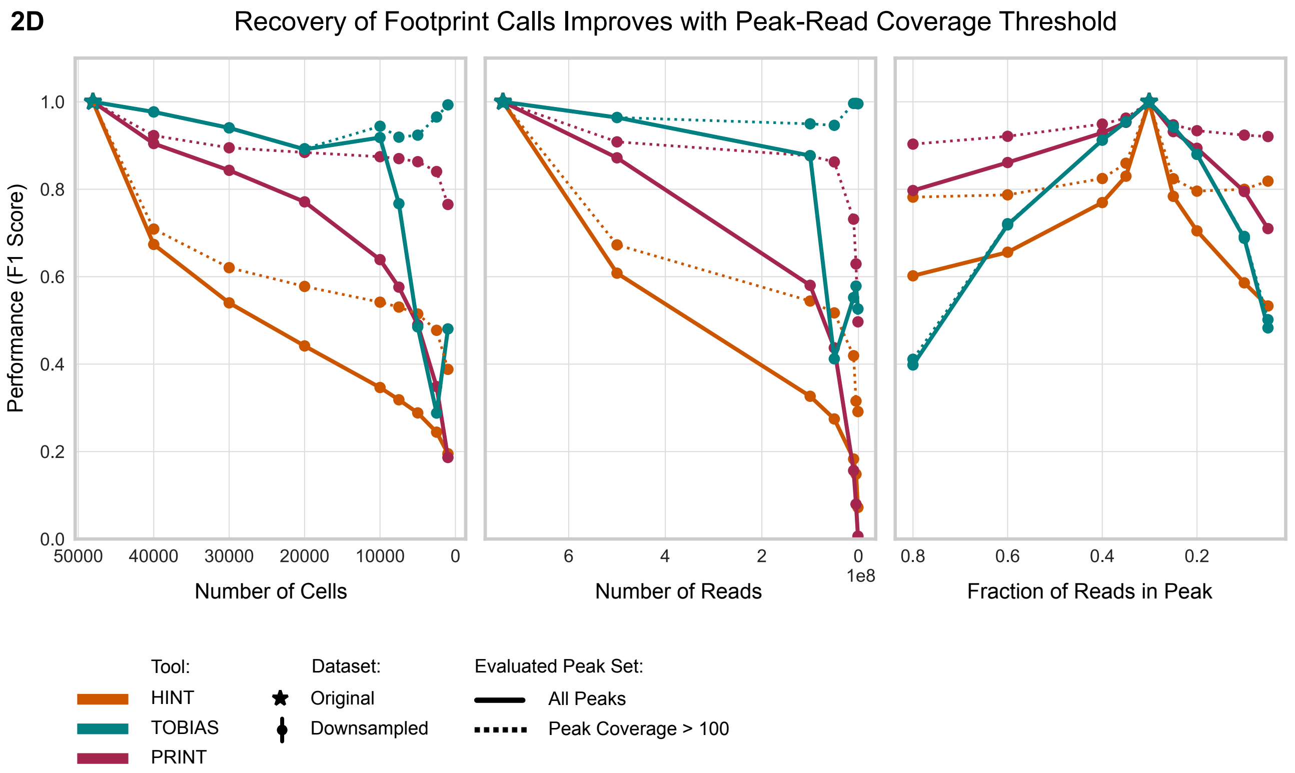

**Figure S2D) Performance improves when analysis is restricted to peaks with sufficient read coverage.** F1 scores are shown for all peaks (solid lines) and for peaks with read coverage > 100 in both original and downsampled datasets (dashed lines). Filtering out low-coverage peaks reduces false negatives across all tools (paired colors). TOBIAS scores remain largely unaffected under FRiP-downsampling due to the method's dependence on background depth.

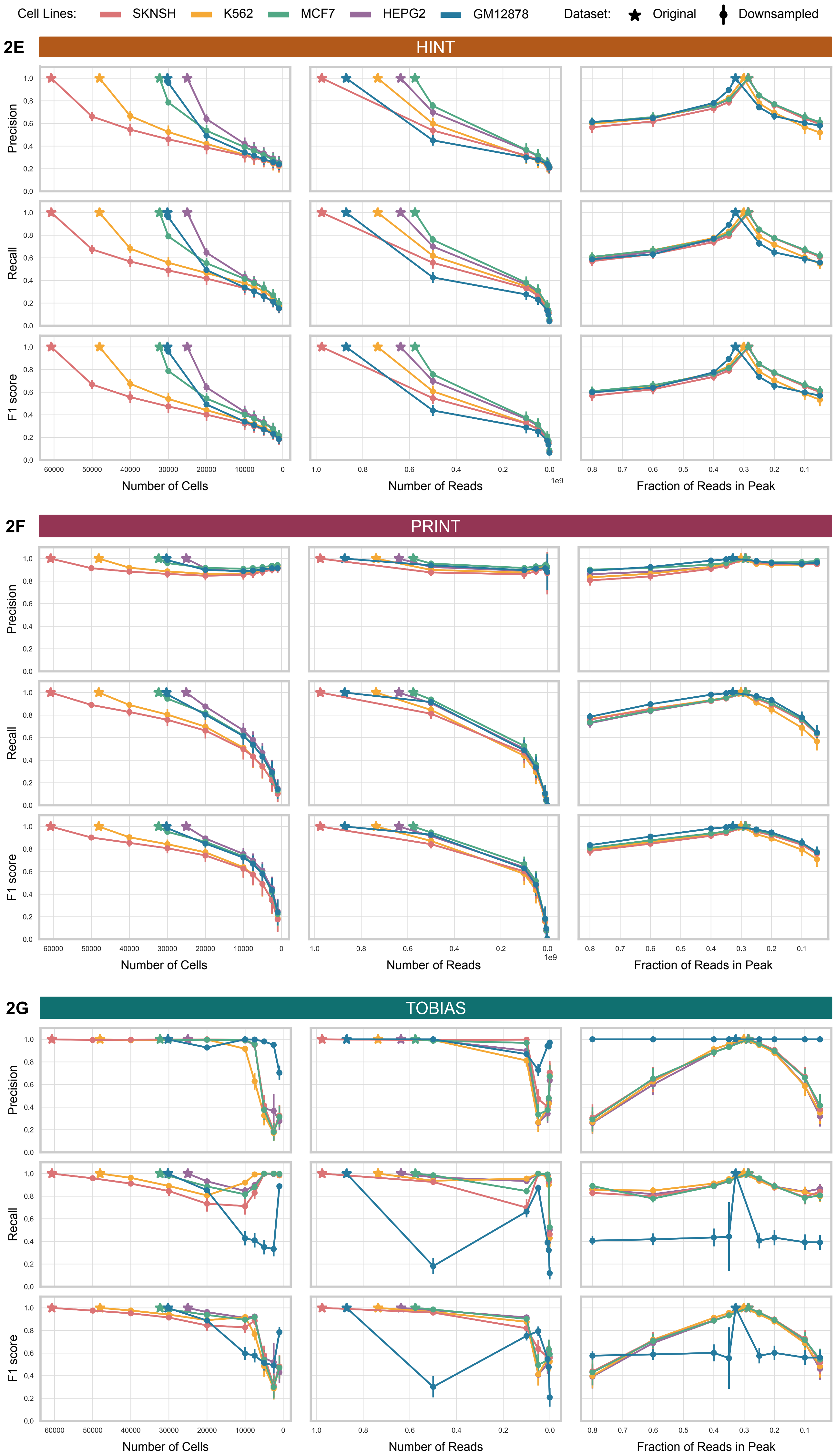

**Figure S2E-G) Expanded view of Figure 2A, showing separate plots for precision, recall, and F1 score for HINT (E), PRINT (F), and TOBIAS (G).**

**2H**

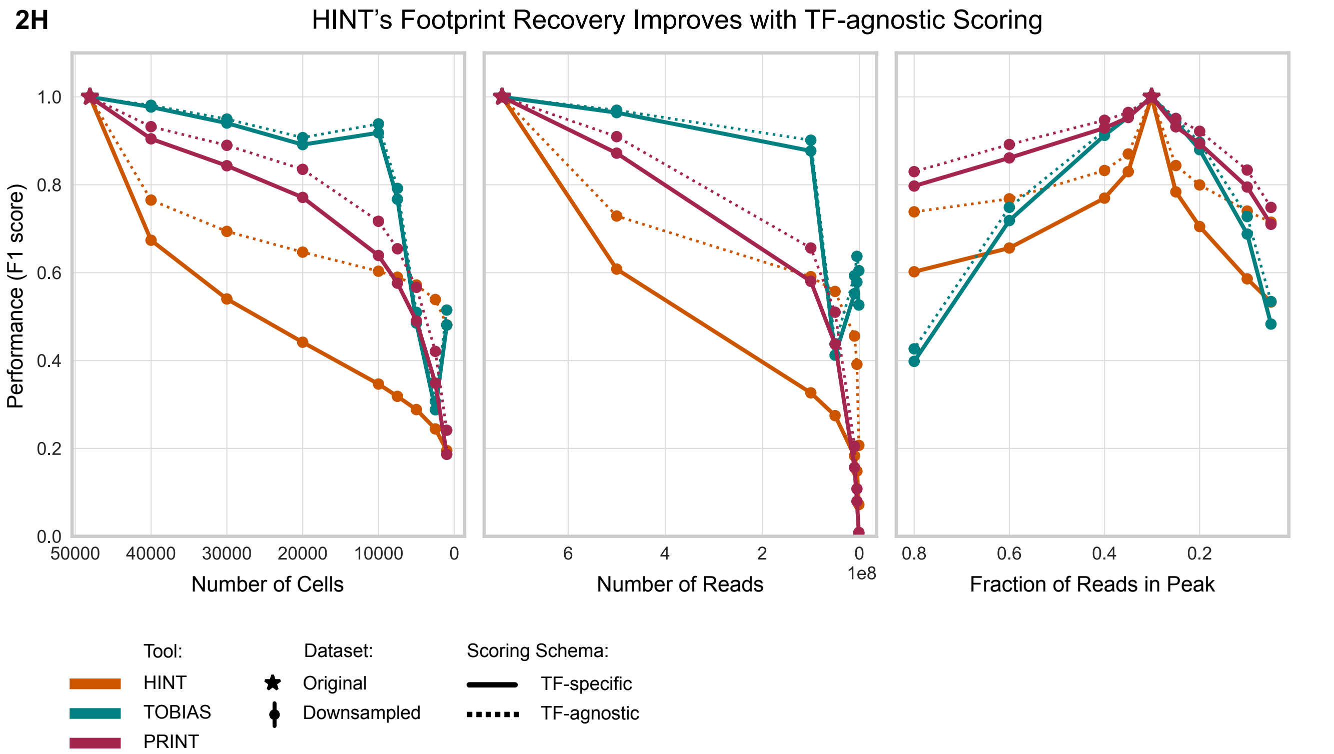

**Figure S2H) Slight variations in PWM matches partially explain HINT's reduced performance.** In our original comparison of footprint regions between original and downsampled datasets, a true positive required both footprints to fall within the same peak and share the same PWM match (solid line). When the PWM-match criterion is excluded (dashed line), HINT's performance (orange) improves substantially, while other tools are minimally affected. As a de novo method, HINT may identify slightly different footprint coordinates within the same peak. These subtle shifts can favor different PWMs, leading to output files that appear more dissimilar than they are. K562 is shown as an example; trends are consistent across all cell lines.

2I

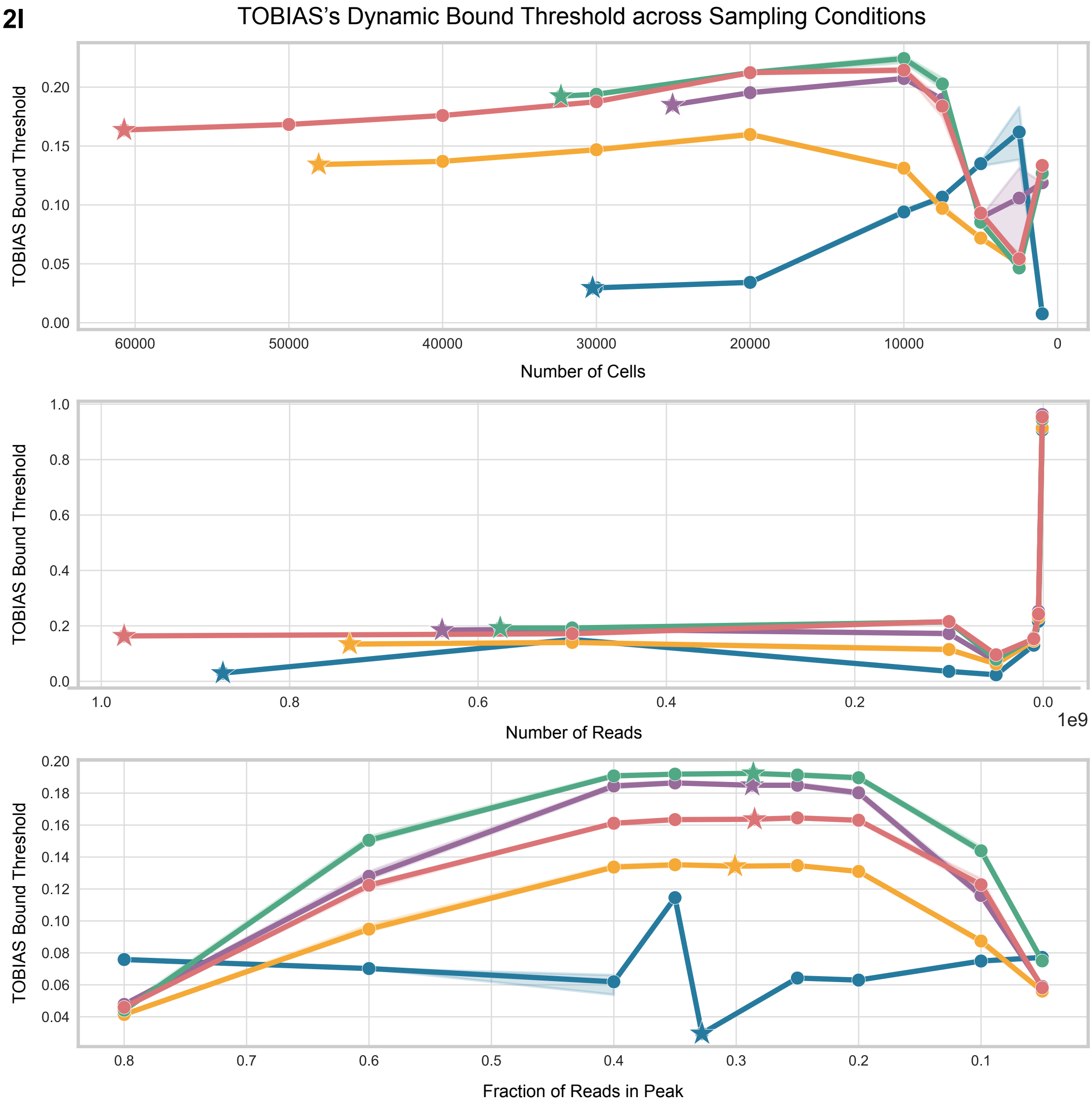

**Figure S2I) The bound threshold estimated by TOBIAS shows considerable variability across the progression of sampling conditions.** The threshold was extracted from the "BINDetect" log files. Points indicate mean ± SD across replicates (shaded fill).

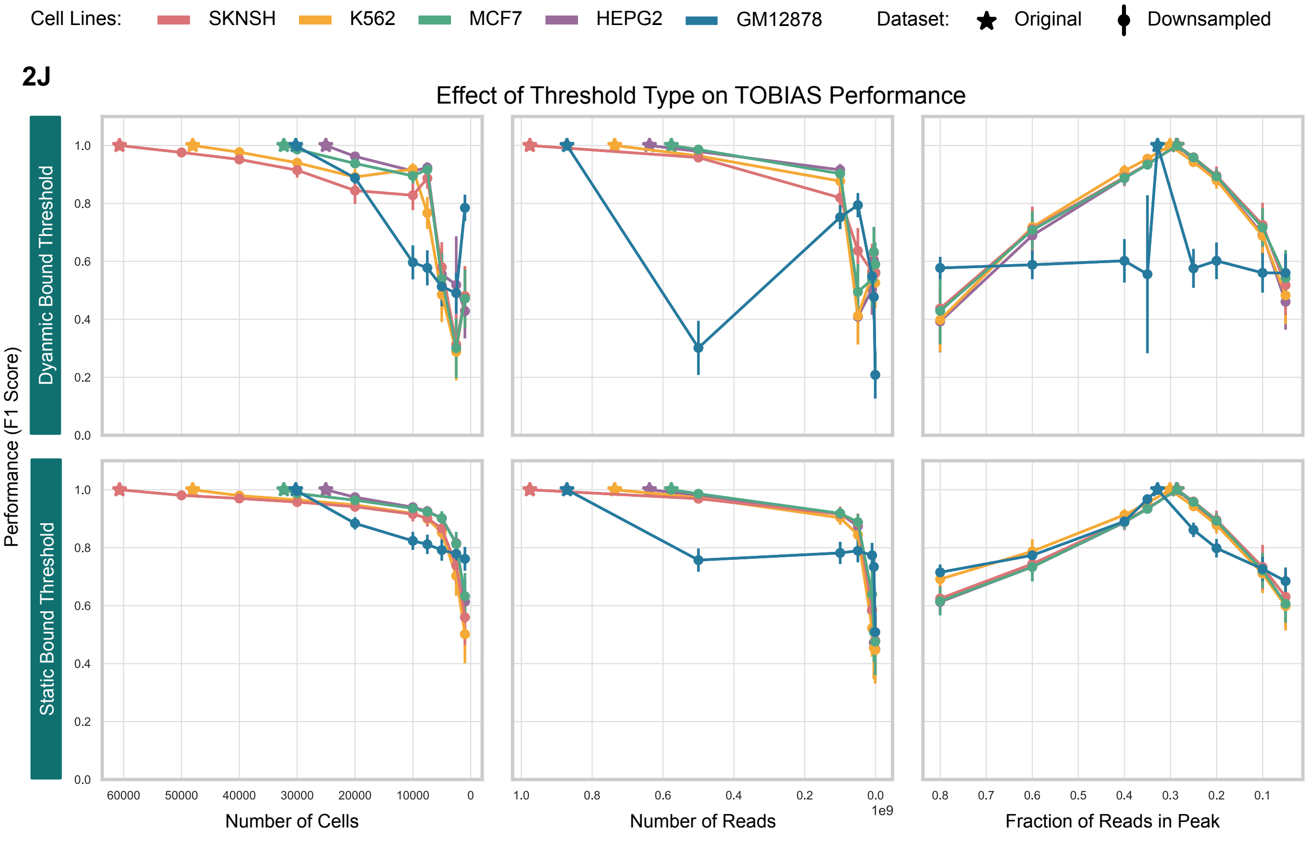

**Figure S2J) Using a static bound threshold with TOBIAS produces performance trajectories more similar to HINT and PRINT.** Line plots follow the same format as Figure 2A. The top row shows results using TOBIAS's dynamic bound threshold, which is computed separately for each input. The bottom row uses a static threshold, defined based on the original (non-downsampled) samples and corresponding to the star marker in Figure S2I.

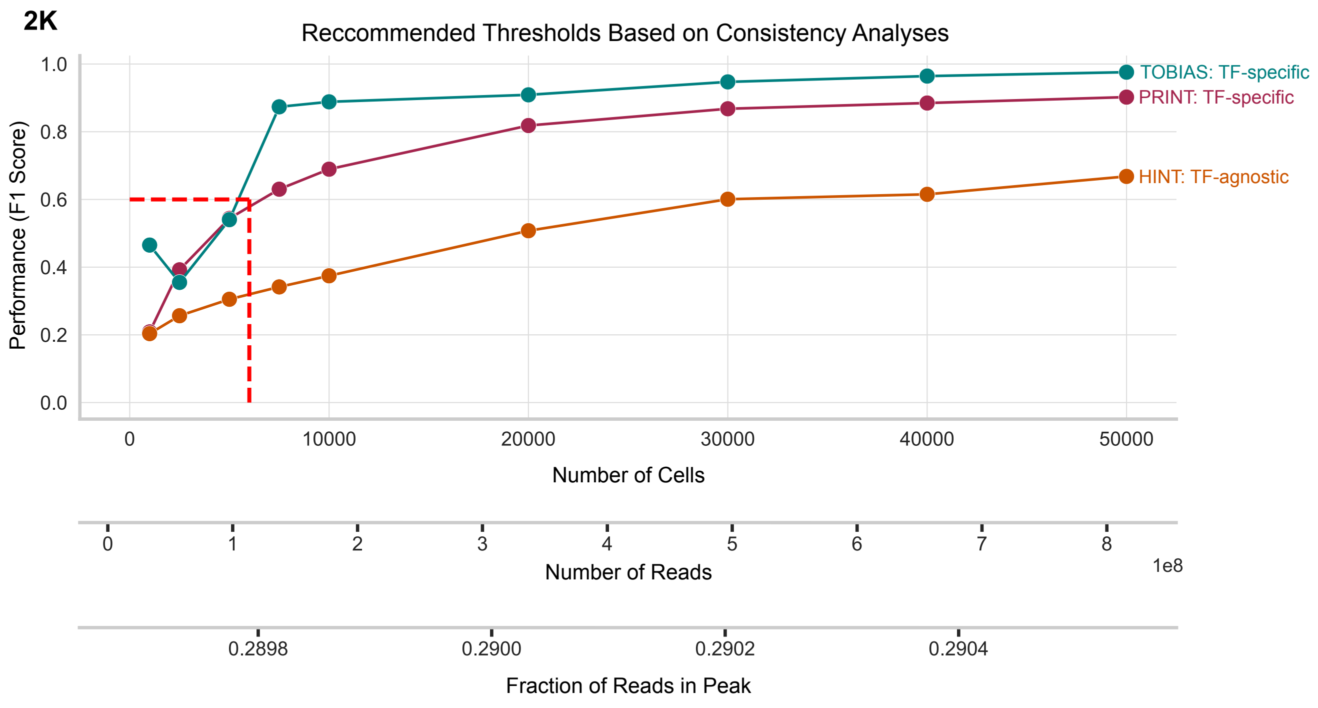

**Figure S2K) Recommended thresholds for achieving F1 scores ≥ 0.6, based on consistency analysis.** Line plots depict changes in performance (F1 score) across cell downsampling conditions. Each point represents the mean ± SD across cell lines. Secondary x-axes map additional data quality metrics (e.g., read count, FRiP) to the same downsampling trajectory. Note that the FRiP axis reflects the pseudobulk FRiP, which will be lower than the mean per-cell FRiP. Each line corresponds to a different tool; HINT is shown using TF-agnostic performance calculations. A dashed red line indicates the 0.6 F1 score threshold and its position along each x-axis, highlighting the minimum data quality needed for consistent performance.

**3A**

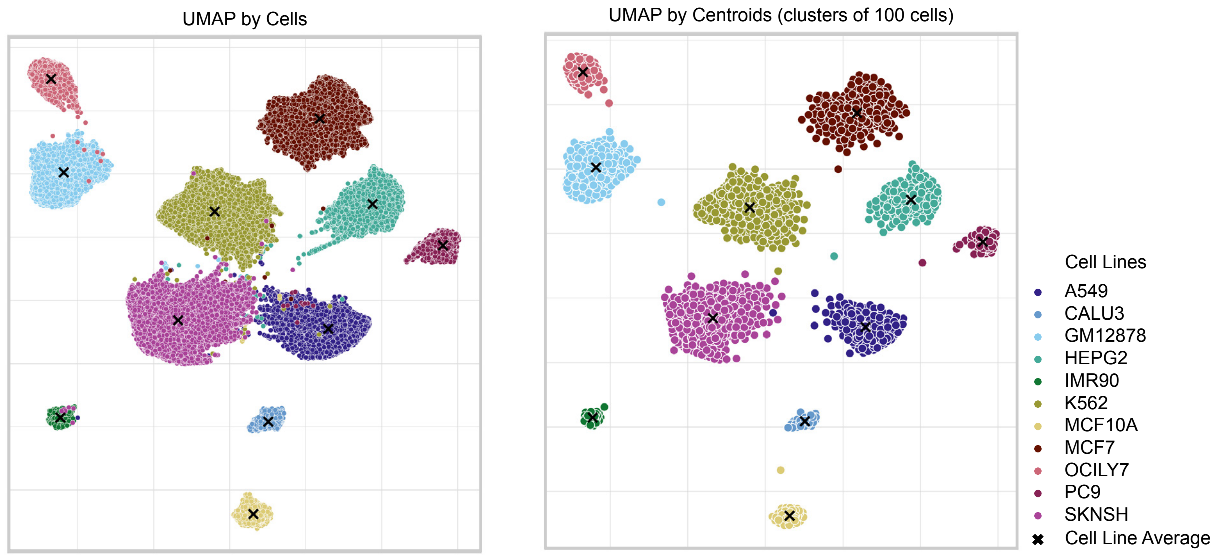

**Figure S3A) Constrained K-means clustering simplifies synthetic population construction.** UMAPs show individual cells (top) and clusters of 100 highly similar cells (bottom), generated using constrained K-means. This process reduces the computational burden when computing the Pearson Correlation matrices. Black Xs indicate the average position of each cell line. While clustering is shown in UMAP space for visualization, all metrics were computed in peak-count matrix space for comparability.

**3B**

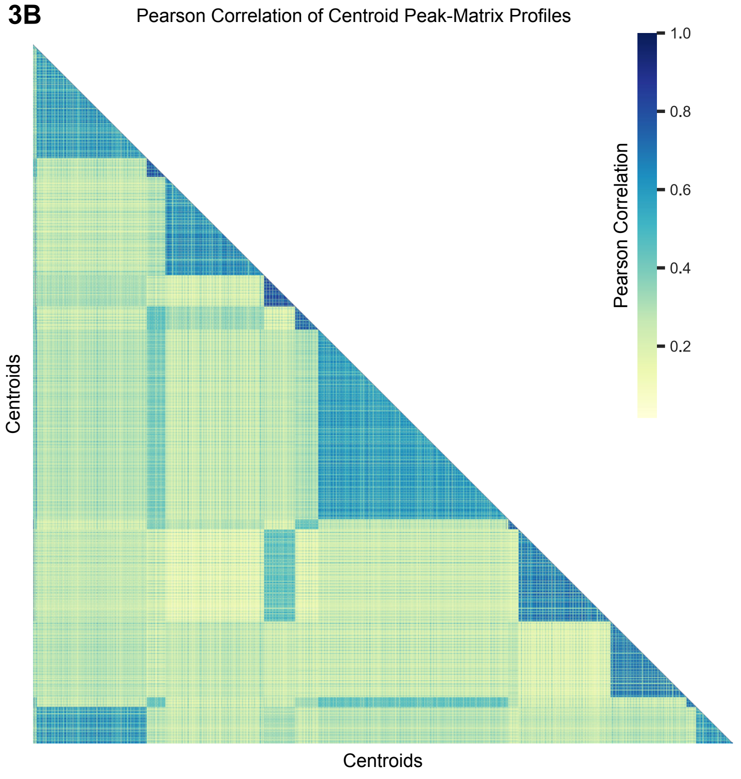

**Figure S3B) Pairwise Pearson correlation matrix between centroids.** After calculating the average peak-count profile for each centroid, pairwise Pearson correlation coefficients were computed. The heatmap shows the lower triangle of the resulting symmetric matrix. Darker diagonal triangles indicate centroids originating from the same cell line.

3C

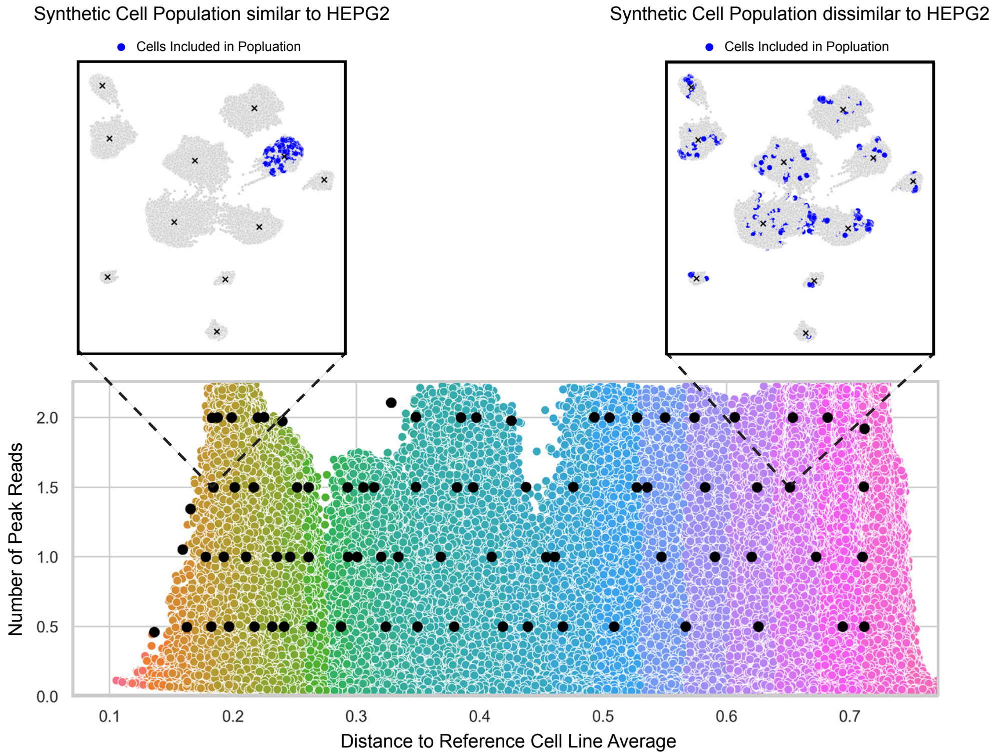

**Figure S3C) Selection of hypothetical cell populations by peak read depth.** Scatterplot shows ~300,000 synthetic pseudobulk populations (points), varying in total peak read depth and distance from the corresponding cell line centroid. To isolate the effects of cell homogeneity from read depth, we defined four target read-depth categories (5e7, 1e8, 1.5e8, and 2e8; dashed black lines) and selected ~20 populations per category, evenly spaced along the x-axis (solid black dots). Two example populations are shown in the UMAPs above, with contributing cells highlighted in blue.

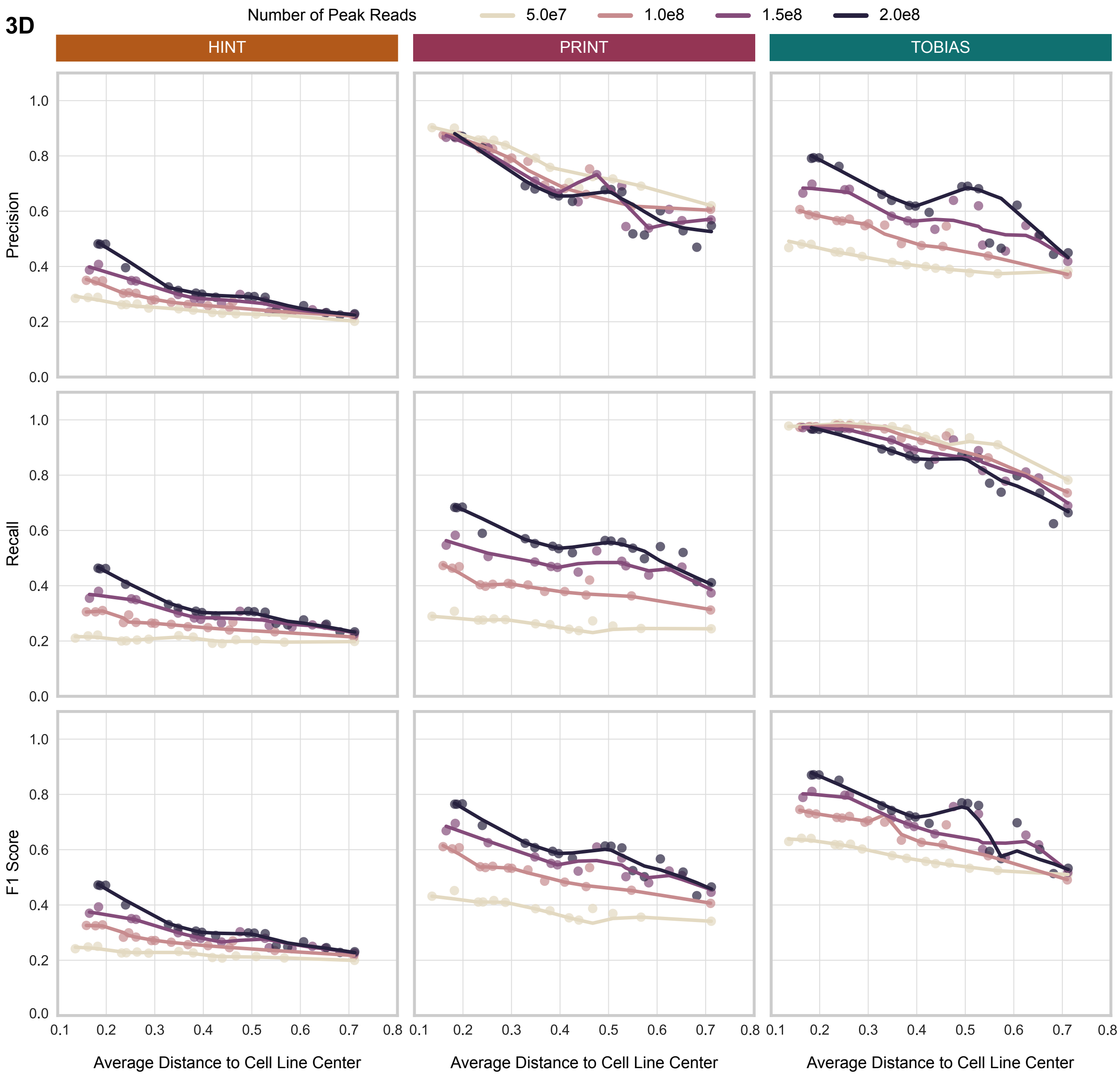

**Figure S3D) Expanded view of Figure 3A, showing separate plots for precision, recall, and F1 score.** Tools are arranged as columns, with performance metrics as rows. The third row (F1 Score) corresponds to the left column of Figure 3A. HINT shows balanced precision and recall, while TOBIAS has higher recall and PRINT shows higher precision.

4A

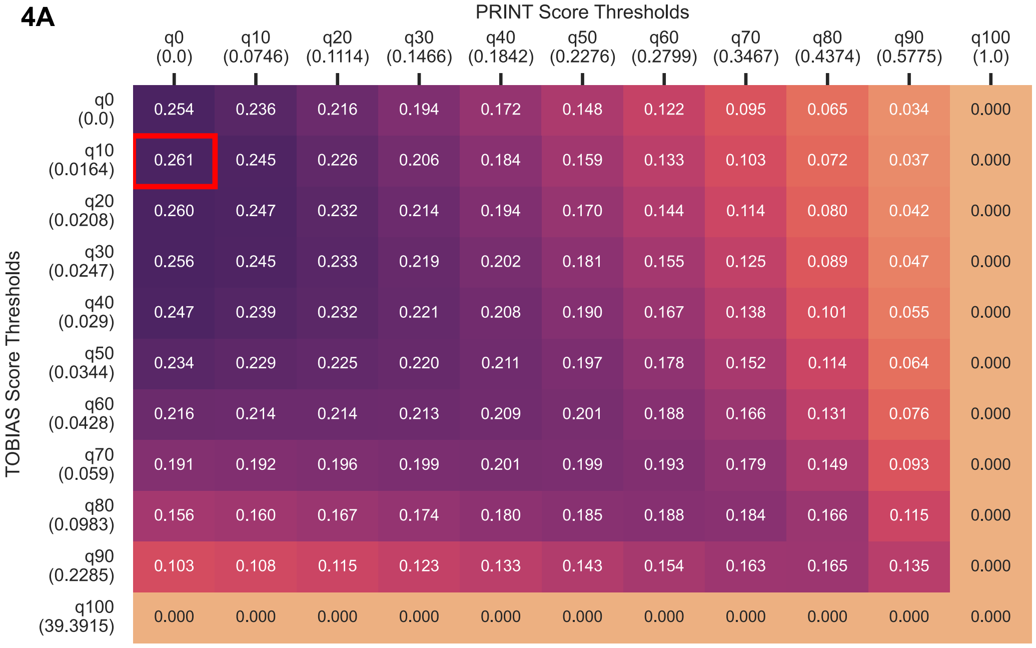

**Figure S4A-C) Low overlap between tools is not explained by bound-threshold selection.** Heatmaps show Jaccard coefficients (measure of overlap) of footprint regions between tool pairs. For each tool, thresholds were defined at every 10th quantile of the score distribution (quantile values shown in parentheses), and footprints above each threshold were compared. The maximum overlap across all threshold combinations is indicated by a red box. Results shown are based on the baseline HepG2 dataset across (A) PRINT and TOBIAS, (B) PRINT and HINT, and (C) TOBIAS and HINT.

4B

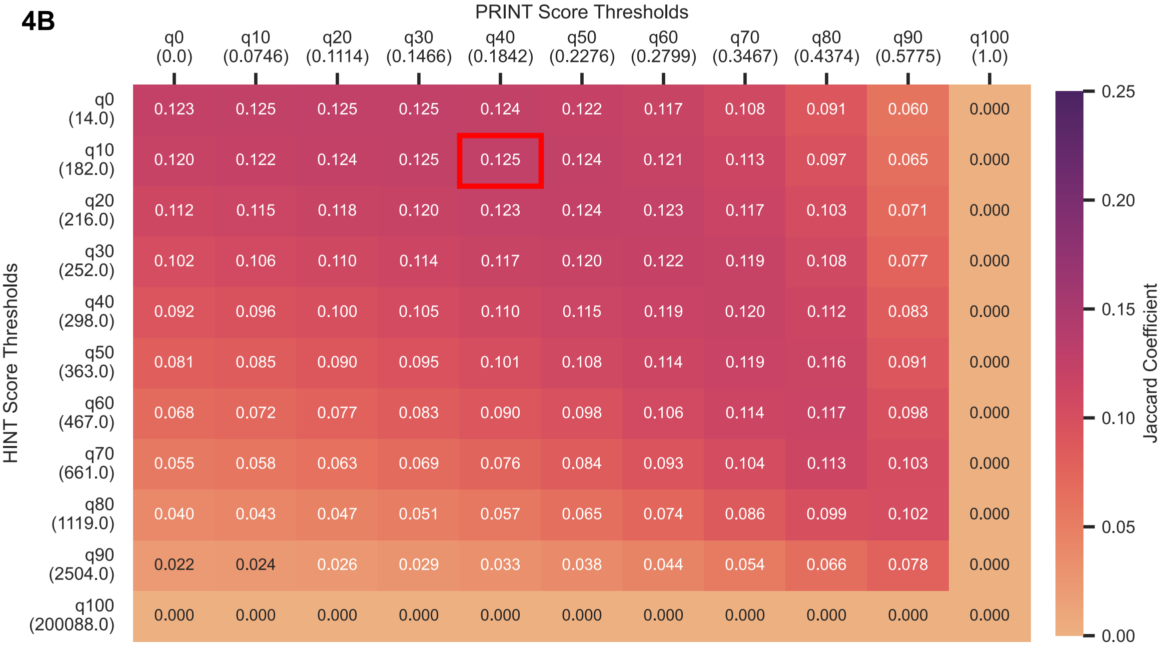

4C

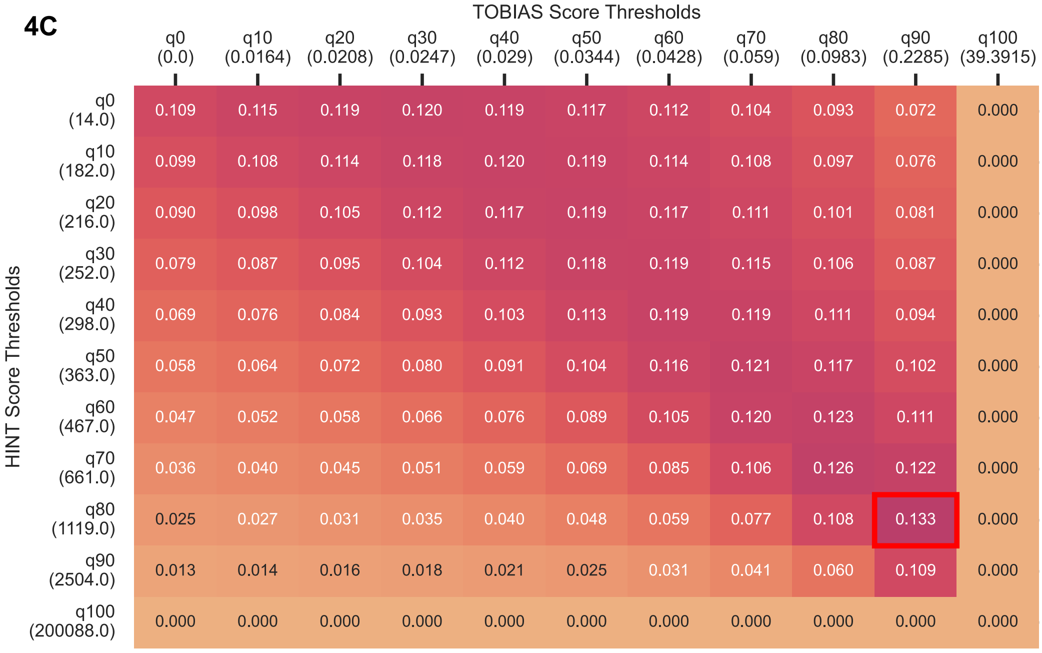

4D

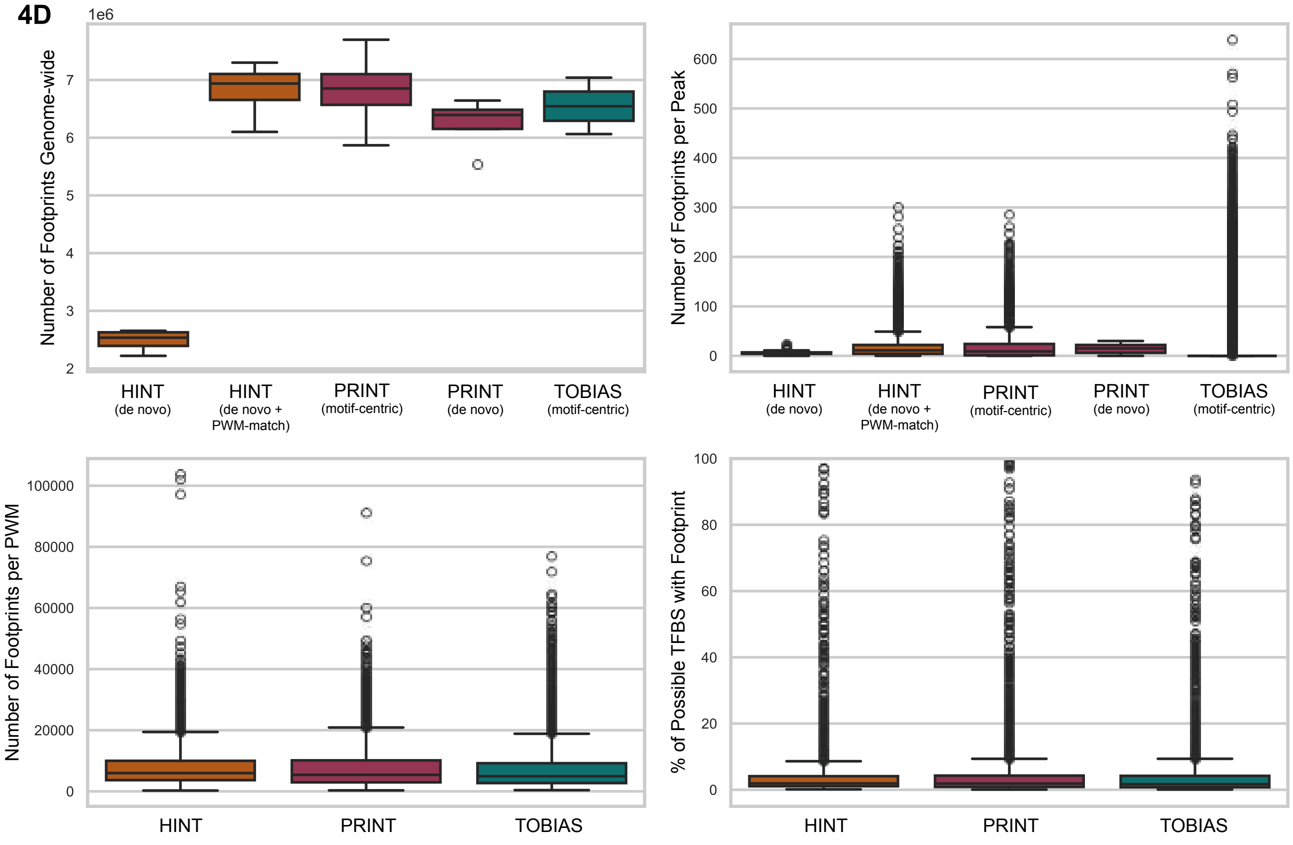

**Figure S4D) Genome-wide footprinting statistics are broadly consistent across tools.** On average, each method identified 6 million footprints, or roughly 15 footprints per OCR and about 8,000 footprints per PWM. Given the large number of potential TFBSs within OCRs for each PWM, this corresponds to an average footprinting rate of ~6%. Boxplots summarize results across four cell lines (HepG2, K562, MCF-7, SK-N-SH) for each method. The top row—number of footprints genome-wide and number of footprints per peak—represents PWM-independent metrics, allowing direct comparison of the footprinting steps across tools. For HINT, both the initial de novo footprinting and the subsequent PWM-matching step are shown. PRINT includes its motif-centric analysis and the initial de novo step. TOBIAS is represented solely by its motif-centric approach. The bottom row—number of footprints per PWM and percent of TFBS occupied by a footprint—reflects motif-specific results based on the full JASPAR database (n = 841 PWMs).

4E

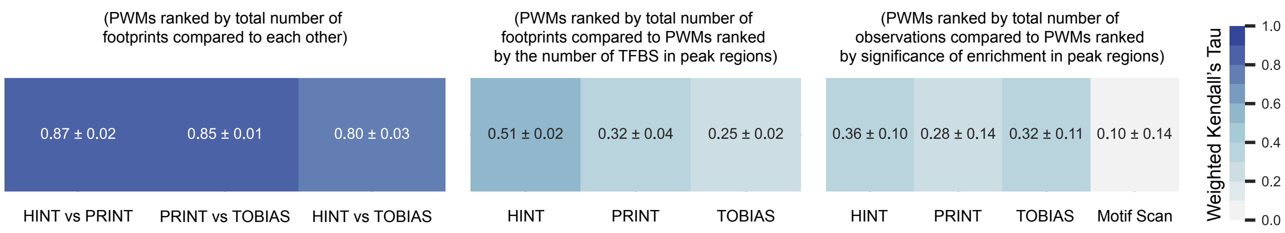

**Figure S4E) Expanded view of Figure 4C including motif enrichment analysis.** Motif enrichment of peak regions was performed using HOMER with a GC-matched background. Motif scanning to identify all potential TFBSs within peaks was carried out using the R package monaLisa, applying a score threshold of 6. Heatmaps display the similarity in PWM rankings across methods, quantified using weighted Kendall's tau (mean ± SD). Rank 1 was assigned to the most frequently observed PWM for each method. Tile color and text indicate the correlation values.

**5A**

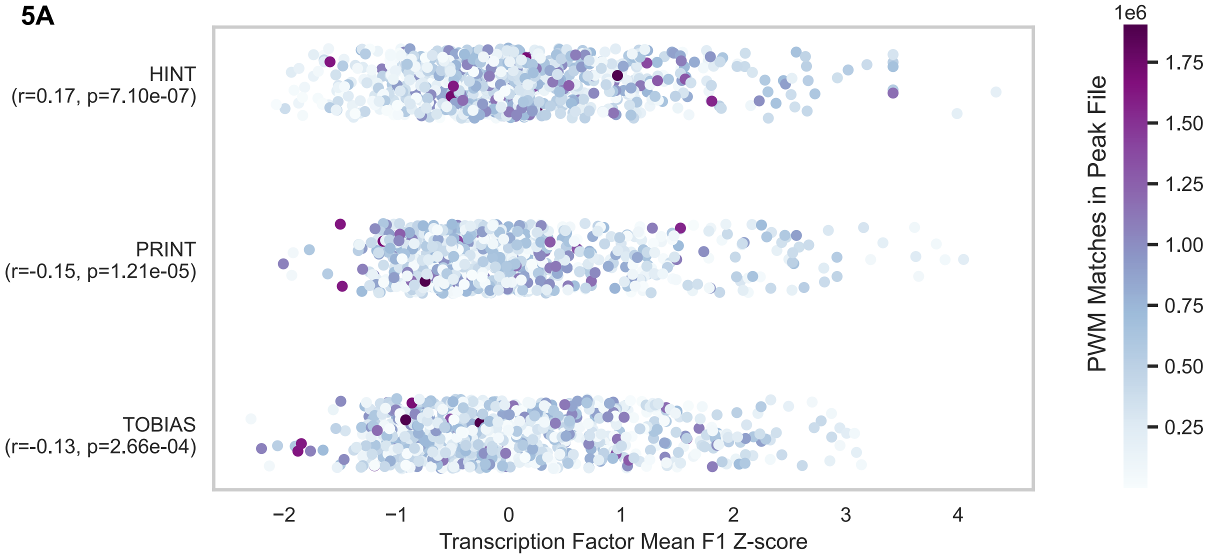

**5B**

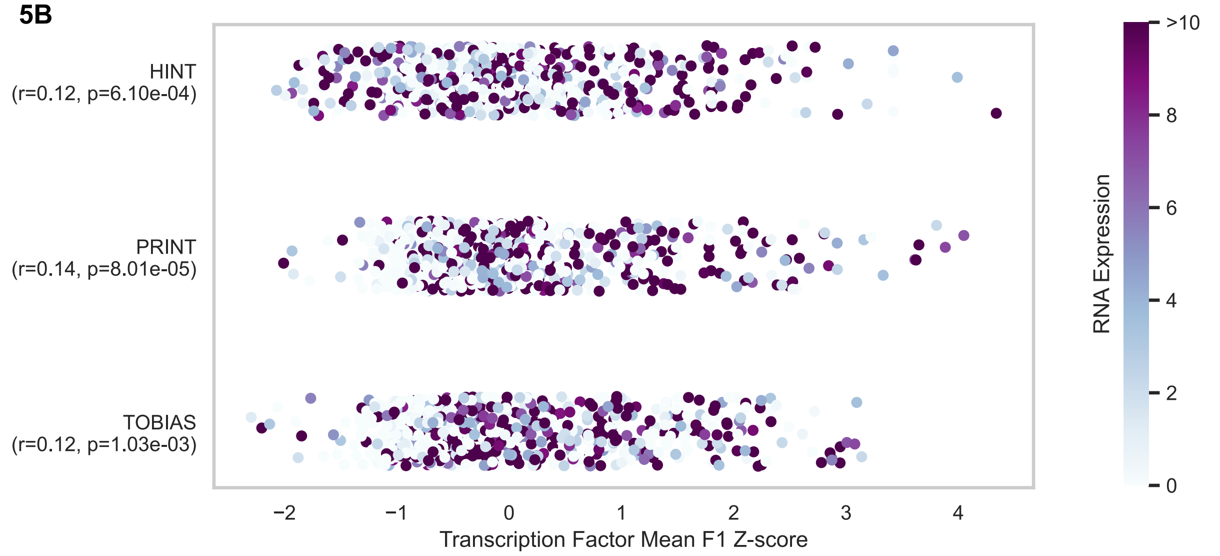

**Figure S5A–B) TF performance is not correlated with TFBS prevalence or RNA expression.** Similar to Figure 5A, strip plots show that neither (A) the number of PWM matches in the peak file nor (B) RNA expression levels are significantly correlated with TF Z-scores across downsampling conditions. HepG2 is shown for illustration.

5C

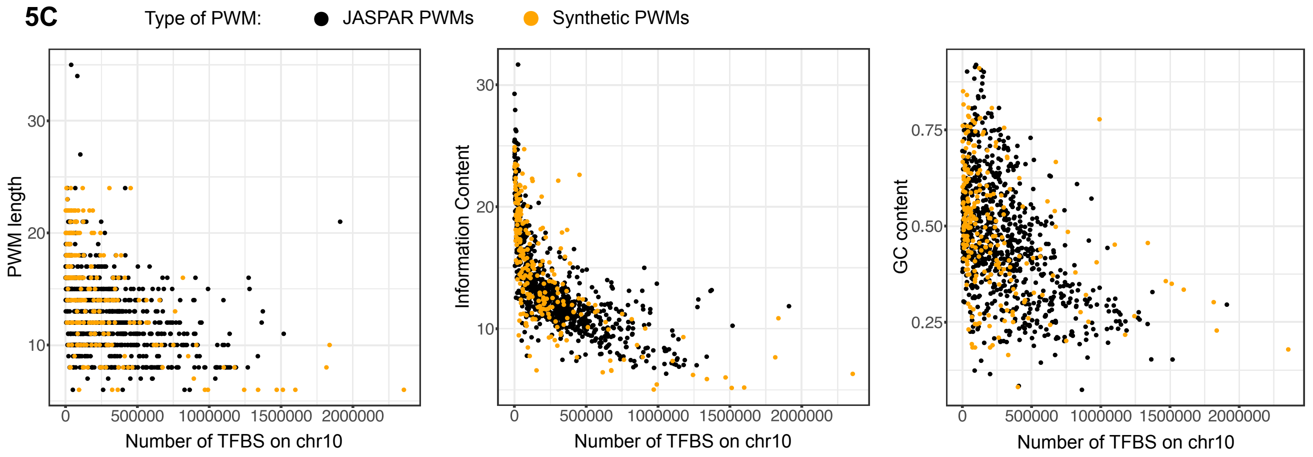

5D

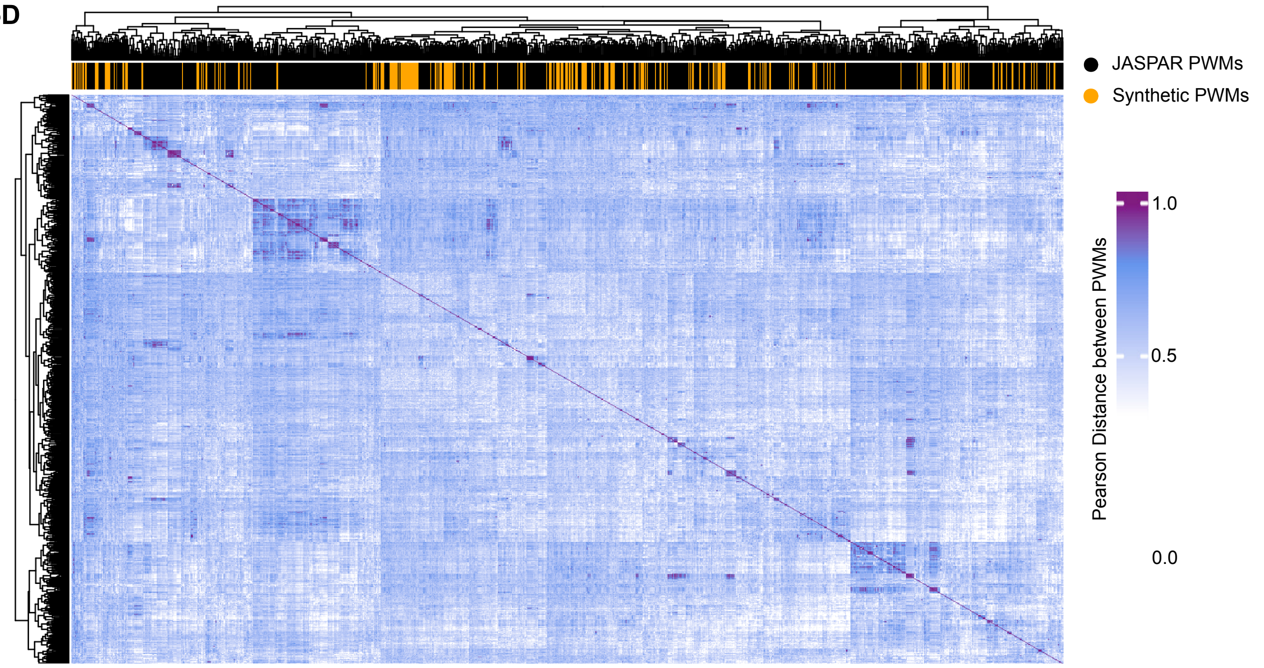

**Figure S5C) Synthetic PWMs closely resemble real JASPAR motifs.** Synthetic PWMs (orange) match JASPAR motifs (black) in length, information content, GC content, and genome-wide frequency on chromosome 10.
**Figure S5D) Synthetic PWMs cluster alongside real JASPAR motifs.** Heatmap shows the pairwise Pearson correlation distance matrix between synthetic (orange) and real JASPAR (black) PWM matrices. Rows and columns represent the full set of motifs, with PWM type annotated on the columns. Both axes were hierarchically clustered using Euclidean distance and complete linkage. Synthetic motifs are intermixed with real motifs and do not cluster separately.

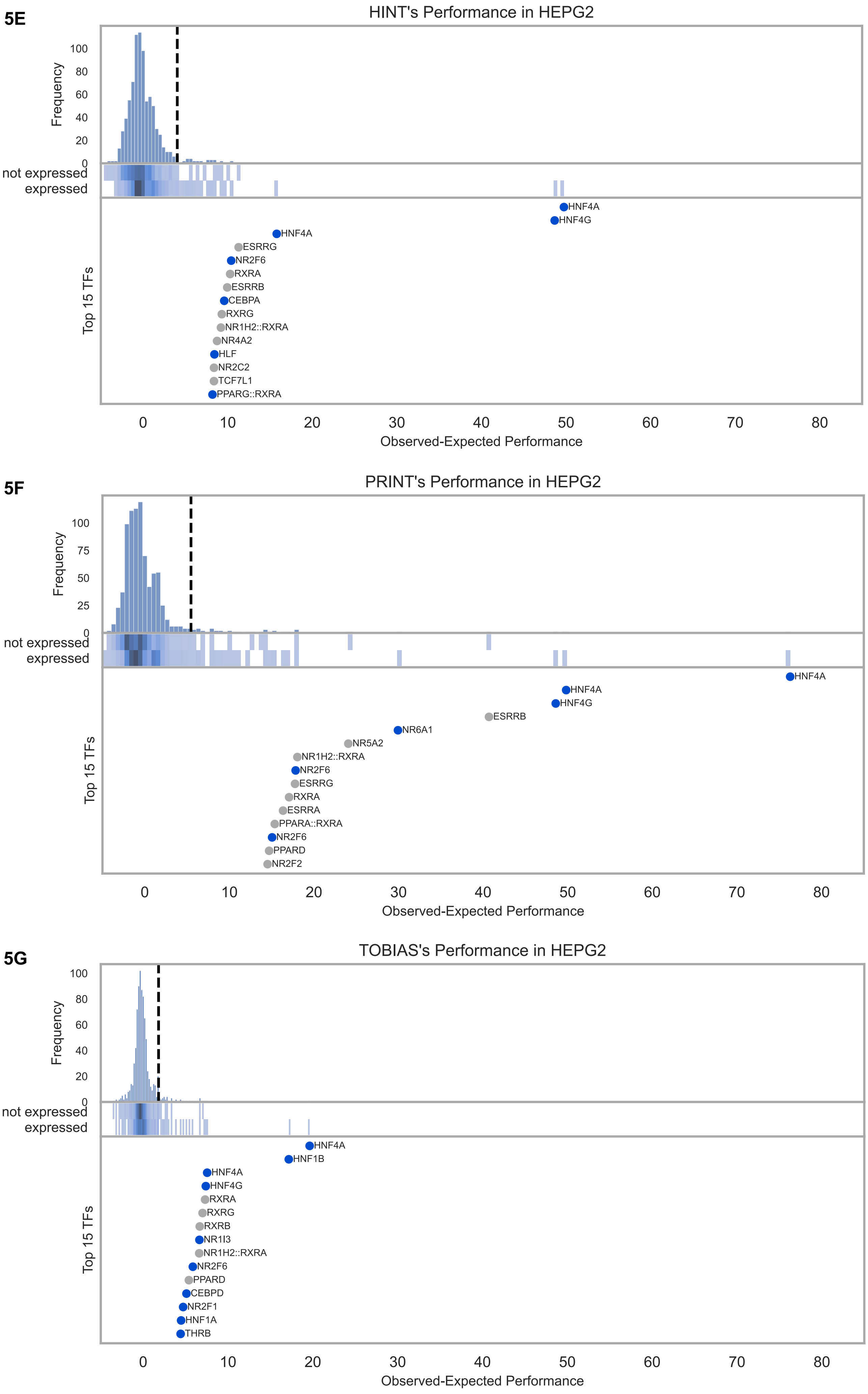

**Figure S5E–G) All tools recover HepG2-specific transcription factors.** Using z-score normalized footprint counts, each tool ranks TFs based on their enrichment in HepG2 relative to the average across other cell lines. Known liver-enriched TFs such as HNF4A are consistently prioritized. The top panel shows the distribution of observed-minus-expected values, with the 95th percentile marked by a vertical black line—corresponding to the threshold used in the UpSet plots in Figure 5C–D. The middle panel presents horizontal density plots indicating where TFs with detectable RNA expression (TPM > 1) fall within the distribution. The bottom panel labels the top 15 ranked TFs, with those enriched in HepG2 RNA-seq highlighted in blue. These rankings correspond to the heatmaps in Figure 5E–F.

**5H**

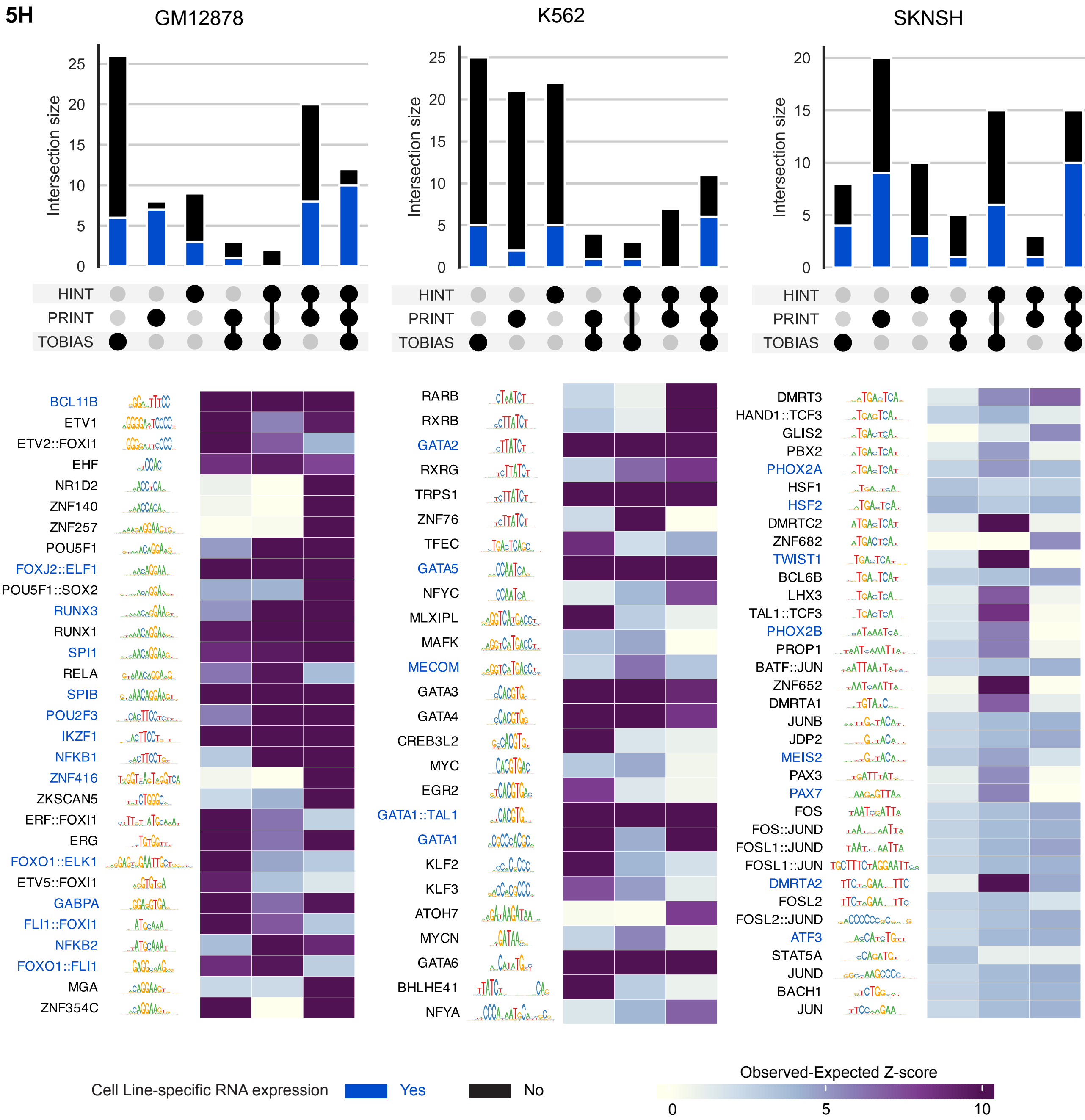

**Figure S5H) Expanded view of Figure 5D-F for additional cell lines.** Heatmaps and UpSet plots, following the same format as Figure 5D–F, are shown for GM12878, K562, and SK-N-SH. Rows remain ordered by hierarchical clustering, though dendrograms are omitted for clarity and space.

**5I**

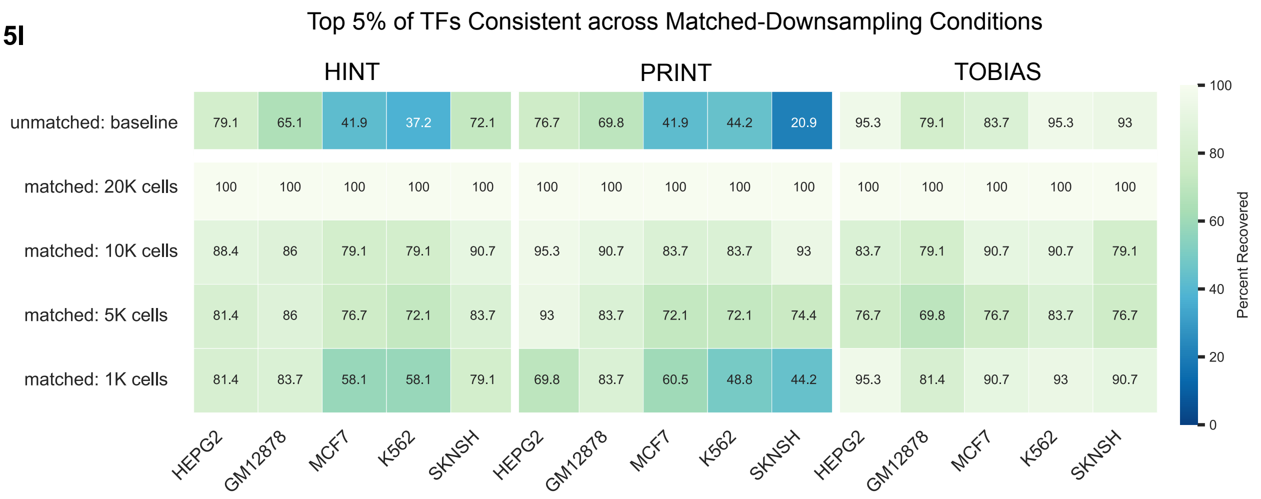

**Figure S5I) Top-ranked TFs remain stable across cell downsampling conditions.** To assess robustness, we tested whether the same TFs appeared in the top 5% of footprint-enriched motifs as cell number decreased. Each cell is colored and labeled by the percentage of TFs recovered relative to the 20,000-cell reference (e.g., 100% indicates full recovery). Across matched downsampling conditions, results were highly consistent, often recovering > 80% of TFs. The largest drop occurred when comparing to the original baseline samples, which had variable starting cell numbers—underscoring the importance of matched data quality.

**6A**

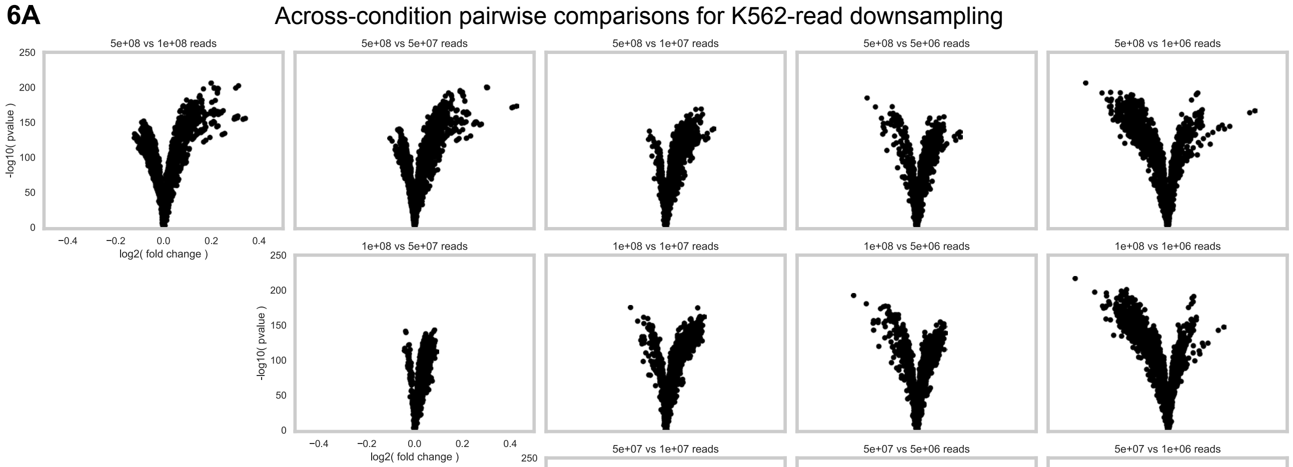

**Figure S6A) Read depth differences drive variation in significance and effect size.** Volcano plots show pairwise comparisons between K562 read-downsampling conditions using TOBIAS, as described in Figure 6B. Panels are ordered from highest to lowest read depth. The consistent leftward shift reflects motifs enriched in the higher read-depth sample.

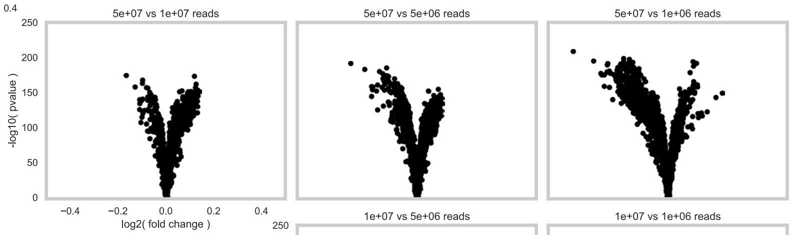

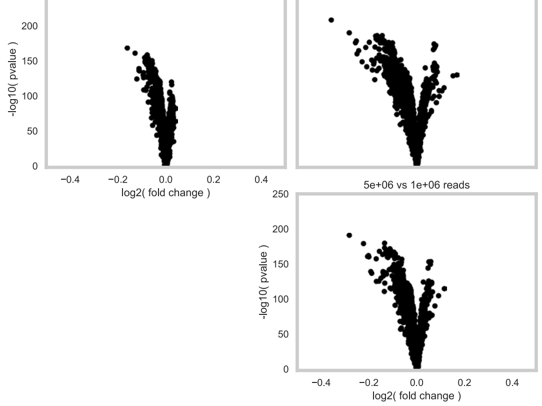

**Figure S6B) Expanded view of Figure 6A with additional sampling types.** Layered volcano plots showing within-condition pairwise comparisons. Comparisons across downsampling conditions are colored; a comparison between distinct cell lines (black) is included for scale reference.

**Figure S6C) Expanded view of Figure 6B with additional sampling types.** Layered volcano plots showing all across-condition pairwise comparisons based on FRiP scores. Larger fold-changes are observed in FRiP-based downsampling, consistent with expectations, though variation due to read and cell depth should be minimized.

**6B**

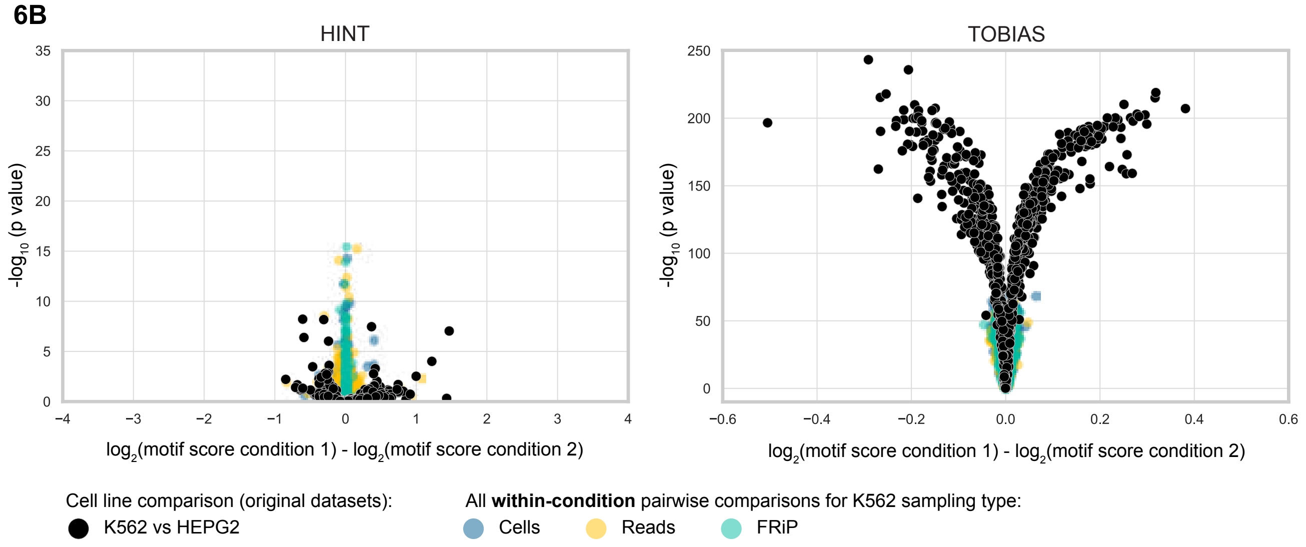

**6C**

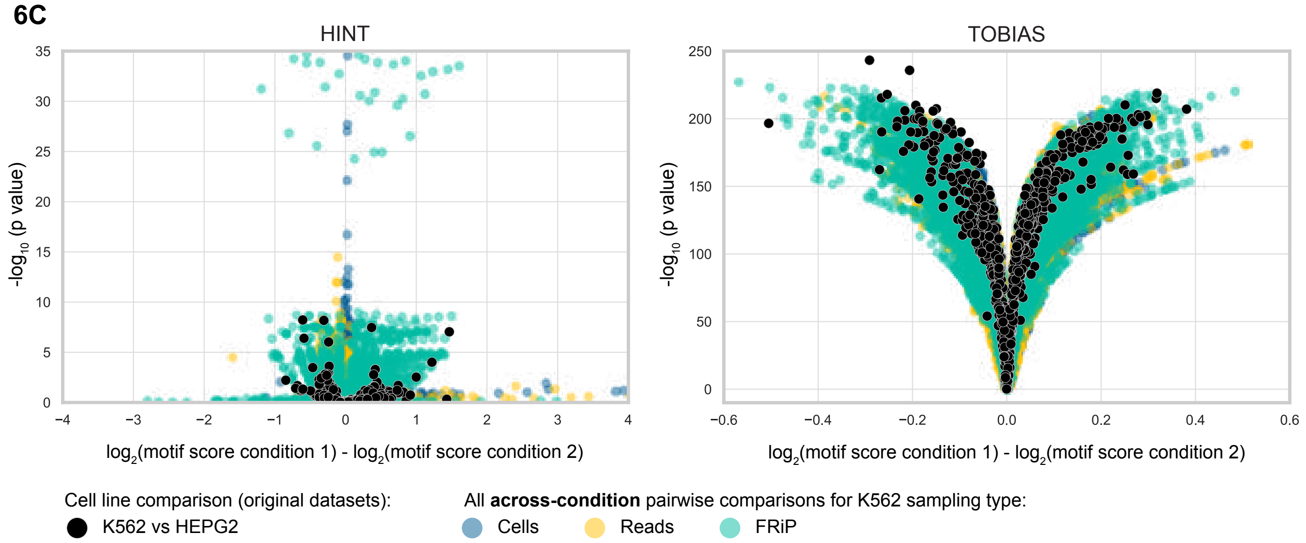

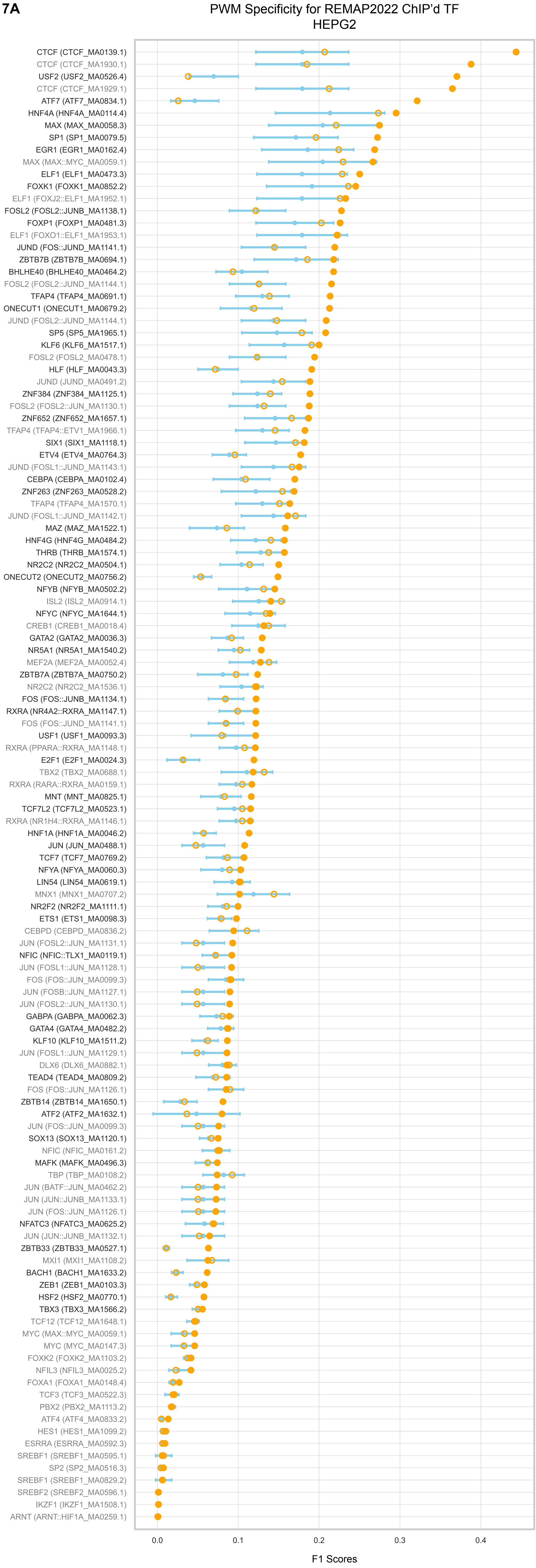

**Figure S7A) PWMs display varying specificity for ChIP'd TF regions.** To assess PWM-TF pair performance, we plotted F1 scores (x-axis) for each TF-PWM pair (y-axis), comparing (1) mismatched PWMs (light blue), (2) the true PWM with shuffled counts (orange no fill), and (3) the true PWM (orange). Only pairs with recall > 0.4 are shown. Pairs were retained if the true PWM outperformed both controls and had an F1 score > 0.5. Final pairs (black) were used in further analyses; excluded TFs are shown in grey.

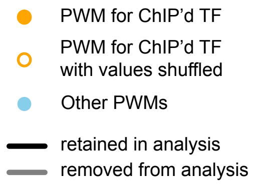

**7B**

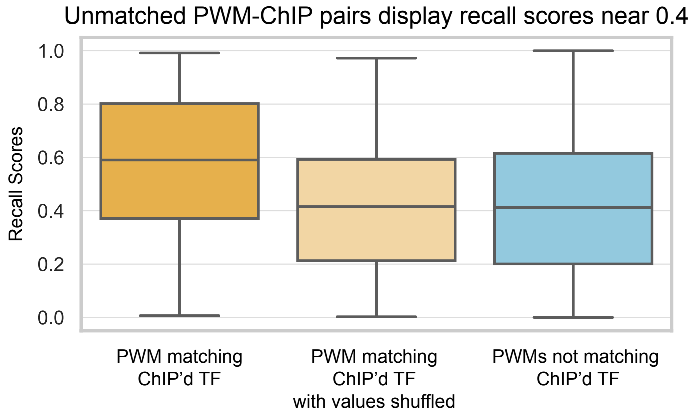

**Figure S7B) PWMs not matching ChIP'd TF display recall scores near 0.4.** To estimate recall levels for random or mismatched PWMs in ChIP experiments, we calculated recall scores for: (1) the correct TF PWM, (2) the TF PWM with shuffled regions, and (3) non-matching PWMs. Box plots show that shuffled and mismatched PWMs typically have recall scores near 0.4. PWM-ChIP pairs below this threshold were excluded. Results are shown for HepG2, K562, MCF-7, and SK-N-SH.

**7C**

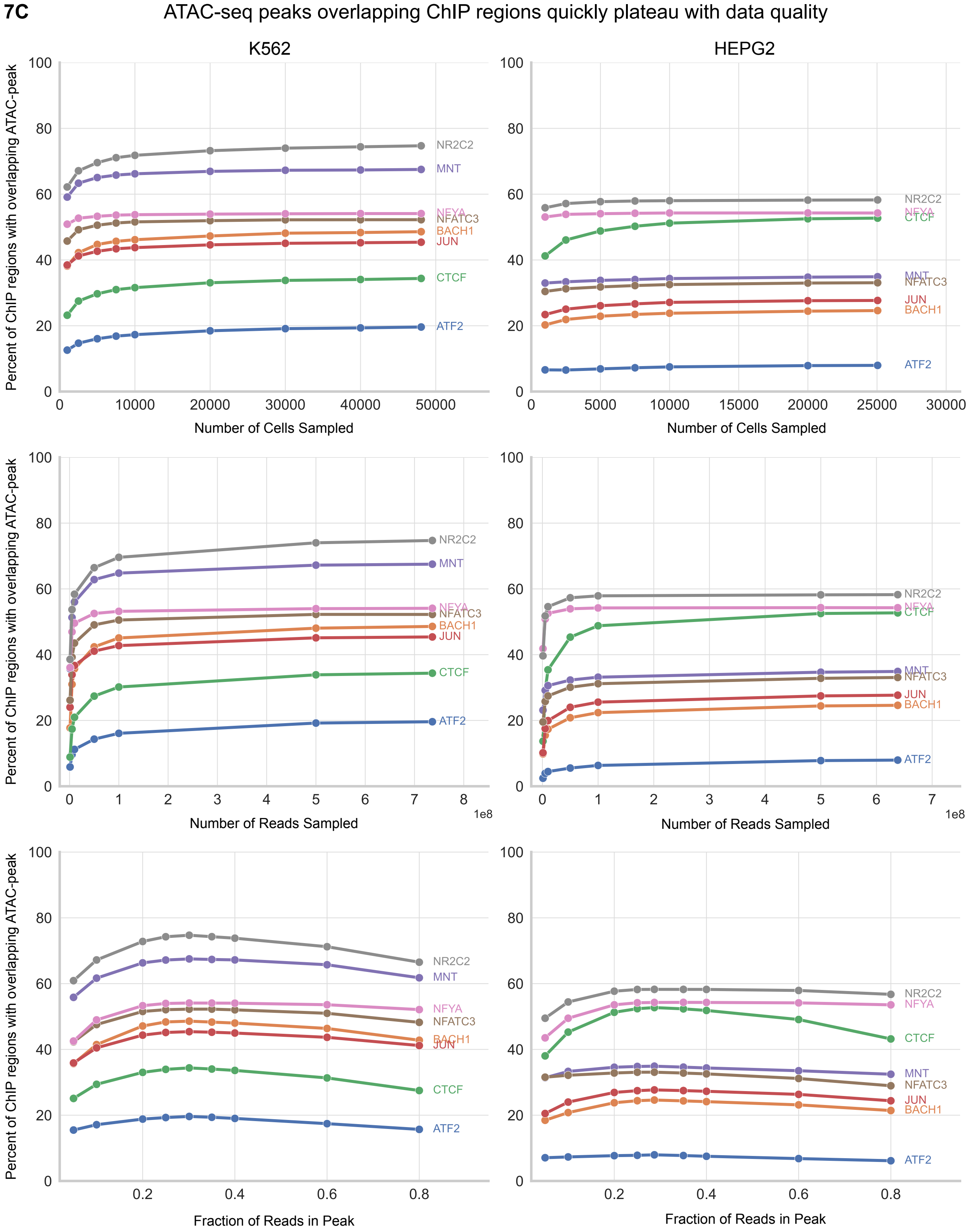

**Figure S7C) Overlap between ATAC-seq peaks and ChIP regions plateaus with reduced data quality.** Line plots show the percentage of annotated ChIP regions that intersect with scATAC-seq peaks across downsampling conditions (rows). Peaks were called using identical commands per BAM file. Each line represents one of eight randomly selected transcription factors (TFs). Columns show two different cell lines, illustrating how relative TF rankings can shift by context.

**7D**

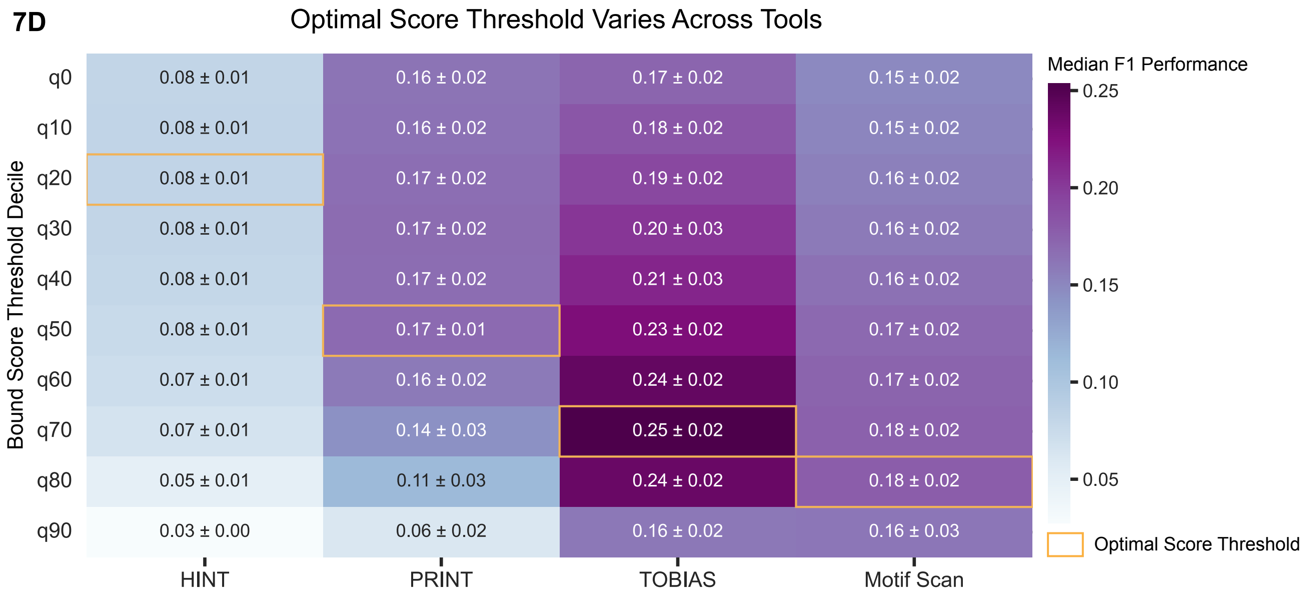

**Figure S7D) Expanded view of Figure 7A including all cell lines.** As in Figure 7A, but here the cell color and annotation reflect the median and standard deviation of F1 scores across all TF–cell line combinations.

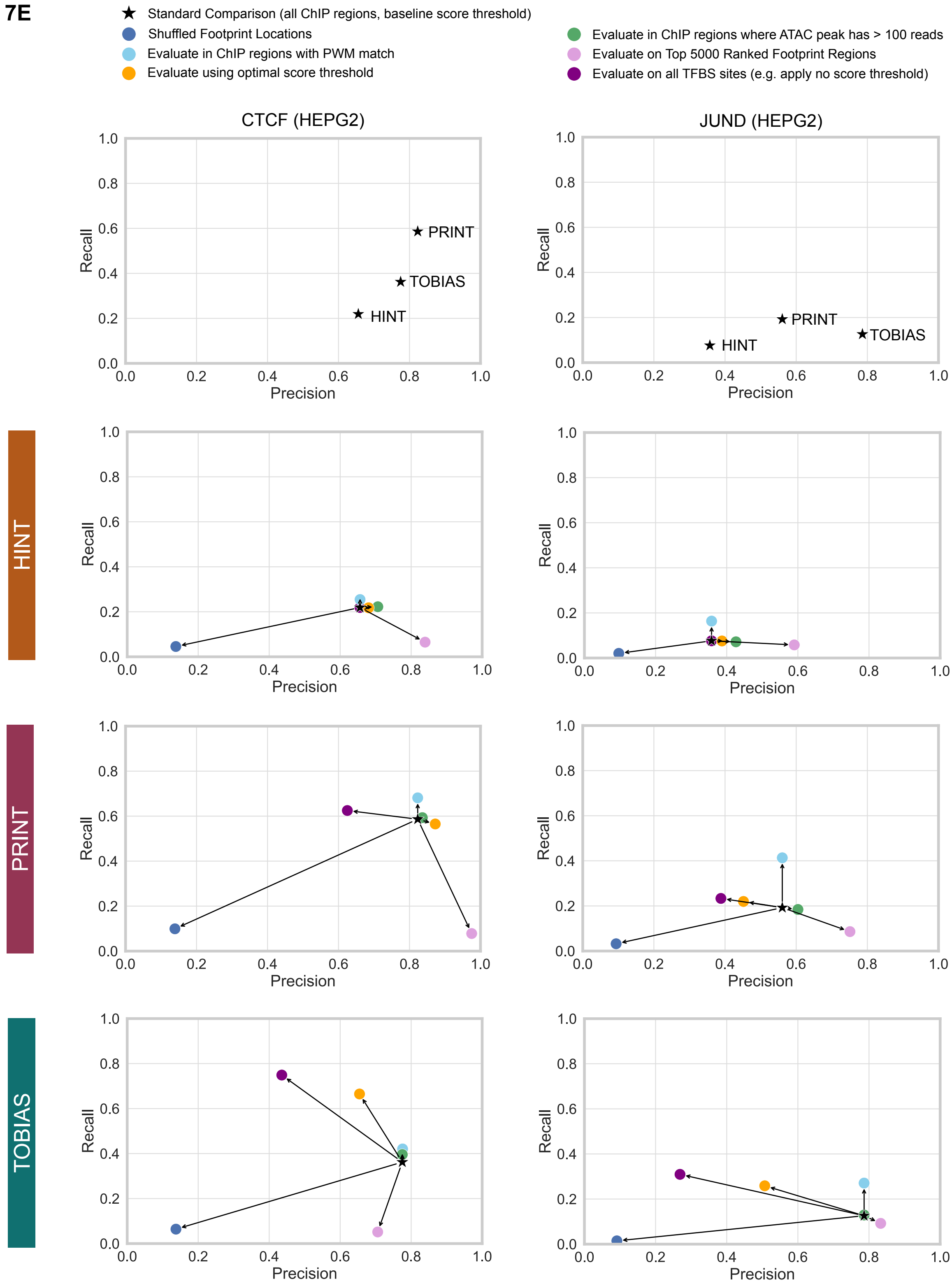

**Figure S7E) Evaluation strategies for comparing footprints to ChIP-seq data affects precision and recall in distinct ways.** Colored dots represent the outcomes of alternative approaches, shown relative to the standard comparison (black star). For example, shuffling footprint positions (dark blue) reduces both precision and recall, while restricting analysis to the top 5,000 highest-scoring footprints (light purple) increases precision but lowers recall. Results are shown for two TFs in HepG2, CTCF and JUND, displayed in columns. The top row shows only the standard comparison for clarity; subsequent rows correspond to individual tools.

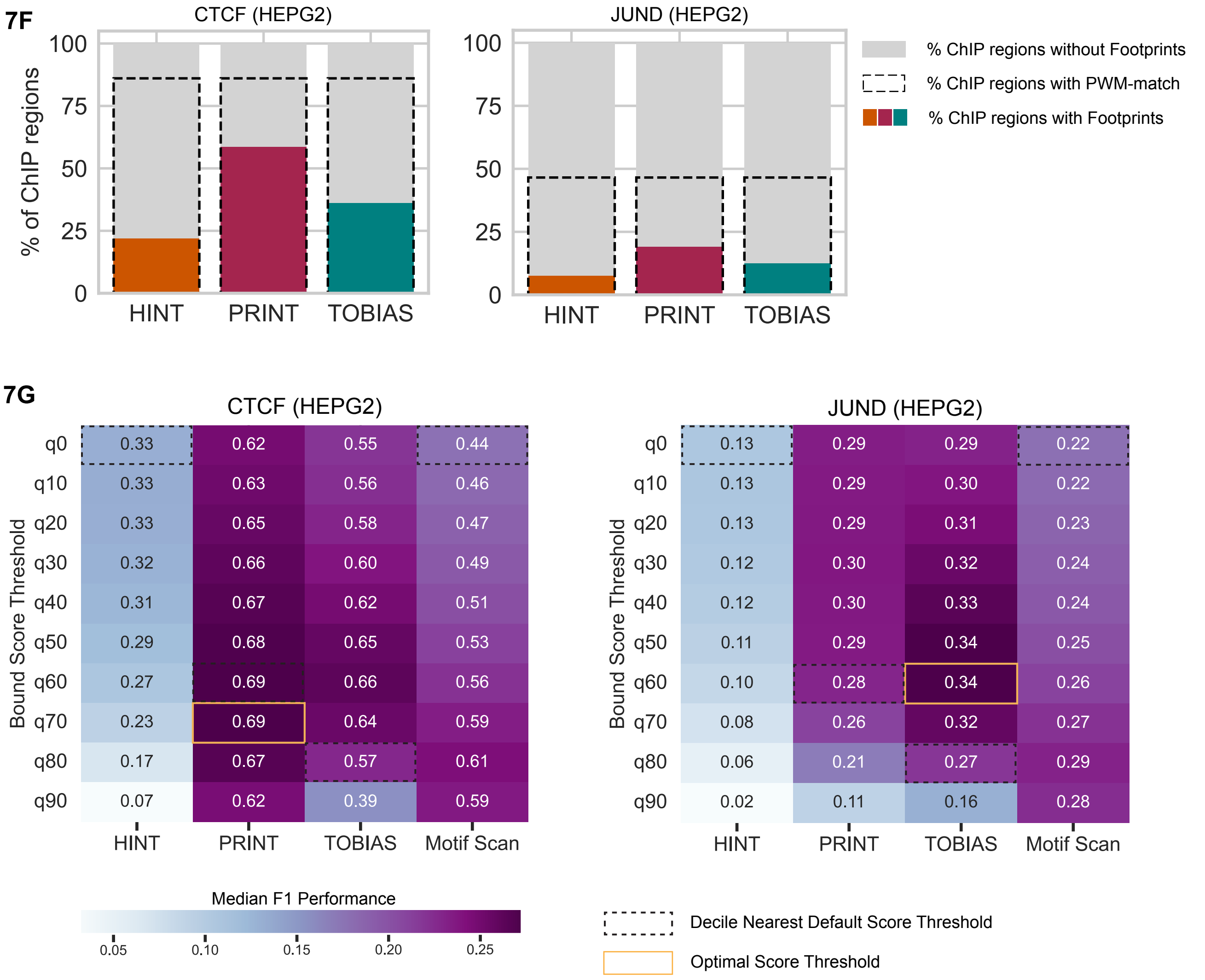

**Figure S7F) Percentage of ChIP regions containing the expected PWM varies across TFs.** Bar plots show the proportion of annotated ChIP regions that contain the expected PWM sequence, illustrating baseline variability across TFs. For example, 86% of CTCF ChIP-seq peaks contain a motif (left), compared to only 46% for JUND (right), which often binds as a heterodimer with FOS or other co-factors. Colored bars indicate the subset of regions with detectable footprints and how this percent varies depending on the chosen upper bound threshold.

**Figure S7G) Optimal performance scores for CTCF and JUND.** As described in Figure 7A, but here the cell color and annotation reflect the F1 score of CTCF and JUND specifically.

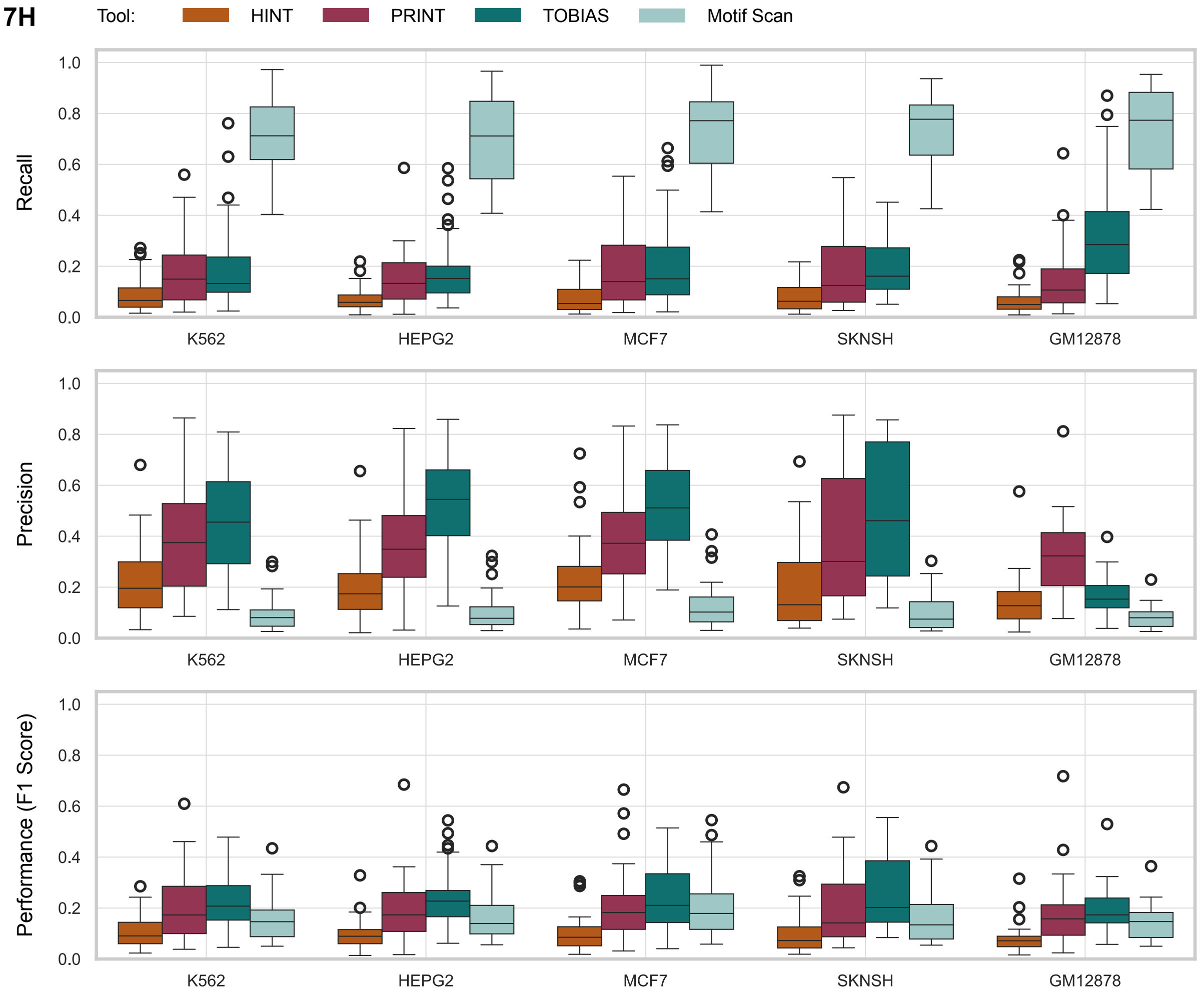

**Figure S7H) Expanded view of Figure 7C including all cell lines and performance metrics.**

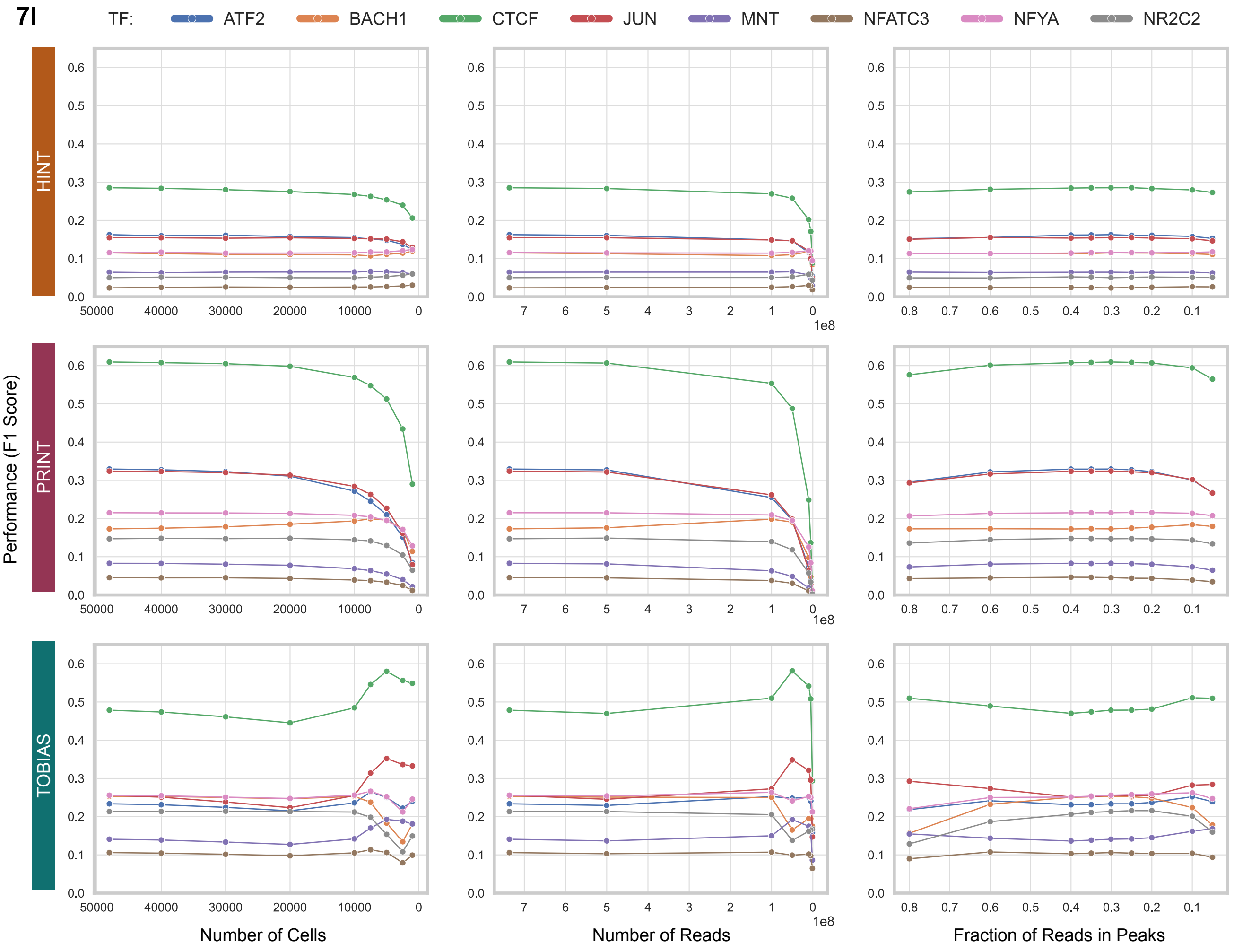

**Figure S7I) TF performance plateaus with data reduction and shows distinct maxima.** Line plots show F1 score trajectories across downsampling conditions using annotated ChIP regions as ground truth. Columns indicate the three downsampling types; rows represent different tools. Results are shown for the K562 cell line; eight TFs were randomly selected for illustration.

**7J**

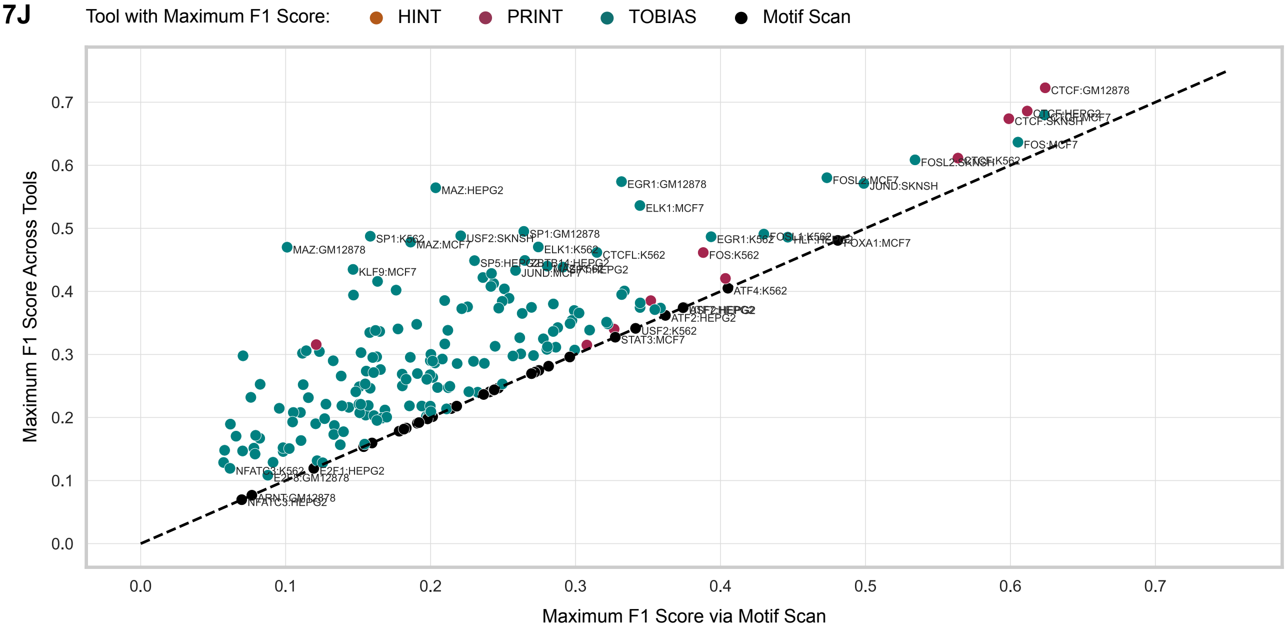

**Figure S7J. Motif scanning shows reasonable accuracy when evaluated against ChIP-seq.** Scatterplot showing the maximum F1 score achievable by motif scanning (x-axis) versus the highest F1 score obtained by any tested tool (y-axis) across all TF–cell line pairs. Each point is colored according to the tool that achieved the best score for that pair. The diagonal (x = y) line is shown as a dashed black line. Top- and bottom-performing TF–cell line pairs are labeled.

**8A**

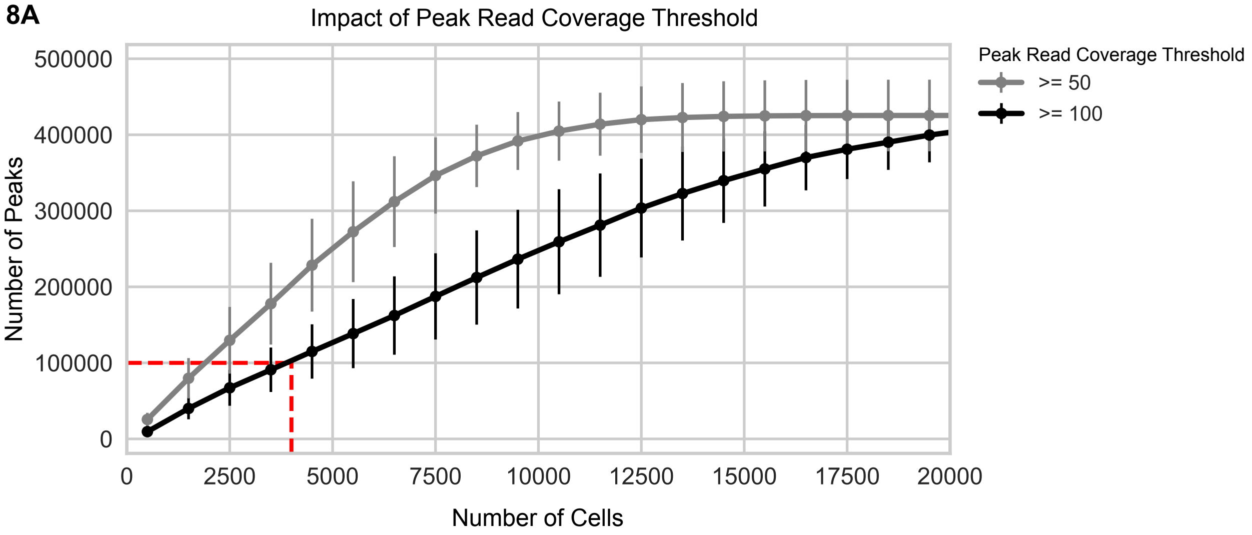

**Figure S8A) The impact of read-coverage thresholds on the number of retained peak regions.** The number of peaks retained after applying a minimum read-coverage threshold of 100 (black) or 50 (grey), shown as a function of the number of cells randomly sampled from each cell line. The mean and standard deviation across cell lines is shown.

**8B**

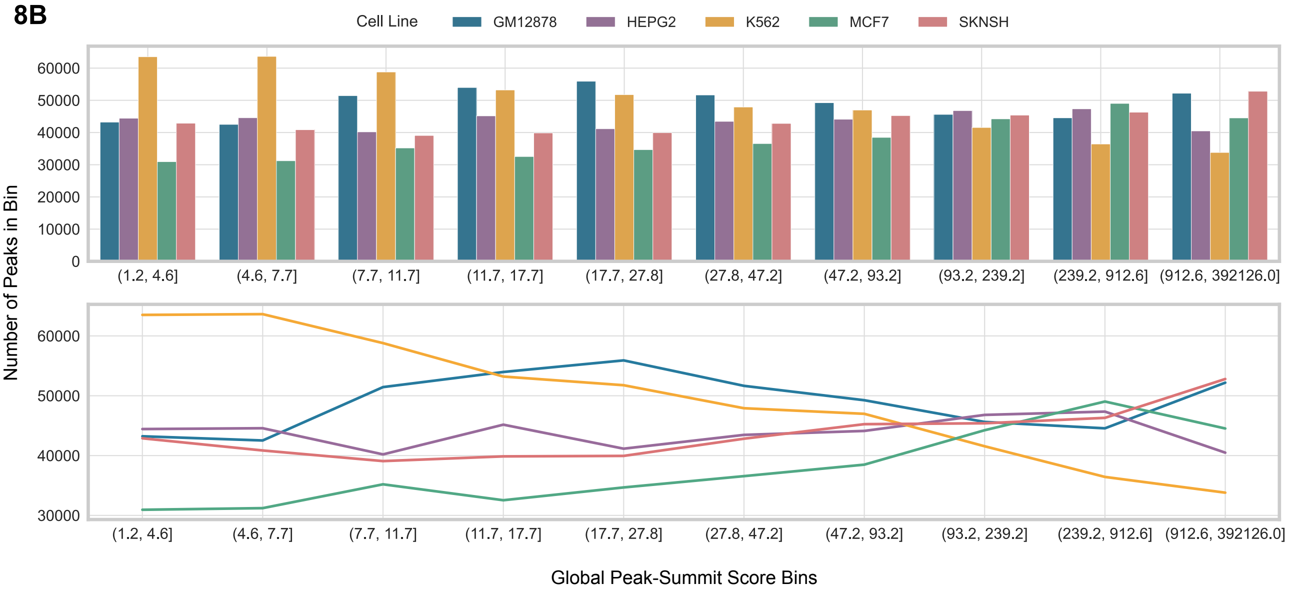

**Figure S8B) Cell lines produce input peak files with variable summit score distributions.** The top row displays the number of peaks (y axis) falling within each summit score bin (x axis) for each cell line (colors). The bottom row presents the same data as connected points to better highlight trajectory differences across cell lines, such as K562 and MCF-7. Anecdotally, applying a MACS2 summit score filter of 20 helps reduce footprinting noise from low read support regions.

**8C**

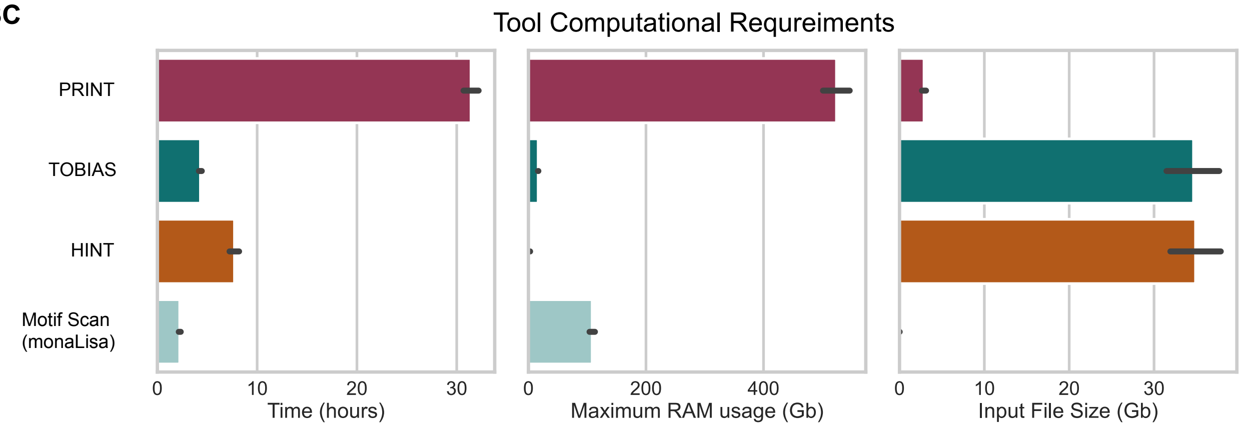

**Figure S8C) Computational requirements across tools.** Bar charts showing the average runtime, memory usage, and input file size requirements for each tool. Averages are calculated across all downsampling conditions and cell lines, with standard deviations indicated by error bars.
